# Using genetic variation to disentangle the complex relationship between food intake and health outcomes

**DOI:** 10.1101/829952

**Authors:** Nicola Pirastu, Ciara McDonnell, Eryk J. Grzeszkowiak, Ninon Mounier, Fumiaki Imamura, Jordi Merino, Felix R. Day, Jie Zheng, Nele Taba, Maria Pina Concas, Linda Repetto, Katherine A. Kentistou, Antonietta Robino, Tõnu Esko, Peter K. Joshi, Krista Fischer, Ken K. Ong, Tom R. Gaunt, Zoltan Kutalik, John R. B. Perry, James F. Wilson

**Affiliations:** Centre for Global Health Research, Usher Institute, University of Edinburgh, Teviot Place, Edinburgh, EH8 9AG, Scotland; Centre for Cardiovascular Sciences, Queen’s Medical Research Institute, University of Edinburgh, Royal Infirmary of Edinburgh, Little France Crescent, Edinburgh EH16 4TJ, Scotland; Center for Primary Care and Public Health, University of Lausanne, Lausanne, Switzerland; Swiss Institute of Bioinformatics, Lausanne, Switzerland; MRC Epidemiology Unit, Institute of Metabolic Science, Cambridge Biomedical Campus, University of Cambridge School of Clinical Medicine, Box 285, Cambridge, CB2 0QQ, UK; Diabetes Unit and Center for Genomic Medicine, Massachusetts General Hospital, Boston, MA, USA; Program in Medical and Population Genetics, Broad Institute, Cambridge, MA, USA; Department of Medicine, Harvard Medical School, Boston, MA, USA; MRC Integrative Epidemiology Unit, Bristol Medical School, Bristol, UK; Estonian Genome Center, Institute of Genomics, University of Tartu, Tartu, Riia 23b, 51010, Estonia; Institute for Maternal and Child Health - IRCCS “Burlo Garofolo”, Trieste, Italy; Department of Medical Epidemiology and Biostatistics, Karolinska Institutet, Nobels väg 12A, SE-171 77 Stockholm, Sweden; MRC Human Genetics Unit, Institute of Genetic and Molecular Medicine, University of Edinburgh, Western General Hospital, Crewe Road, Edinburgh, EH4 2XU, Scotland; Institute of Molecular and Cell Biology, University of Tartu, Tartu, Riia 23, 51010, Estonia

**Author notes:** Correspondence should be addressed to Nicola Pirastu. Authors contributed equally to this work (Ciara McDonnell, Eryk J. Grzeszkowiak).

## Abstract

Despite food choices being one of the most important factors influencing health, efforts to identify individual food groups and dietary patterns that cause disease have been challenging, with traditional nutritional epidemiological approaches plagued by biases and confounding. After identifying 302 individual genetic determinants of dietary intake in 445,779 individuals in the UK Biobank study, we develop a statistical genetics framework that enables us, to directly assess the impact of food choices on health outcomes. We show that the biases which affect observational studies extend also to GWAS, genetic correlations and causal inference through genetics, which can be corrected by applying our methods. Finally, by applying Mendelian Randomization approaches to the corrected results we identify some of the first robust causal associations between eating patterns and cancer, heart disease, obesity, and several other health related risk factors, distinguishing between the effects of specific foods or dietary patterns.

## Introduction

Given their profound impact on human well-being, diet is one of the most studied human behaviours. Quality, quantity, and patterns of consumed foods are associated with a wide range of medical conditions such as metabolic, inflammatory, or mental health diseases^1^. However, despite the growing number of studies reporting associations between diet and health outcomes, it has been challenging to establish causal relationships due methodological limitations such as measurement error, confounding, and reverse causation. To date, several methods have been devised to try to account for intrinsic limitations in nutritional studies such as calibration of food records^2^ or the implementation of domiciled feeding studies (ie. the PREDICT study^3^) in which participants are instructed to eat only the food provided by the study. Although these methods have helped in addressing some the limitations related to food consumption measurement, problems still remain especially when it comes to measure the effects of food on health over a long period of time.

In this context genetics may represent an alternative approach through the use of Mendellian Randomization. Mendelian Randomization (MR) is a methodological approach in which genetic variants associated with a phenotype of interest are used as instrumental variables to measure the “life-long effect of an exposure” to an outcome.^4^ To date, several MR studies have been designed to investigate the associations between the consumption of single food groups, such as alcoholic beverages^5^, coffee^6^, milk^7–9^ and specific health outcomes, but a systematic study investigating the overall role of diet is missing. In addition, previous MR studies have not accounted for the fact that genetic variants associated with reported dietary intake may be primarily associated with other risk factors or social determinants of health which may confound the causal estimates if used. In addition, previous studies on single food groups have not accounted for inter-relationships between different foods thus limiting the interpretability of the findings.

Given the complex number of factors that are driving the association between diet and health outcomes, the present study was designed to initially identify the genetic variants associated with reported food consumption, and then to leverage a causal inference statistical framework to systematically investigate the causal effects of dietary factors on health outcomes, while accounting for the effects that health determinants have on habitual dietary intake reporting.

## Methods

### Study population and genome-wide association for dietary intake

The UK Biobank^10^ is a large population-based cohort including 500 000 adults aged between 40 and 69 years at baseline across 22 assessments centers in the United Kingdom. Data were collected based on clinical examinations, assays of biological samples, detailed information on self-reported health characteristics, and genome-wide genotyping. Dietary intake in UK Biobank was assessed using a food frequency questionnaire which included questions about the frequency of consumption specific foods and beverages over the past year. The number of samples used for each trait can be found in table S1 while a detailed description of the phenotypes, can be found in the in the supplementary methods 1.2 and table S2.

We used the BOLT-LMM software^11^ to assess the association between the genetic variants across the human genome and 29 food phenotypes. Analyses were conducted on genetic data release version 3 imputed to the HRC panel^12^, as provided by the UK Biobank (http://www.ukbiobank.ac.uk/wp-content/uploads/2014/04/UKBiobank_genotyping_QC_documentation-web.pdf). Population stratification was assessed using LD-score regression as implemented in LD Hub^13,14^ using the LD scores provided with the software. Table S15 reports for each food trait the LD regression intercept and heritability estimation using ldsc. Cluster analysis conducted on the foods identified 5 main groups of traits (see additional online methods paragraph 1.8 and 2.2 for details of group definition) and we thus set the genome-wide significance threshold at 1×10^−8^. Work within was conducted under UKB application 19655. Participants enrolled in UK Biobank have signed consent forms. Replication analyses for identified signals associated with food phenotypes were conducted independently by using genetic and dietary data from the EPIC-Norfolk Study^15^ and the Fenland Study^16^. Details additional online methods 1.4.

### Investigating the effect of health outcomes on reported food intake using MR

Univariable MR analyses were initially conducted to measure the causal effect of health outcomes on food consumption using the TwoSampleMR^17^ R package. Exposures of interest were selected amongst those for which nutritional advice is given and included body mass index (BMI), low density lipoprotein cholesterol (LDLc), high density lipoprotein cholesterol (HDLc), Total cholesterol, Triglycerides, Diastolic and Systolic blood pressure, Type 2 diabetes, and coronary artery disease. In addition, we included educational attainment as a proxy of socio-economic status which is likely to affect food consumption. The full list of studies from which the summary statistics were derived is detailed in Table S6. For each exposure we selected all SNPs with *p*<5 x 10^−8^ and r^2^<0.001 to be used as instruments in the MR analysis. After performing stepwise heterogeneity pruning we performed MR analysis using the inverse variance method^18^. We then tested if the intercept from the MR-Egger^19^ regression was different from zero (*p*<0.05). If this was the case, MR-Egger was used for the analysis instead.

### Measuring the direct effects of food types on health outcomes and identifying genetic variants with predominantly direct-effects

One of the most important assumptions in MR is that the effect of the instrument on the outcome must be mediated only through the exposure of interest (sometimes referred as exclusion restriction criteria)^20^. In this light the instruments whose effect on food is mediated through the health outcomes or through educational attainment may violate this assumption acting as confounders in the relationship between the exposure and the outcome. Moreover if the mediating trait is acting on the reporting of food consumption and not food consumption itself it would mean that the genetic variant is not truly associated to food consumption and it would thus not be a valid instrument. It is thus important to estimate the direct effect(i.e., the effect that acts directly on food intake rather than is mediated through other factors see Figure 1) the SNPs are exerting on actual food consumption in order to properly select the genetic variants to be used as instrumental variables.

**Fig 1.**
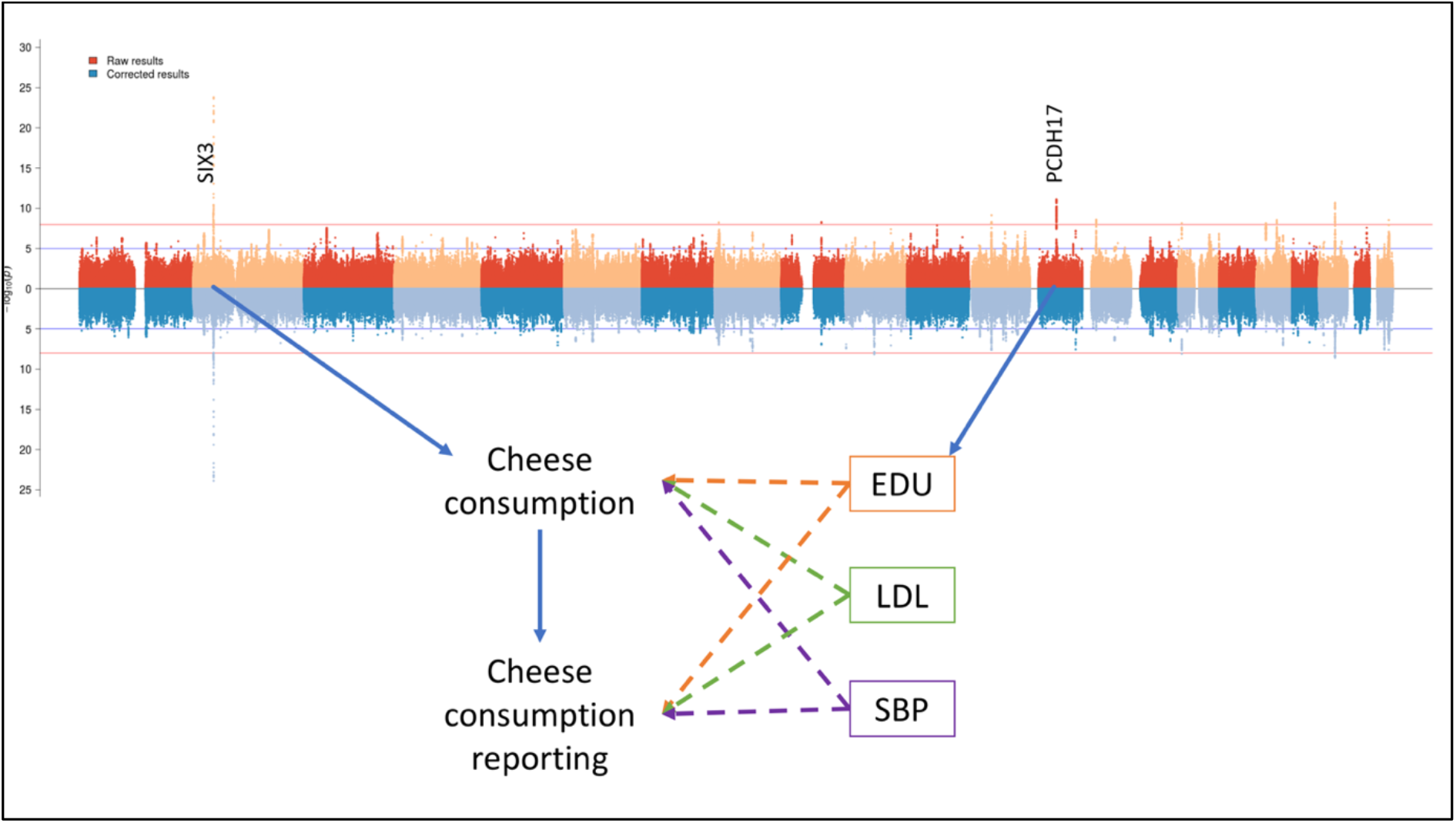
Direct and indirect SNP effects. The plot shows the causal path of exemplar genes identified for cheese consumption. In the multivariable MR model cheese consumption is causally influenced by educational attainment (EDU), low density lipoprotein cholesterol levels (LDL) and systolic blood pressure (SBP). The effect of PDCH17 and is mediated through educational attainment, while SIX3 has a direct effect on cheese consumption. The mediated effects cannot be used reliably as MR instruments as they could be affecting either consumption or its reporting. Moreover, they could act as confounders in the MR analysis and thus they need to be identified.

To this end we use a modified version of the method implemented in bGWAS^21^. This method consists of a first step were the phenotype of interest (i.e., food consumption) is used as outcome in multivariable MR. Next, exposures of interest are selected using a forward step wise regression selection algorithm where each exposure is added until their p-value is less than 0.05. The method provides a corrected estimate for each genetic variant of its effect on the outcome trait once all mediated effects are removed. Further details can be found in supplementary methods 1.6. In order to identify genetic variants with only a direct effect on the phenotype of interest we defined the corrected to uncorrected ratio (CUR) as the ratio between the corrected and the uncorrected effects (see additional methods 1.7 for a detailed explanation).

The threshold to define genetic variants with non-mediated effects (CUR=1±0.05) is based on simulations provided in the supplementary note 2.1 and on the genetic variants with known biological function (ie. bitter receptors). We defined as “non-mediated” those SNPs whose CUR fell within the defined ranges while “uncertain” the others. We applied bGWAS to all 29 food phenotypes. As potential mediators, we used the same cardiometabolic phenotypes as before except total cholesterol to avoid collinearity issues with LDL and HDL cholesterol, and we added summary statistics from Crohn’s disease and ulcerative colitis as they are likely to affect dietary patterns. A Detailed discussion of this approach can be found in supplementary methods 1.6.

### Genome-wide genetic correlations between corrected dietary intake and health outcomes

We used LD-score regression implemented in LD Hub^13,14^ to estimate genome-wide genetic correlations between dietary intake phenotypes and 844 health outcomes and intermediary phenotypes. Genetic correlations were estimated both with the corrected and uncorrected GWAS summary statistics using the bivariate LD-score regression model. Stratified LD-score regression^22^ analyses were implemented using ldsc and the annotation files available on the ldsc website.

### Definition of food group variables

In order to define measures of dietary patterns we first performed cluster analysis of the 29 food items applying iCLUST^23^ to the corrected genetic correlation matrix between the different foods. iCLUST clusters items in different groups based on a hierarchical structure (Details additional methods 1.8). Figure 2 shows the resulting dendrogram and its comparison with the genetic correlation matrix.

**Fig 2.**
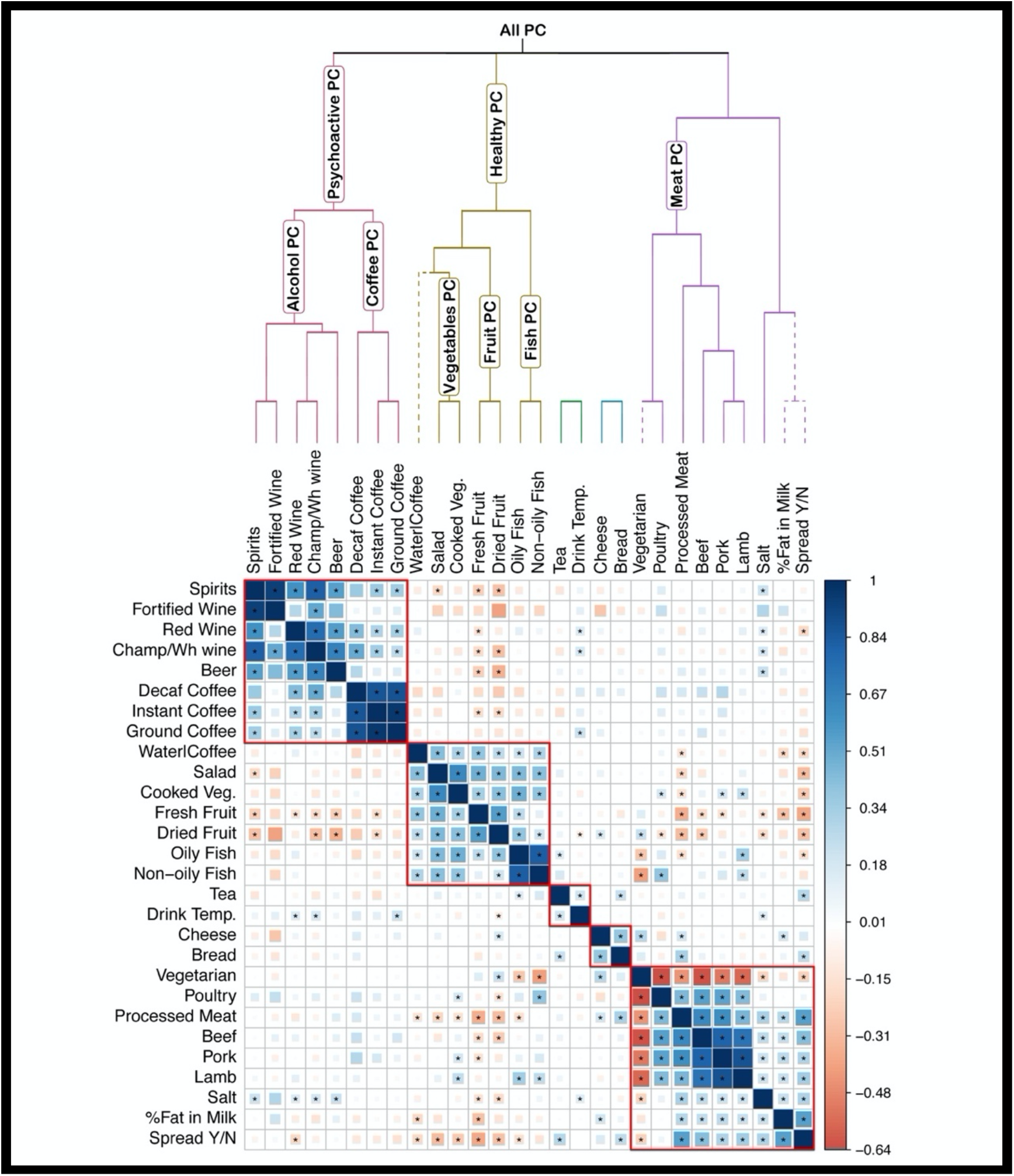
Clustering of the food traits and definition of measures of dietary patterns. The plot reports the genetic correlation plot amongst the food traits after applying the correction. The stars report the Bonferroni-corrected significant correlations. The dendrogram and the boxes represent the clustering according to the ICLUST algorithm. The labels on the dendrogram branches show the traits used to define each measure of dietary pattern. The dashed line represents the traits excluded from the estimation of the dietary patterns traits. The “Vegetarian” trait was excluded from the “Meat PC” trait but was included in the overall dietary pattern measure (All PC).

We then defined based on the resulting structure several measures of dietary pattern at different levels of the dendrogram as shown in Figure 2. For each measure we performed principal component analysis of the items which participated to each group. The rotation matrix was derived from the eigen decomposition of the correlation matrix of the foods in the PC trait of interest. For example for the Coffee PC measure we performed principal component analysis of “Ground Coffee”, “Instant Coffee” and “Decaf Coffee”. Once the rotation matrix was estimated for each SNP its effect on the new measure was estimated as the linear combination of the effect on each food trait using as weights the loadings on each PC. A correlation plot of the loadings of each item onto the PC traits can be found in figure S3.

### MR analyses to assess causal relationships between food intake and health outcomes

MR analyses were conducted to estimate the effects of the food phenotypes on 79 health related phenotypes (see table S17 for details) available in MR-base.^17^ Genetic instruments for each exposure of interest included independent genetic variants (p<5×10^−8^ and pruning for LD (r^2^<0.001)). For dietary patterns exposures SNPs were selected as outlined in additional methods 1.12. For the main analysis we restricted the genetic instruments to those that only had evidence of a direct effect (i.e., not affecting the main exposure through a different pathway; CUR 1±0.05). Discussion of the relationship with other methods can be found in supplementary note 2.7. Weights for the genetic instruments were based on the uncorrected effects. To verify the effects of using only direct effect only SNPs on MR, all the analyses were also conducted without applying the CUR filtering.

After selecting the genetic instruments, exposure and outcome data were harmonised. The MR estimates were tested for heterogeneity and outliers were removed using the MR-Radial method.^24^ MR analyses were based on the inverse variance weighted method, which estimates the causal effect of an exposure on an outcome by combining ratio estimates using each variant. A random effect model was used if significant heterogeneity between the different estimates was detected. We then tested for the presence of directional pleiotropy using the intercept from the MR-Egger regression. MR median and MR-Raps were used as sensitivity analyses. All results have been made available through an online app (https://npirastu.shinyapps.io/Food_MR/) and can be found in additional table S18.

### Patient and public involvement

This research did not involve patients or the public as it uses data from the UK Biobank study that were previously obtained from a cohort of people who had already been recruited. As such, no patients or member of the public were involved in the design or implementation of this study or the research questions addressed.

## Results

### Genetic variants associated with food intake

In a GWAS of 29 food phenotypes we identified 414 genetic associations in 260 independent loci (Fig 3 and additional table S4) at Bonferroni corrected level of significance (P< 1×10^−8^).

**Fig 3.**
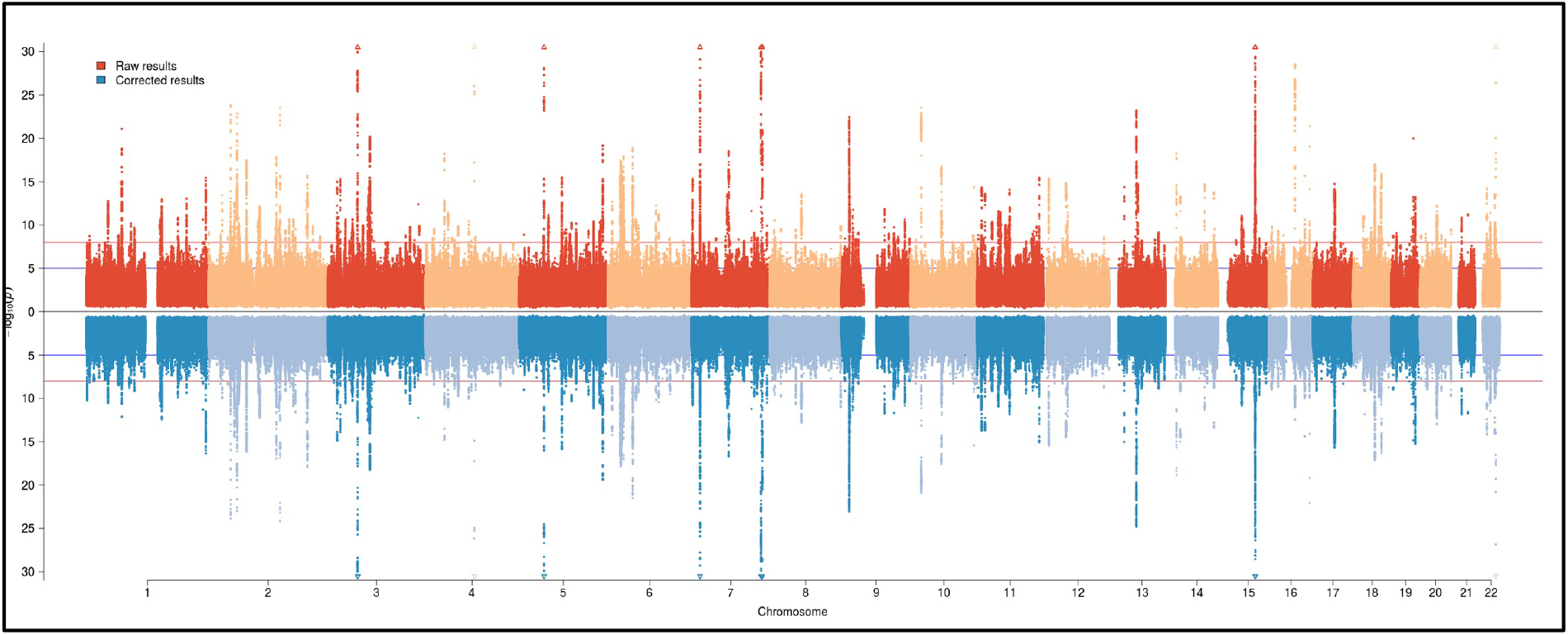
302 independent genomic loci associate with food choices. Results for both univariate (260 loci) and multivariate (additional 42 loci see paragraph S2.3) analyses are included. For each SNP the lowest p-value for all traits was plotted. The upper panel represents the unadjusted GWAS associations while the lower panel represents the association with food choices, after adjustment for mediating traits, such as health status.

Replication was sought in two additional UK-based cohorts including up to 32,779 participants. Despite relatively limited power in replication cohorts, concordant direction of effect was observed for 82% of the signals (p=7.82×10^−35^, Binomial test; Table S5), and nominal significance was achieved by 32% of the signals (p=9.47×10^−54^). Gene prioritization is described in supplementary methods 1.10 while biological annotation, network analysis and tissue enrichment analysis are discussed in additional paragraphs 1.11, 2.4 and 2.5. Several of the identified loci have been previously associated with BMI. However, contrary to our expectations, the BMI-raising allele was consistently associated with lower reported consumption of energy-dense foods such as meat or fat, and higher reported intake of low-calorie foods.

### Genetic variants associated with food intake are strongly influenced by other phenotypes

In univariable MR we identified 81 instances in which health-related traits significantly influencing food intake (Fig. 4 additional table S7). In particular BMI and Educational attainment influenced more than 50% of the food traits. Similar effects extend to a broad range of traits, for example LDL and triglycerides influenced 15 and 18 traits respectively. Higher genetically-determined CAD associates with higher consumption of fish and red wine, and lower consumption of whole milk, salt and lamb. These findings suggest that some of the signals identified in GWAS for reported food phenotypes are not directly associated with food intake but are mediated through a wide range of potential confounders.

**Fig 4.**
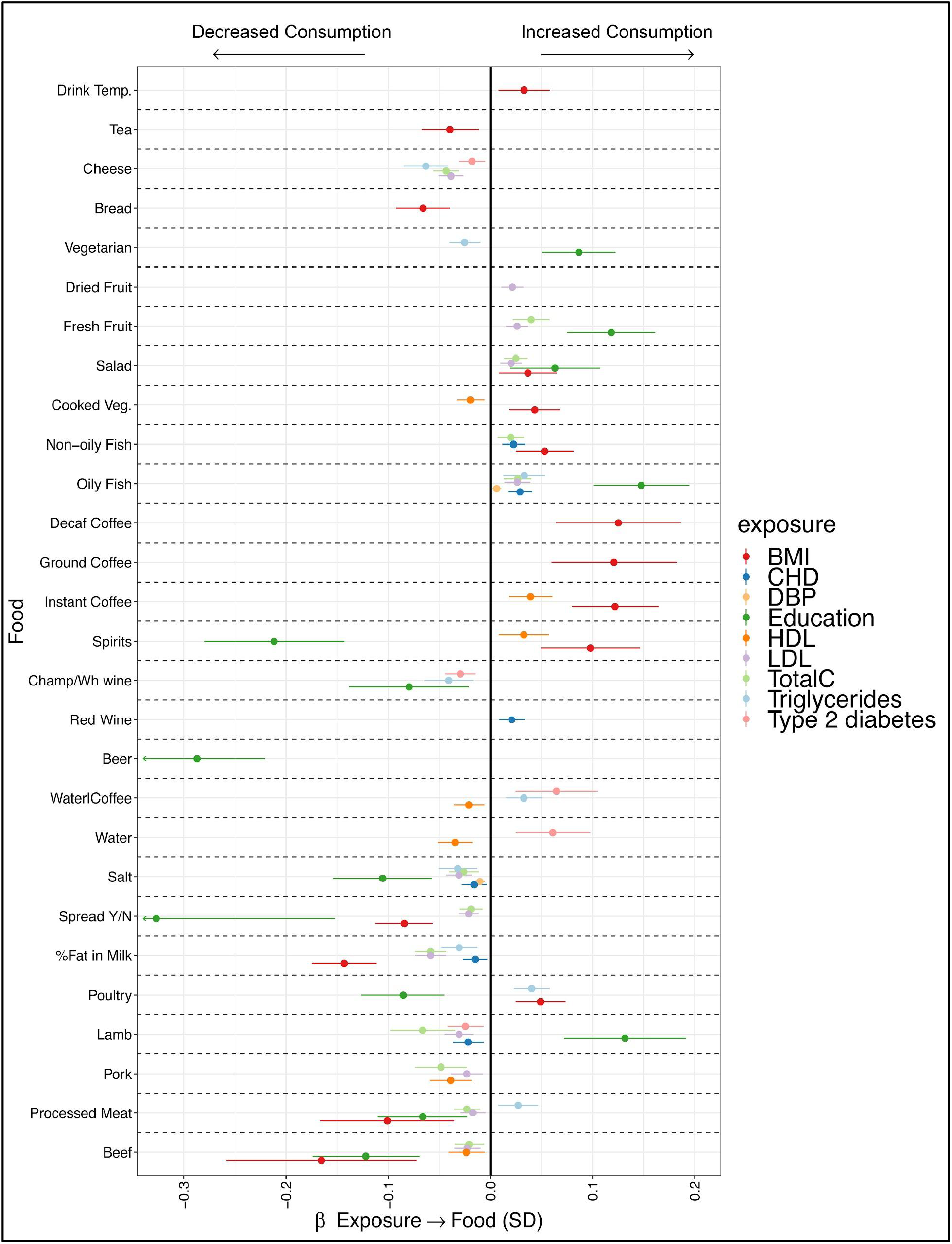
Health status influences reported food choices. The plot reports only the univariable MR results which were significant at FDR<0.05. For each food outcome the effect estimate (β) is reported in standard deviations of the exposure trait, together with 95% confidence intervals. Each colour represents a different exposure. BMI, body mass index; CHD, coronary heart disease; DBP, diastolic blood pressure; HDL, high density lipoprotein cholesterol; LDL, low density lipoprotein cholesterol; TotalC, total cholesterol. Champ/Wh wine, champagne, white wine. Temp, temperature.

The Multivariable MR confirmed the univariable MR results (Supplementary Fig S4 panel A and Supplementary Table S8). The percentage of genetic variance for the reported food phenotypes explained by health determinants ranged from 42% for cheese to ∼0% for fortified wine and white wine/champagne (Supplementary Fig S3 panel B and Supplementary Table S16). We systematically compared the estimated effect sizes of each genetic variants influencing food consumption before and after correcting for the effect of health determinants and showed that in many loci the variant initially identified for food phenotypes changed dramatically after taking into account the effect of health factors (Fig. 3, see Supplementary file 1 for trait-specific plots). For example, the effect size of the lead *FTO* variant (rs55872725, *p*=2×10^−29^) on milk fat percentage chosen decreased three-fold after accounting for the mediated effects. To further explore the magnitude of this indirect effect on food intake phenotypes, we compared the correlation patterns between the 29 food phenotypes and 832 phenotypes present in the LD hub^14^ database identifying great differences. For example, low fat milk intake was correlated with a beneficial effect on body fat percentage (r_G_ = −0.43) but this association diminished to near zero (r_G_ = −0.04) after accounting for indirect effects (Supplementary Data 2.2 and additional table S10). The effects of the correction procedure on the genetic correlation amongst the traits and with the 844 health traits are discussed in supplementary note 2.2 while full results can be found at in table S9 and browsed at https://npirastu.shinyapps.io/rg_plotter_2/. These findings highlight the relevance of biases and confounding in genetic correlation studies, and provide the framework to study complex physiological relationships.

### Causal inference analyses for diet phenotypes and health outcomes

A total of 230 out of 414 genetic variants initially associated with food phenotypes (corresponding to 169/260 loci) were categorized as “non-mediated” associations (Table S3). The balance of uncertain to non-mediated genetic associations varied by food group, ranging from none uncertain for tea, spirits and processed meat, to all uncertain for percentage fat in milk and adding spread to bread (Table S3).

In two-sample MR analyses we found 141 significant associations between food phenotypes and health outcomes after multiple test correction (pFDR < 0.05, Table S18).

Of these 89 showed no sign of heterogeneity amongst the estimates (heterogeneity test p >0.05). Figure 5 reports full results for all significant food exposure trait outcome pairs.

**Fig 5.**
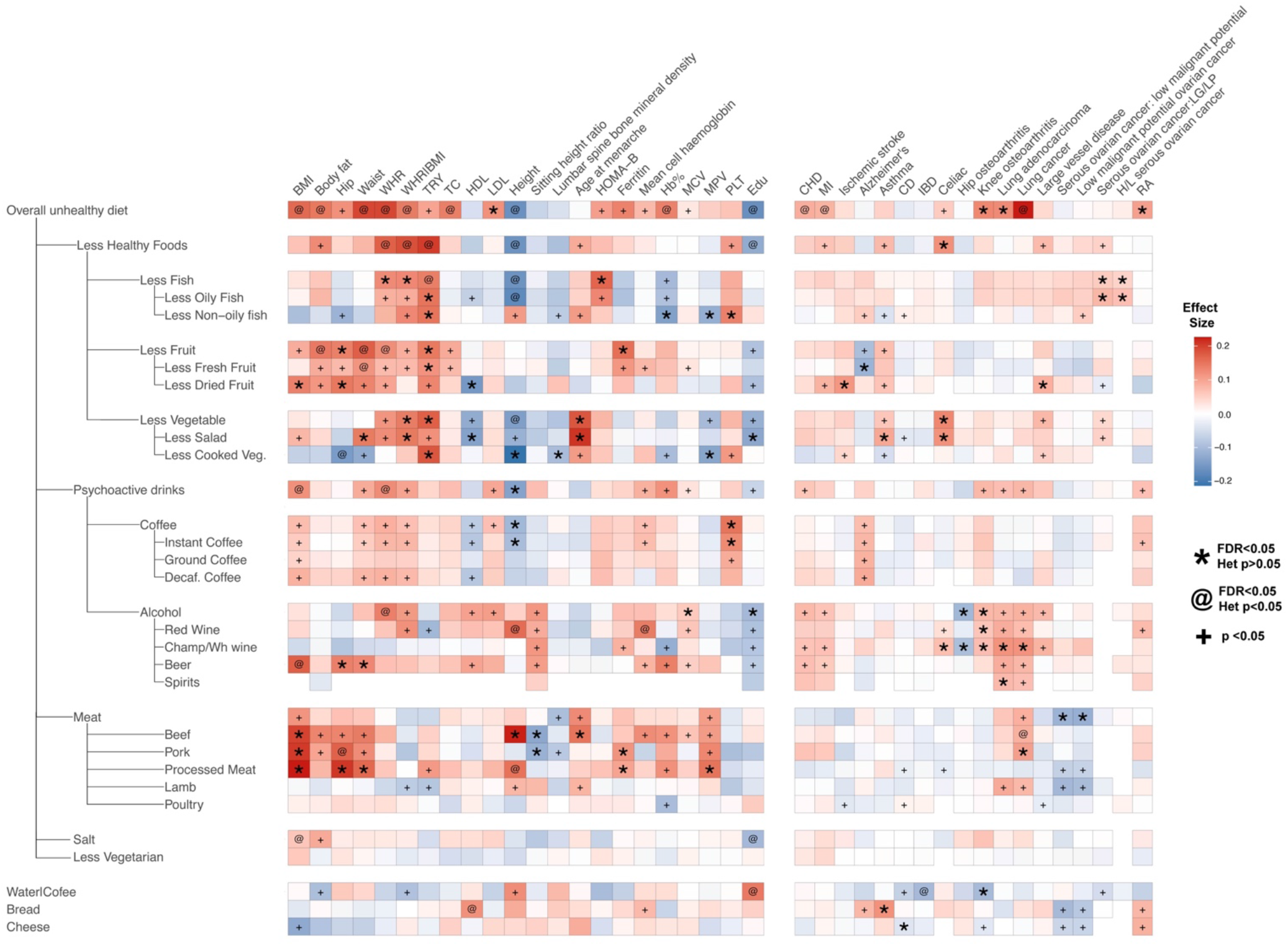
Significant effects of food choice on disease related traits. The heatmap reports the results for all significant food trait exposure trait outcome. Only dietary pattern exposures summarising the overall group consumption (PC1) have been reported. All exposures have been aligned to have a positive loading onto the “overall unhealthy diet” measure. Significant food/trait association are indicated with * if they show no sign of heterogeneity while @ if they show significant heterogeneity. To facilitate meaningful visualisation and maximise the appearance of signal rather than noise, we applied a shrinkage method - imposing a bayesian prior assumption on the distribution of beta (mean 0, SD 0.1), and conjugating that with the likelihood of our results and then taking mean beta from the resulting distribution, thus shrinking estimates with larger SEs more towards 0. Abbreviations: BMI Body Mass Index, WHR Waist to Hip Ratio, TRY tryglicerides, TC total cholesterol, HDL HDL cholesterol, LDL LDL cholesterol, Hb% Haemoglobin percentage, MCV Mean Corpuscolar Volume, MPV Mean Platelet Volume, PLT Platelet count, Edu Educational attainment, CHD Coronary Heart Disease, MI Myocardial Infarction, CD Chron’s Disease, IBD Inflammatory Bowel Disease, Serous ovarian cancer:LG/LP low grade low potential. H/L serous ovarian cancer High and Low grade serous ovarian cancer, RA Rheumatoid Arthritis.

Overall we found evidence supporting the beneficial effect of a healthy diet on health outcomes. For example, for obesity/adiposity outcomes, genetically-determined unhealthy diet leads to very similar effects across, increasing obesity measurements. For lipid-related outcomes, the overall unhealthy diet is associated with higher levels of LDLc with no significant heterogeneity, but no association with any of the other dietary traits. The overall unhealthy diet was also strongly associated with Lung adenocarcinoma (OR 1.4xSD CI 1.2-1.9) which seemed to be driven mostly by alcoholic beverages.

We identified 51 instances in which we would have not detected a significant result without filtering out the non-direct effect instruments such as the effect of increased fruit consumption on triglycerides levels (estimated uncorrected effect= −0.03 (SE=0.05) vs. estimated corrected effect = −0.17 (SE=0.05)) or the effect of increased beef consumption on height (uncorrected effect = −0.02 (−0.17, 0.13) vs corrected effect = −0.52 (0.29, 0.74). In addition, we found 124 food/trait relationships which were not significant after applying CUR filtering, showing that either confounding effects or reduced power explain the lack of association (see additional note 2.6). For example, red wine consumption was initially associated with increased BMI (uncorrected effect =0.22 (SE 0.05)) and waist circumference (uncorrected beta= 0.26 (SE 0.07), but after correcting for CAD liability, both effects disappeared (corrected effect for BMI 0.05 (SE 0.06), corrected effect for WC 0.005 (0.08)). On the flip side, we showed that the effect of red wine on mean corpuscular volume remains substantially unchanged when applying the filtering approach (beta 0.07 (SE 0.02) uncorrected and 0.065 (SE 0.02) corrected), suggesting that our approach could precisely identify relevant biological relationships.

A full description of our findings are found in table S18 and have been made available through an online app (https://npirastu.shinyapps.io/Food_MR/).

## Discussion

In this study we have provided quantitative data about the complex interplay between diet and health outcomes showing that the causal path from food intake to adverse health outcomes is not unidirectional and may be influenced by reverse causation and confounding even when MR is used. We showed that genetic correlations and causal inference can be improved by leveraging statistical approaches that take into account this mediated effects and identify genetic variants that have a only non-mediated effects on the exposure of interest. This information allowed us to perform causal inference analyses that helped identifying more reliable potential causal effects of food on health outcomes.

## Results in context

Previous MR studies have mainly focused on specific food groups such as coffee, alcohol and milk consumption while none has comprehensively investigated the role of different food groups on health outcomes. Our results support previous observations such as the effect of alcohol consumption on coronary artery diseases reported in previous MR studies. In addition, we were able to confirm similar previous results detecting no evidence of an effect on IBD and CD^25^, ovarian cancer^26^ or rheumatoid arthritis^27^.

Findings from this study also suggest that the same biases that affect measures of food consumption such as reporting bias, confounding and reverse causation are reflected also in studies focusing on genetic associations. We have shown that these issues extend beyond obesity and socio-economic status including a broader range of intermediate factors. For example blood LDL and triglycerides concentration influence a wide variety of food traits thus being important factors to be considered as potential sources of bias, yet to our knowledge this is the first time this has been reported. For our analyses we have used UK biobank in which participants were aged between 40 and 60 at the time of the questionnaire, it is likely that a younger cohort will suffer less from some of these (ie. LDL cholesterol or blood pressure) as it is unlikely that they will display pathological level of these traits.

Our results are in contradiction to some previous studies in which no evidence of reverse causation influencing genetic susceptibility for dietary patterns was reported.^28,29^ We believe that this difference is due to our novel approach, which is not based on using the potential mediators as covariates, but rather exploits MR, which should be able to distinguish the forward and reverse effects when the causal relationship is bidirectional. We have thus shown that it is possible, through the use of available data and methods, to disentangle these different colliding effects and to select the instrumental variables which show a non-mediated effect, thus enabling the use of MR for the assessment of causal relationships between food and health.

Many studies have looked at the relationship between nutritional composition and health outcomes. One of the most salient examples is the relationship between saturated fat intake and cardiovascular disease and all-cause mortality, in which recent studies suggest that food sources of saturated fatty acids are more important than saturated fat content per se^[Citation error]^. Our study provide a new angle on the importance of food sources by providing evidence that foods with similar nutrient profile, for example cheese and meat, which are both relatively high in saturated fat and protein, have opposite effects on some metabolic risk factors such as BMI (Figure S24 A) but there is no difference in other phenotypes such as blood lipids. A similar conclusion can be drawn if we look at the foods which have the greatest effect on triglycerides, fruit, vegetables and fish; all with very similar lowering effects (Figure S24 B), which have relatively different macronutrient compositions. While the findings require further investigations in mechanisms and related behaviours, our genetic evidence lends support for the importance of studying foods in their complexity and not as a mere mixture of nutrients. This approach, in fact, does not consider that the sources of the nutrients are not equal due to the food matrix, the different preparations and that foods are seldom consumed by themselves but in patterns which are likely to modify the effects on health.

Our findings illustrate that the effect of diet on health outcomes is complex, and components of specific food groups have a differential association with health. In this case, although fish and fruit and vegetables have a very different macronutrient composition it was impossible to separate their effect on triglyceride concentrations. This suggests that at least in this case the macronutrient composition is not as important as the an overall tendency to eat certain foods and it highlights the importance of always including the assessment of dietary patterns before claiming health effects of single foods or nutrients.

Some of the effects we have identified are more complex to explain and will need different sources of evidence to be understood. For example we have found that the overall unhealthy diet is associated to a higher risk of both lung andenocarcinoma and lung cancer. When looking more closely to which of food explain this association the most we can see that Alcohol seems to be driving the overall effect. One possibility is that this relationship is confounded by smoking through a common tendency to addictive behaviours. However a recent GWAS on cigarette smoking in Japan Biobank^30^ reported a strong association between the ALDH2 gene and number of cigarettes per day smoked which has also been associated to differences in alcohol consumption^31^, suggesting a causal effect of increased alcohol consumption on increased smoking thus predisposing to lung cancer. Regardless of the interpretation this example shows how complex the interpretation of MR results are when behavioural traits are involved as they influence each other constantly creating a complex net of interrelationships. This also points to the need of extreme care when claiming beneficial health effects of food and multiple sources of evidence and approaches should always be used before translating these findings into public policies.

Our study has several potential limitations. First, the number of items available in the dietary questionnaire in the UK BioBank is limited, and therefore it limited our ability to capture overall diet or specific food groups not detailed. The inclusion of white and relatively healthy and educated participants from UK Biobank may have limited the generalisability of our findings. Estimated effect sizes could be inflated because of the underestimation of the SNP effects on the actual food trait consumption, rather than its self-report, if so, this will have inflated our estimates of the effects of food on health, due to the noise in the questionnaire responses, and warrants further statistical investigations. Even so, our method should not have falsely identified a causal effect or reversed its direction, but further studies are needed to assess the precise effect sizes.

In conclusion, our findings show that overall what is generally considered a healthy diet leads to many favourable health outcomes and to reducing a wide range of risk factors broadly agreeing with current guidelines aimed at reducing meat and alcohol consumption while increasing fruit vegetables and fish. We also show that some of these effects are mostly reconductable to specific food or group of foods which however are not characterized by common nutrient composition thus adding granularity to our knowledge on the effect of diet on health. This information can be useful to inform the design and implementation of future studies to reduce the burden of diet-related diseases.

## Author Contributions

NP,JFW,JRBP,ZK,EJG,FRD,KKO contributed to the study design.

JFW,TE,JRBP,AR,TG,FI,KKO,FRD contributed data.

NP,CMD,EJG,NM,FI,JZ,NT,KAK,MPC, performed the statistical analyses. NP, JFW,ZK,

JRBP, JM,TE, NT,KF,CMD,LR,EJG,FI,KKO,FRD contributed to the interpretation of the results. All authors contributed to writing and editing of the text.

## Acknowledgements

J.F.W. acknowledges support from the MRC Human Genetics Unit quinquennial programme grant “QTL in Health and Disease” (MC_UU_00007/10). This research has been conducted using the UK Biobank Resource under Application Number 19655. We would like to thank Professor George Davey Smith for the precious feedback, Erin MacDonald-Dunlop, and Pascale Lubbe for help with statistical analyses and Dr. Nana Matoba for providing the results from the smoking GWAS.

EGCUT was funded by Estonian Research Council Grant IUT20-60, PUT1660 (T.E.), PUT1665 (K.F.), the European Union through the European Regional Development Fund grant no. 2014-2020.4.01.15−0012 GENTRANSMED and 2014-2020.4.2.2, and Estonian and European Research Roadmap grant no.2014-2020.4.01.16−0125. The EPIC-Norfolk study (DOI 10.22025/2019.10.105.00004) has received funding from the Medical Research Council (MR/N003284/1, MC-PC_13048, and MC-UU_12015/1). The Fenland study (DOI: 10.1186/ISRCTN72077169) was funded by the Medical Research Council and the Wellcome Trust (Ref: 074548). J.P., K.O., F.I., and F.R.D. were funded by the UK Medical Research Council Epidemiology Unit core grant (MC-UU_12015/2, MC_UU_12015/5). T.R.G. receives funding from the UK Medical Research Council (MC_UU_00011/4). Z.K. received funding from the Swiss National Science Foundation (31003A_169929). We are grateful to all the participants who have been part of the project and to the many members of the study teams at the University of Cambridge who have enabled this research. J.M. was partially supported by a fellowship funded by the European Commission Horizon 2020 program (H2020-MSCA-IF-2015-703787) and by the National Institutes of Health grant P30 DK40561

## Data Availability

All GWAS results will be made available through GWAS catalog at the time of publication. All results from the MR analyses have been shared in the additional tables.

## References

1. Misra, A. & Khurana, L. Obesity and the Metabolic Syndrome in Developing Countries. The Journal of Clinical Endocrinology & Metabolism vol. 93 s9–s30 (2008).

2. Naska, A., Lagiou, A. & Lagiou, P. Dietary assessment methods in epidemiological research: current state of the art and future prospects. F1000Res. 6, 926 (2017).

3. Berry, S. E. et al. Human postprandial responses to food and potential for precision nutrition. Nat. Med. 26, 964–973 (2020).

4. Zheng, J. et al. Recent Developments in Mendelian Randomization Studies. Curr Epidemiol Rep 4, 330–345 (2017).

5. Millwood, I. Y. et al. Conventional and genetic evidence on alcohol and vascular disease aetiology: a prospective study of 500 000 men and women in China. Lancet 393, 1831– 1842 (2019).

6. Cornelis, M. C. & Munafo, M. R. Mendelian Randomization Studies of Coffee and Caffeine Consumption. Nutrients 10, (2018).

7. Bergholdt, H. K. M., Nordestgaard, B. G., Varbo, A. & Ellervik, C. Milk intake is not associated with ischaemic heart disease in observational or Mendelian randomization analyses in 98,529 Danish adults. Int. J. Epidemiol. 44, 587–603 (2015).

8. Vissers, L. E. T. et al. Dairy Product Intake and Risk of Type 2 Diabetes in EPIC-InterAct: A Mendelian Randomization Study. Diabetes Care 42, 568–575 (2019).

9. Hartwig, F. P., Horta, B. L., Smith, G. D., de Mola, C. L. & Victora, C. G. Association of lactase persistence genotype with milk consumption, obesity and blood pressure: a Mendelian randomization study in the 1982 Pelotas (Brazil) Birth Cohort, with a systematic review and meta-analysis. Int. J. Epidemiol. 45, 1573–1587 (2016).

10. Sudlow, C. et al. UK biobank: an open access resource for identifying the causes of a wide range of complex diseases of middle and old age. PLoS Med. 12, e1001779 (2015).

11. Loh, P.-R. et al. Efficient Bayesian mixed-model analysis increases association power in large cohorts. Nat. Genet. 47, 284–290 (2015).

12. McCarthy, S. et al. A reference panel of 64,976 haplotypes for genotype imputation. Nat. Genet. 48, 1279–1283 (2016).

13. Bulik-Sullivan, B. K. et al. LD Score regression distinguishes confounding from polygenicity in genome-wide association studies. Nat. Genet. 47, 291–295 (2015).

14. Zheng, J. et al. LD Hub: a centralized database and web interface to perform LD score regression that maximizes the potential of summary level GWAS data for SNP heritability and genetic correlation analysis. Bioinformatics vol. 33 272–279 (2017).

15. Day, N. et al. EPIC-Norfolk: study design and characteristics of the cohort. European Prospective Investigation of Cancer. Br. J. Cancer 80 Suppl 1, 95–103 (1999).

16. Lotta, L. A. et al. Integrative genomic analysis implicates limited peripheral adipose storage capacity in the pathogenesis of human insulin resistance. Nat. Genet. 49, 17–26 (2017).

17. Hemani, G. et al. The MR-Base platform supports systematic causal inference across the human phenome. Elife 7, (2018).

18. Burgess, S., Butterworth, A. & Thompson, S. G. Mendelian randomization analysis with multiple genetic variants using summarized data. Genet. Epidemiol. 37, 658–665 (2013).

19. Bowden, J., Davey Smith, G. & Burgess, S. Mendelian randomization with invalid instruments: effect estimation and bias detection through Egger regression. Int. J. Epidemiol. 44, 512–525 (2015).

20. Davies, N. M., Holmes, M. V. & Davey Smith, G. Reading Mendelian randomisation studies: a guide, glossary, and checklist for clinicians. BMJ 362, k601 (2018).

21. Mounier, N. & Kutalik, Z. bGWAS: an R package to perform Bayesian Genome Wide Association Studies. Bioinformatics (2020) doi:10.1093/bioinformatics/btaa549.

22. Finucane, H. K. et al. Partitioning heritability by functional category using GWAS summary statistics. doi:10.1101/014241.

23. Revelle, W. Hierarchical cluster analysis and the internal structure of tests. Multivariate Behav. Res. 14, 57–74 (1979).

24. Bowden, J. et al. Improving the visualization, interpretation and analysis of two-sample summary data Mendelian randomization via the Radial plot and Radial regression. Int. J. Epidemiol. 47, 2100 (2018).

25. Georgiou, A. N., Ntritsos, G., Papadimitriou, N., Dimou, N. & Evangelou, E. Cigarette smoking, coffee consumption, alcohol intake, and risk of crohn’s disease and ulcerative colitis: A Mendelian randomization study. Inflamm. Bowel Dis. (2020) doi:10.1093/ibd/izaa152.

26. Zhu, J., Jiang, X. & Niu, Z. Alcohol consumption and risk of breast and ovarian cancer: A Mendelian randomization study. Cancer Genet. 245, 35–41 (2020).

27. Jiang, X. & Alfredsson, L. Modifiable environmental exposure and risk of rheumatoid arthritis-current evidence from genetic studies. Arthritis Res. Ther. 22, 154 (2020).

28. Meddens, S. F. W., de Vlaming, R., Bowers, P. & Burik, C. A. P. Genomic analysis of diet composition finds novel loci and associations with health and lifestyle. bioRxiv (2018).

29. Cole, J. B., Florez, J. C. & Hirschhorn, J. N. Comprehensive genomic analysis of dietary habits in UK Biobank identifies hundreds of genetic loci and establishes causal relationships between educational attainment and healthy eating. doi:10.1101/662239.

30. Matoba, N. et al. GWAS of smoking behaviour in 165,436 Japanese people reveals seven new loci and shared genetic architecture. Nat. Hum. Behav. 3, 471–477 (2019).

31. Matoba, N. et al. GWAS of 165,084 Japanese individuals identified nine loci associated with dietary habits. Nat. Hum. Behav. 4, 308–316 (2020).

