## Additional materials for "Using genetic variation to disentangle the complex relationship between food intake and health outcomes"

### 1. Online Materials and Methods

#### 1.1 Discovery population.

Analyses were conducted on data collected in the UK Biobank project[1] under project 19655. All subjects gave written informed consent. UK Biobank has approval from the North West Multi-Centre Research Ethics Committee (MREC), In Scotland, UK Biobank has approval from the Community Health Index Advisory Group (CHIAG). We included only subjects who completed the food frequency questionnaire and were of European descent as defined in Uk biobank.

#### 1.2 Phenotype modelling

Quantitative food and drink intake phenotypes were all converted to weekly consumption, e.g. consuming 3 cups of tea a day was converted to 21 cups/week. Semi-quantitative descriptors: never, 2-4 times a week, 5-6 times a week, once or more daily were converted to 0, 3, 5.5 and 7 respectively. All “Prefer not to answer” and “Do not know” answers were excluded from the analysis.

All coffee traits were stratified by type (instant, ground, decaffeinated) to account for differences in consumption patterns such as cup size and caffeine concentration. Participants who did not specify the type of coffee they usually consume were excluded from the analysis.

Coffee consumption (any type of coffee, including unspecified type of coffee) was treated as a covariate for water consumption due to a very high negative phenotypic correlation between water and coffee consumption. Some semi-quantitative traits do not directly refer to the amount of food or drink consumed but to its type. Fat content in milk was calculated as the fat percentage and non-dairy types of milk (e.g. soy) were removed. Drink temperature (very hot, hot and warm) were converted to an arbitrary 3-unit scale (3, 2, 1). People who did not report consuming hot drinks were excluded from the analysis. Summary statistics for the traits are reported in Supplementary data 10.

All traits including the binary ones were treated as being quantitative. For binary traits this is the same as running a trend test. We did this because standard software cannot perform logistic regression on such a large sample in a timely fashion. Residuals for each trait using a linear model were first estimated in R using age and sex and then used as phenotype for the association analysis using BOLT-LMM[2]. Coffee and tea consumption were added as covariates for the analysis of water consumption as described before.

In order to verify the best function for trait normalisation in the association analysis and to ensure that most quantitative traits demonstrated a right-tailed distribution, we applied both log10(x+1) and sqrt(x) transformations to the traits and then regressed them against the covariates. The best transformation was chosen by visually inspecting the Q-Q plots of the residuals from these regressions and checking which one better approximated a normal distribution: in all cases the residuals were properly normalised. Supplementary Table S2 gives full details of the phenotype modelling.

**1.3 Genome wide association study (GWAS).**

Association analyses were conducted on SNPs imputed to the HRC panel[3], as provided by the UK Biobank, using BOLT-LMM[2]. Population stratification was assessed using LD-score regression as implemented in ldsc[4] both for both GWAS and after meta-analysis using the LD scores provided with the software: no evidence of residual stratification was observed. Table S15 reports the LD regression intercept and $h^2$ estimation using ldsc. Given that we identified 5 main clusters of traits (see paragraph 1.9 for details of group definition) we set the genome-wide significance threshold at $1x10^{-8}$. Genomic loci were defined by first removing all snps with $p>1e^{-5}$. Two adjacent loci were defined independent when the distance between two consecutive SNPs was greater than 250kb. Overlapping loci from different GWAS traits where then merged into a single locus.

**1.4 Replication Analysis**

Replication analyses were conducted independently by using genetic and dietary data from the EPIC-Norfolk Study[5] and the Fenland Study[6]. Both are on-gonig population based cohort studies conducted in the East of England. At baseline of the EPIC-Norfolk Study (1993-1997) and the Fenland study (2005-2015), the same food-frequency questionnaires (FFQs)[5,7] were administered to participants. Each participant was requested to report frequency of consumption of 131 food items by selecting one of nine categories of frequency of food consumption (‘never or less than once/month’, ‘1-3 per month’, ‘once a week’, ‘2-4 per week’, ‘5-6 per week’, ‘once a day’, ‘2-3 per day’, ‘4-5 per day’, and ‘6+ per day’). As performed in the analysis of the UK BioBank, the quantitative information on frequency of consumption was assigned to each response for 131 food items. We summed up frequencies of consumption of multiple food items in each food group (e.g. margarine and butter for bread spread; different types of vegetables for total vegetable consumption), so that resulting variables were comparable with those used in the UK BioBank.

Participants were genotypes with different genotyping arrays. In the EPIC-Norfolk Study, Affymetrics Axiom UKBiobank was used (n=21,044). In the Fenland Study, three arrays were used: Affymetrics Axiom UKBiobank (n=8,994), Illumina Metabochip (n=3,217), and Illumina ExomeChip v1.0 Human Exome-12v1-B (n=1,650). Further details are available elsewhere[6]. Missing information of SNPs was imputed to the HRC and UK10K panels by population and genotyping array. After we excluded participants without either dietary or genetic information, 32,779 participants in total became available for the replication analysis of EPIC-Norfolk Study (n=21,337; $n_{original}$=25,639) and the Fenland Study (n=11,442; $n_{original}$=12,731). Association analyses were conducted with BOLT-LMM in each of the four population-array strata.

**1.5 Univariate MR**

Univariate MR to measure the causal effect of health-related traits on food was conducted using the TwoSampleMR[8] R package. We decided to focus on traits for which dietary medical advice is given due to medical conditions in particular: body mass index (BMI), low density lipoprotein (LDL) Cholesterol, high density lipoprotein (HDL) Cholesterol, Total Cholesterol, Triglycerides, Diastolic and Systolic blood pressure,, Type II diabetes and Coronary Heart Disease. Educational attainment  was added as for it’s impact on food choices and as a proxy for socio economic status. The full list of studies from which the summary statistics were derived is given in Table S6.

For each trait we selected all SNPs with $p<5 \times 10^{-8}$ and $r^2<0.001$. We then performed stepwise heterogeneity pruning, first estimating heterogeneity using the Q statistic, if $p<0.05$, we removed one SNP at a time, looking at which removal would improve the statistic more. This procedure was repeated until $p>0.05$. We then tested if the intercept from the MR-Egger regression was different from zero ($p<0.05$). If this was the case, MR-Egger was used for the MR analysis otherwise the Inverse Variance method was used. We considered as significant those relationships in which the Benjamini and Hochberg FDR<0.05.

**1.6 Estimation of prior expected effect through bGWAS and genome-wide mediation analysis.**

One of the main issues in GWAS studies is to decide which covariates to apply in the regression model. When deciding to include a covariate or not, depending on the causal relationships between the traits, we may risk creating different types of biases. One approach could be to include in the model just the non-heritable covariates (i.e. sex and age) which will avoid spurious results due to collider bias. The problem with including heritable covariates is that the GWAS will also detect those SNPs which are associated with the covariates which are causally related to the trait, making the interpretation of the results harder. For example if educational attainment causally influences BMI, if the study is powered enough, it is possible that the genes from educational attainment show up on the BMI GWAS. Moreover, this limits the generalisability of at least part of the results to other populations in which the phenotypic architecture of the trait may be different. For example, following the previous example educational attainment was causal to the trait in the European population and not in East Asia the SNPs causal from Educational attainment will not be replicated in East Asian populations affecting also the generalisability of results.
It is thus extremely important to identify a technique which allows to distinguish between those SNPs which are directly causal of the trait of interest from those which are associated through other mediators.

There are 3 possible scenarios:

1) The covariate is causing the trait of interest in which case it would be correct to include it in the model (i.e. diet and socioeconomic status).

2) The trait of interest is causing the covariate in which case including the covariate in the model could result in collider biases.

3) The trait and the covariate are causing each other. In this case, using the covariate in a regression framework will correct for the overall effect while in truth we are interested in correcting only for the effect of the covariate on the trait and not vice versa.

So, in order to properly correct our analyses, we need to determine if a covariate is causing the trait of interest and at the same time estimate the size of the effect. In this respect, we can use two-sample MR to establish both causality and effect size, using a multivariable MR approach if there are multiple covariates. In principle, once this is done we can correct the phenotype given the covariates and run the GWAS based on this new corrected trait. However, given that in many cases information on covariates may be missing, a method which exploits available GWAS summary statistics would be more desirable. Such a method would need to first estimate which covariates are causal to the trait of interest and then based on their causal estimates, perform mediation analysis for each SNP. In the second step, for each SNP an expected mediated effect is estimated, combining the effect of the SNP on each of the causal traits with their multivariable causal effects. The expected effect can be subtracted from the observed effect on the trait of interest to get the “direct causal effect” of the SNP on the trait.

For the estimation of the prior expected effect, we used a Bayesian GWAS (bGWAS)[9]. The bGWAS approach leverages information from studies of related traits to carefully build informative priors for each SNP. To analyse food choices, we decided to include information from the same traits used for the univariate MR plus Crohn’s disease and ulcerative colitis. Given that the traits were meant to be used in a multivariable model, total cholesterol was removed to avoid strong collinearity with LDL and/or HDL cholesterol. MR was used to derive multivariate causal effects of the set of related traits on the different food choice phenotypes, using independent instruments (association p-value below 5 x $10^{-8}$ for at least one related trait, LD pruned $r^2$<0.001). For each food choice phenotype, a stepwise selection approach was used to select only the related traits significantly affecting the focal phenotype. To calculate the prior for SNPs on a given chromosome, we first apply multivariate MR (masking the focal chromosome) using the significantly related traits identified by the stepwise selection to estimate causal effects. We next use the causal effect estimates in combination with GWAS summary statistics of the related traits to estimate the prior effects. The prior effect of a SNP *i* ($\hat{\mu}_i$) is calculated using the observed standardised effects (Z-scores) for the *T* different related traits ($Z_{i,t}$ ) and the causal effects estimated masking one chromosome ($\widehat{\beta}_t$):

$$\widehat{\mu_i} = \sum_{t=1}^{T} Z_{i,t}\, \hat{\beta}_t \quad (1)$$

The prior estimated by bGWAS is on the scale of the z-score of the GWAS from the trait of interest, so the non-mediated z-score can be easily then estimated as the difference between the original z-score and the prior. The prior can be thought of as the total indirect effect while the corrected effect as the pure direct effect from mediation analysis[10]. Keeping the standard error constant, it is then easy to derive the corrected beta as z-corrected*se. It is important to note that when we estimate the prior expected effect we do not take into account the error of the multivariable estimates and we use the point estimates directly. This is because in MR the standard errors linked to each beta estimate are relatively large and if taken into account would lead to a very large final standard error and thus all pure direct effect estimates would have extremely large errors making them not usable. Although this is of course an approximation this is not unlike the estimation of polygenic risk scores where the SNP point estimates are used as weights for the score. It is important to note that for the further causal inference we used uncorrected betas and thus this will not influence the effect estimation.

This approach has several advantages compared to correcting the phenotype directly. First, it allows the GWAS to be corrected for covariates which have not been measured directly on the same samples. This is a great advantage in terms of the models that can be explored, for example in our case we have corrected the GWAS for LDL although LDL had not been measured in UK biobank at the time of the analyses, and this is also useful for phenotypes such as Crohn's disease, for which relatively few cases are present in UKB.

Moreover, it is possible to compare the effects before and after correction, giving us information on the likelihood the observed effect is directly associated with the trait of interest or is mediated. it is important to remember that conditioning on a phenotypic covariate will not necessarily completely correct the mediated effect, due to noise, and thus the comparison of the two effects is more informative. Finally, we should be able to trace back the path of mediation looking at the different components of the prior for each SNP thus helping greatly in interpreting the results and planning subsequent studies.
All the exposure traits GWAS have been first imputed using SSimp[11] (https://github.com/zkutalik/ssimp_software) and the UK10K genotypes as the reference panel. Finally, all A/T or G/C SNPs were removed to avoid errors in the harmonisation of effects coming from different GWAS, due to strand errors. The proportion of genetic variance of the food traits explained by the health-related traits was measured by taking the squared genetic correlation between the expected Z-score and the observed one.

**1.7 Identification of the SNPs directly associated with the food traits.**

One of the main objectives of GWAS is to identify genes directly responsible for the trait of interest, in our specific case, however, we have shown that looking just at the genome-wide hits is not sufficient and does not exclude SNPs truly associated with other causal heritable traits, due to vertical pleiotropy. In order to distinguish between these two types of SNPs, we decided to look at the ratio between the corrected trait and the original trait or corrected-to-uncorrected ratio (CUR). To understand this choice let's suppose we have a trait of interest (Y) and a second heritable trait (X) which is causal with effect, $\beta_{X \to Y}$. We will call SNPy, the SNPs directly causing Y with effect, $\beta_{SNPy \to Y}$ and SNPx, the SNPs which are directly causing X with effect, $\beta_{SNPx \to X}$. If the whole effect of SNPx on Y is mediated through X its effect will be given by

$\beta_{(SNPx \rightarrow Y)expected} = \beta_{SNPx \rightarrow X} \times \beta_{X \rightarrow Y}$ (2)

$\beta_{X \rightarrow Y}$ can be estimated through MR, while $\beta_{SNPx \rightarrow X}$ can be retrieved from the GWAS of X. Assume we measure the effect of a SNP for which it is unknown if the effect is mediated through X or not (as is the case in real data). We define $\beta_{SNP \rightarrow Y}$, the observed effect of the SNP on Y

if $\beta_{SNP \rightarrow X}$ is truly 0, then we can estimate the expected mediated effect of the SNP through X as

$\beta_{(SNP \rightarrow Y)expected} = \beta_{SNPx \rightarrow X} \times \beta_{X \rightarrow Y} \approx 0$ (3)

and

$\beta_{(SNP \rightarrow Y)corrected} = \beta_{SNP \rightarrow Y} - \beta_{(SNP \rightarrow Y)expected} \approx \beta_{SNP \rightarrow Y}$ (4)

thus

$CUR = \frac{\beta_{(SNP \rightarrow Y)corrected}}{\beta_{SNP \rightarrow Y}} \approx 1$(5)

On the other hand if

$\beta_{SNP \rightarrow X} \neq 0$ (6)

then

$\beta_{SNP \rightarrow Y\ expected} = \beta_{SNP \rightarrow X} \times \beta_{X \rightarrow Y} \neq 0$(7)

$\beta_{(SNP \rightarrow Y)corrected} = \beta_{SNP \rightarrow Y} - \beta_{(SNP \rightarrow Y)expected} \neq \beta_{SNP \rightarrow Y}$ (8)

then

$CUR = \frac{\beta_{(SNP \rightarrow Y)corrected}}{\beta_{SNP \rightarrow Y}} \neq 1$(9)

***Figure S1. Directed acyclic graph explaining the two possible scenarios for the effect of a SNP on the trait of interest Y. (a)*** *The SNP has a direct effect on Y not mediated through X. Then the estimated effect of SNP on X will be normally distributed around 0, thus the corrected and uncorrected effects will be similar and their CUR will be close to 1.* ***(b)*** *The SNP effect is mediated through X, thus the corrected effect will deviate from the observed one and CUR will deviate from 1.*

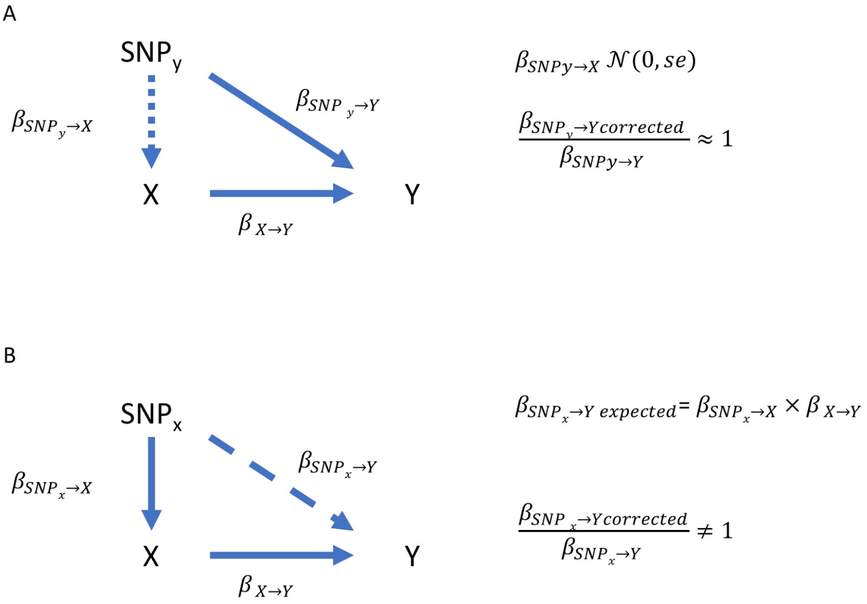


In real-life situations, the betas are estimated and thus will carry an error due to chance and it is thus important to understand what values the CUR may assume under different scenarios. We can, however, summarise three types of relationship between the trait of interest and its causal factors:

1) X→Y

2) X→Y and Y→X

3) U→Y and U→X

Where U is a heritable confounder responsible for the relationship between X and Y. We thus use simulations to understand how the relationships between the traits influence the CUR and our ability to use it to distinguish between SNPs, the effects of which are mediated through X or U, and SNPs which are causal to the trait of interest without mediation. The details and results of the simulations are reported in Supplementary Data 2.1.

**1.8 Clustering of food and drink consumption traits.**

Genetic correlations between the food traits were estimated using LD-score regression as implemented in the ldsc software[35] separately for the corrected and uncorrected GWAS. Hierarchical clustering of the two genetic correlation matrices was performed using two different algorithms: “complete” as implemented in the hclust() function from R and the ICLUST algorithm[17] from the R package psych. ICLUST assigns items to the same cluster based on the loadings of an underlying common factor. Items are then iteratively added to the clusters only if they increase the internal consistency of the cluster. The algorithm also allows for addition of traits in case of strong negative correlation. This has a compelling advantage for food consumption, as for example the intake of fatty foods has a strong negative correlation with eating healthy food, thus both can contribute to the same grouping. Differences in clustering were compared graphically using a tanglegram in both cases. Given that ICLUST seemed to give more stable results compared to the “complete” clustering algorithm, the clusters produced with this algorithm were used for further analyses.

**1.9 Multi-trait genome-wide association analysis**

For each of the three main clusters of phenotypes (Meat/Fat, Healthy foods and Psychoactive Drinks), we performed multi-trait genome scans using a MANOVA-based multivariate analysis method implemented in the MultiABEL package[37] (https://cran.r-project.org/package=MultiABEL). The method can take genome-wide summary association statistics to infer phenotypic correlation coefficients and conducts a MANOVA test for each variant across the genome. This overcomes the issue of non-overlapping samples (e.g. it would be impossible to directly combine people drinking different type of coffee). The phenotypic correlation coefficient of any two traits can be estimated in an unbiased manner via the correlation of the genome-wide z-scores, and for binary outcomes, this is proportional to the phenotypic correlation on the liability scale. The MultiABEL package also calculates the best linear combination of multiple phenotypes that is associated with each variant.

**1.10 Locus definition and prioritisation of genes.**

To define a locus, we first selected all SNPs with p-value $<1\times10^{-5}$ and then estimated the distance between each consecutive SNP located on the same chromosome. Two consecutive SNPs were identified as belonging to different loci if they were more than 250 kb apart. A locus was then considered significant if it contained at least one SNP with p-value below the previously described significance threshold: we thus identified 582 significant locus-phenotype associations. Given the high pleiotropy between different traits, we merged overlapping loci, which resulted in 302 independent loci.

In order to define for each locus which gene is more likely to be responsible for the observed association, we proceeded with custom prioritisation according to the following criteria. We first ran Haploreg v4.1[38] using $r^2=0.8$ as threshold (Supplementary Data 11). We also ran SMR[39] on each locus in order to identify eQTLs compatible with the observed association pattern. We used the tissue-specific significant eQTL from the Gtex Project website[26] (Supplementary data 12). We then proceeded to prioritise the genes according to the following criteria; if the locus met one of them the following ones were not tested:

1) The sentinel SNP is itself or is in strong LD ($r^2>0.8$) with a non-synonymous SNP
2) There is evidence of an eQTL (as tested with SMR) and the sentinel SNP and the eQTL are in strong LD (min $r^2=0.5$). Given the high number of significant eQTLs detected by SMR we used a dynamic selection starting from $r^2\geq0.99$ and decreasing by 0.05 at each step until an eQTL was found or until $r^2\leq0.5$.
3) The sentinel SNP is itself or is in strong LD ($r^2>0.8$) with a coding SNP (synonymous or in the untranslated region of the gene)
4) The top SNP is intronic or is in complete LD with an intronic SNP in the gene.
5) The top SNP is in strong LD ($r^2>0.8$) with an intronic SNP in the gene.
6) The closest gene.

The category and prioritised gene for each locus is annotated in supplementary table 3.

##### 1.11 Prioritised gene annotation and network construction

Tissue enrichment using MAGMA[12] was run using FUMA[13]. For each available SNP, we chose the lowest p-value from all corrected analyses. For the functional annotation of the prioritised genes, we focused only on those coming from the "direct effect only" loci. First, we used the gene2Function tool from FUMA to identify enrichment in the same tissue used for the analysis with MAGMA. For both analyses, only tissues with Bonferroni corrected $p < 0.05$ were considered significant (Table S11-S12 report full results).

We then constructed an interaction network using STRING[14] (Table S13). After removing the genes which were not connected with any of the others, we ran community detection using Leuvain's method (Table S14 for membership). Tissue enrichment analysis was then performed for each community as done for the full gene set, focusing only on the overexpressed tissue analysis. Given the much higher number of tests performed, we used Storey's q-values to define significant tissues. Gene ontology enrichment was performed for each community using the compareCluster() function from the clusterProfiler R package[15]. We considered significant those terms which had a FDR<0.05 using the Benjamini and Hochberg method.

**1.12 Selection of genetic instrumental variables for dietary patterns.**

Given that we did not run a GWAS for each PC trait the genetic instrumental variables for the dietary patter traits were selected as follows:

1) All SNPs with $p<5e^{-8}$ and CUR =1±0.05 in each of the food traits were merged together.
2) To each SNP was assigned the lowest p-value across all used traits
3) LD pruning was applied using r2<0.001 as threshold
4) The effect of the SNP on the PC trait was calculated as the linear combination of the betas using as weights the loadings from the eigen decomposition of the genetic correlation matrix
5) SE where estimated using the same weights and the phenotypic correlation matrix between the traits.

Figure S3 reports the corrplot of the loadings of each single item on the PC traits.

**Figure S3.** Corrplot of the loadings of each food item onto the measures of dietary pattern. All items have been aligned to the “Overall unhealthy diet” measure. The items that have been flipped are noted as “Less” to clarify the direction of the relationship.

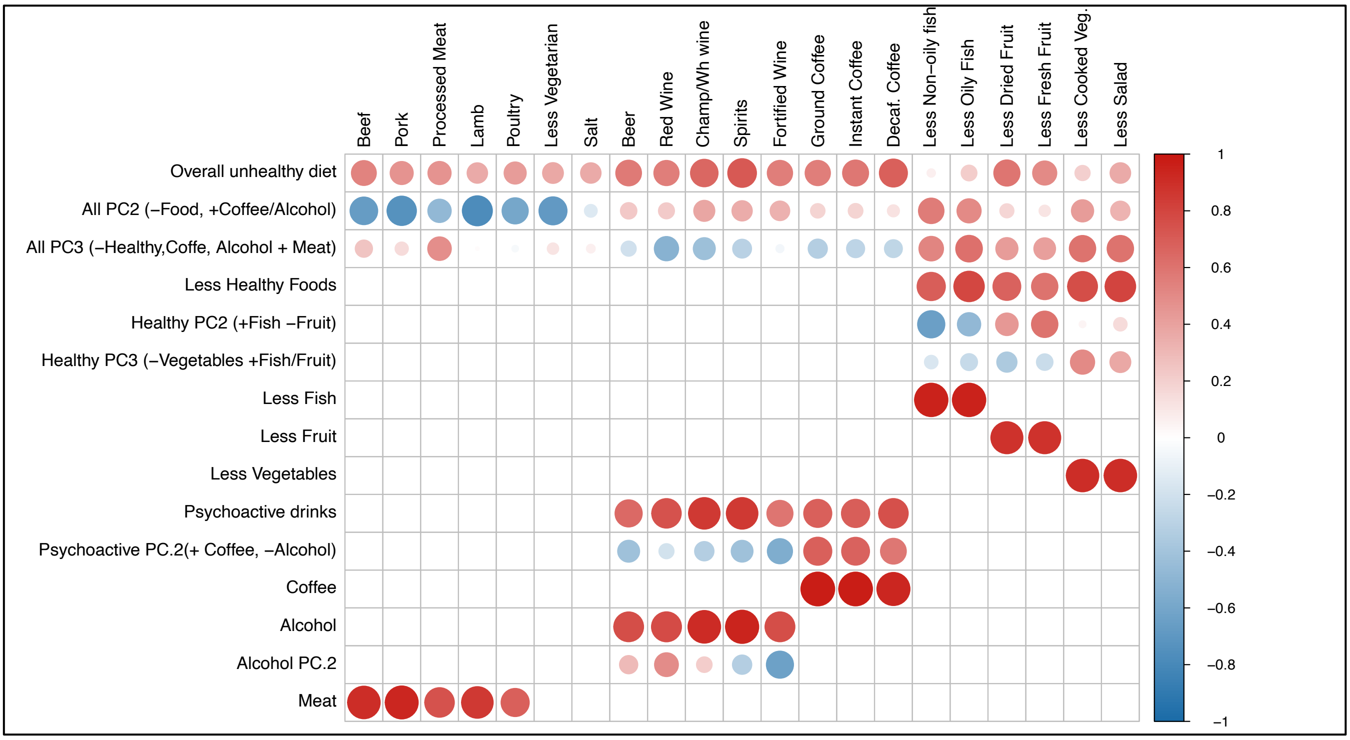

#### 2. Extended Figures/Results

***Fig S4. Results for the Mutlivariable MR.*** *Panel A The heatmap represents the effect of the health related traits on each food trait using from the multivariable model. The color is proportional to the effect size. Panel B. The plot represents the proportion of genetic variance which is explained by the effect of the health related traits on the food traits. Clearly some of the food traits are extremely biased having up to 40% of genetic variance due to the mediation of the health related traits.*

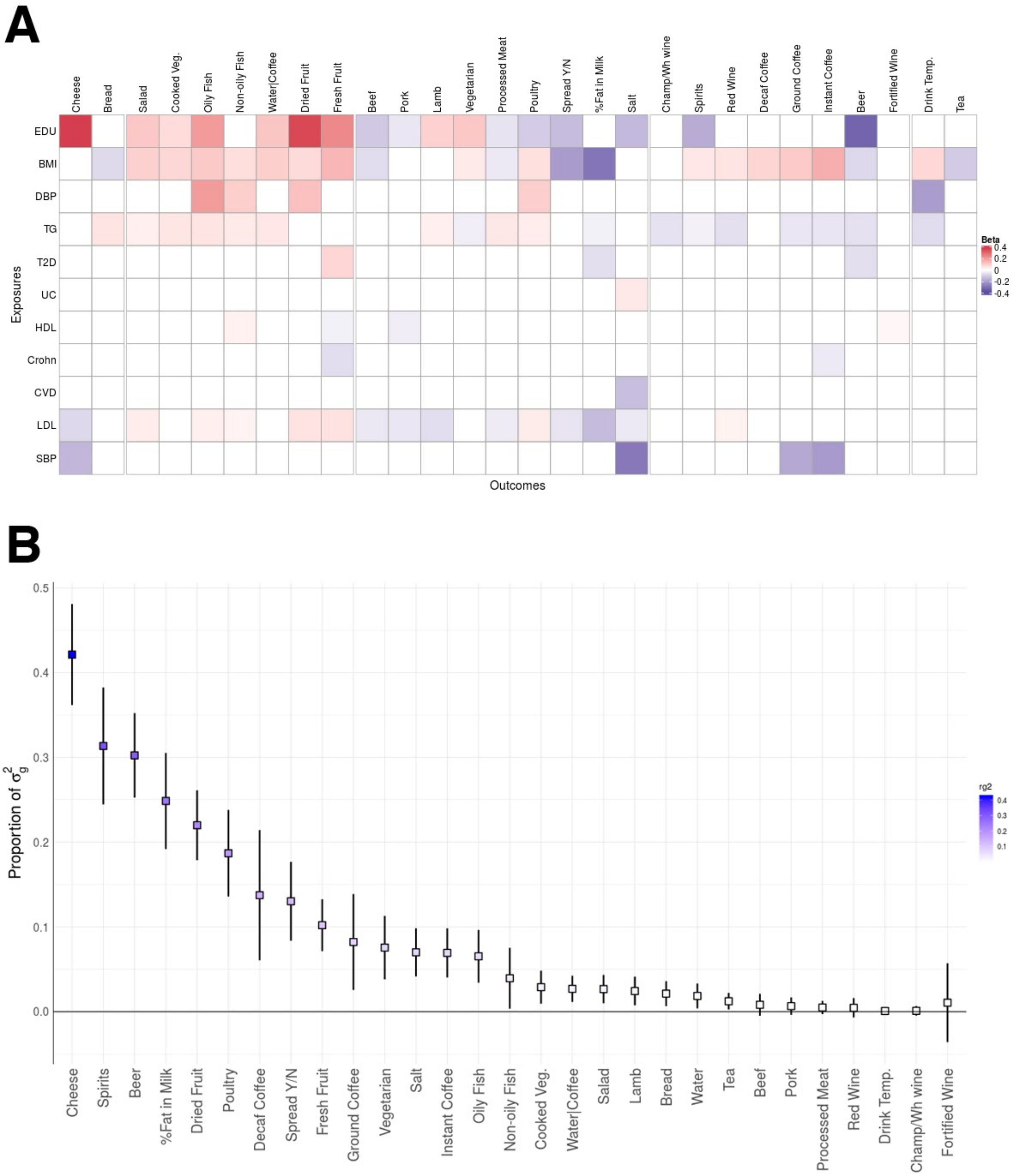

#### 2.1 Simulation results for optimal parameter tuning for selecting SNPs directly associated with the trait of interest.

The objective of the simulations was to understand if the corrected/uncorrected beta ratio limits chosen based on the genes for which biology is well known are correct. The simulations are particularly complex to set up since the DAG of the studied relationships involves bidirectional causal effects. Another important limitation is the fact that the exposure trait is not directly and correctly observed but it is the result of a FFQ in which the noise is extremely high with a test-retest correlation that can be as low as 0.5 ($r^2$=0.25).

The simulations include 4 different normally distributed traits:

$Y_t$ which is the true trait of interest (food consumption in our case) without the effects of the outcome. X and U represent each the sum of all traits causal to Y. The difference between X and U is that U traits are also causal to X (so they act as a confounders) while X traits may also be subject to reverse causality by $Y_t$. $Y_o$ is the observed trait which thus includes all causal effects and the noise due to the use of the questionnaire.

**Figure S5.** Diagram describing the relationships between the simulated traits and their relative parameters. Gy refers to the genetic variants which directly affect Y before any influence of confounding or other mediated traits (Yt). Gu represents the genetic component of a confounder trait U which causally affects both Y and X. Gx represents the genetic component of the outcome trait X which is in turn causally affecting the trait Yi. Yo represents the actual observed trait to which we add noise to reflect the test-retest correlation in FFQ data.

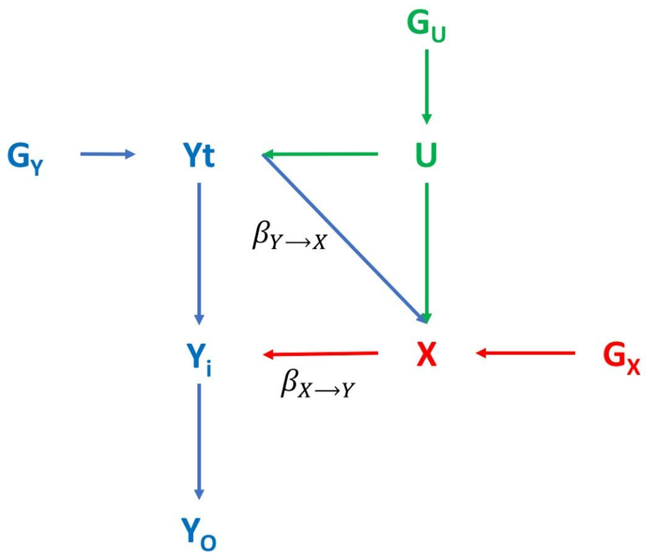

For simplicity each of the first 3 traits is determined by 10 SNPs each of which has a frequency of 0.3 and explains 1% of variance. Overall 30 SNPs have been used for each simulation and they are denoted as $SNP_Y$, $SNP_X$ and $SNP_U$ depending if their direct effect is through Y, X or U. The relationships between the different traits are summarised in Figure S4.

Where:

$\beta_{U\text{->}Y}$ represents the effect of the confounder on $Y_t$,

$\beta_{U\text{->}X}$ represents the effect of the confounder on X,

$\beta_{Yt\text{->}X}$ represents the effect of $Y_t$ on X,

$\beta_{X\text{->}Yt}$ represents the effect of X on $Y_t$.

$\beta_{Yo\text{->}X}$ represents the causal effect we would be able to measure through MR. This measure is not of interest for the scope of our simulations.

For simplicity the $\beta_{U\text{->}Y}$ and $\beta_{U\text{->}X}$ were both set so that the confounder explained 20% of the variance of Y and X. Simulations were then conducted for a large array of values of $\beta_{X\text{->}Yt}$ and $\beta_{Yt\text{->}X}$ , which ranged from 50% of variance explained to 0%, with both positive and negative effects. Values of $\beta_{Yt\text{->}X}=0$ simulates the case where no reverse causality exists. A denser grid was used between $r^2=0\text{-}0.1$ to examine more closely the results at smaller effects which likely better resembles most real cases. Each set of parameters was run 10 times so that 100 SNPs were simulated for each category and set of parameters. To replicate the conditions of the paper we simulated two different independent populations so that we could apply the MV MR correction procedure to study the effects in a setting which resembles the real life scenario. Both populations were simulated so that N=400,000.

After simulating the two populations we proceeded to perform the association analysis for all 30 SNPs with all three observed traits, leaving out the original trait of interest $Y_t$, assuming we would not be able to directly observe it (as is the case for food consumption measured with FFQ). We then performed the MV MR of X and U on Y using as IV all SNPs which had $p<5\text{x}10^{-8}$ in either the GWAs from X or U assuming we had no way of distinguishing the source of the SNP. For this analysis we used population 2 for the exposure betas and p values while population 1 for the effects of Y. Finally we used the betas combined with the population one $Y_o$ association results to estimate the corrected/uncorrected ratio (CUR).

The main objective was to verify if using the proper limits of the corrected/uncorrected ratio allows the $SNP_Y$s to be correctly distinguished from the $SNP_X$s and the $SNP_U$s. Figure S5 shows the scatterplot of the CUR for the 3 categories of SNPs for each combination of simulated parameters. $R^2$ refers to the amount of variance explained by the causal trait. The values are both positive and negative to reflect the direction of the correlation. The CUR range for the plot is limited to values between 0 and 2 because $SNP_Y$s never assumed values outside this range.

***Figure S6***. *Scatterplot of the CUR values for Gx (in red), Gy (in green) and Gu (in blue) at the different values of the effect of Y on X and of X on Y.*

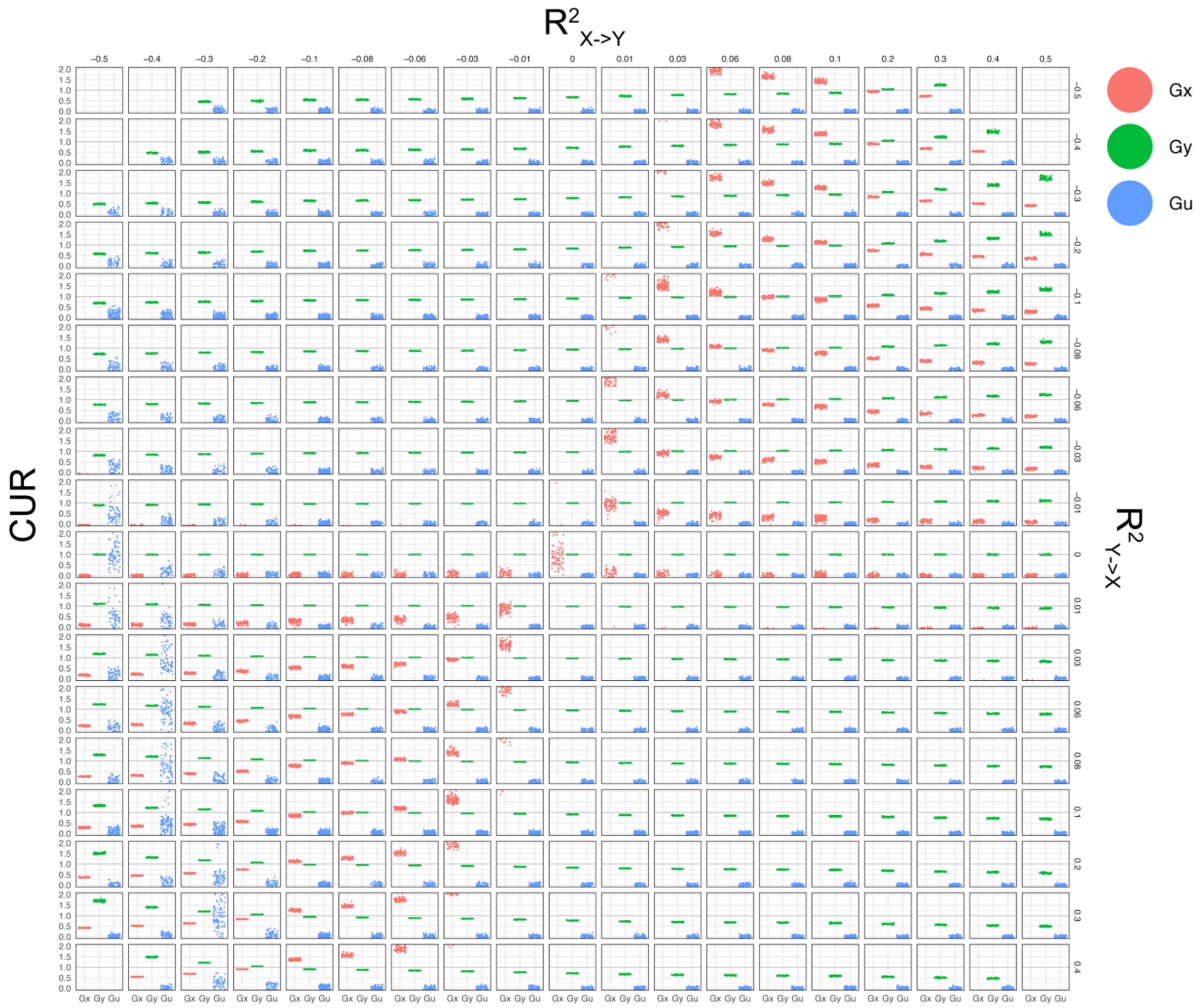


Clearly for most combination of parameters it is possible to easily separate the $SNP_Y$s from the $SNP_U$s. The task becomes slightly more complex in the case of the $SNP_Y$s and $SNP_X$ in which some overlap is possible, especially when $\beta_{X\text{->}Y}=\beta_{Y\text{->}X}$ however this particular case is probably unlikely in real case scenarios as it would mean that the two effects cancel each other out. From the previous figures, it is clear that if we were to know $\beta_{X\text{->}Y}$ and $\beta_{Y\text{->}X}$, we would be able to determine which values of CUR to use for the selection of the IVs in most cases. However without knowing these *a priori* it is however quite difficult as they would require prior knowledge of valid IVs for both Y and X, which is the objective of the method. Thus, the real question is if there is a range of values for the CUR which maximises the probability of not discarding the $SNP_Y$s while not including $SNP_X$s or $SNP_U$s.

Therefore we verified how the probability of actually detecting a $SNP_Y$, P(detection), and the probability that an IV which met the criteria was actually a $SNP_Y$, P($SNP_Y$), at different ratio limits.

For the values selected for our study (1±0.05), the P(detection) is 0.27 overall, and 0.57 for the combination of the weak effects, while P($SNP_Y$) is >80% in both cases (Figure S6). Given that, for almost all traits we have at least 1 SNP which meets these criteria and that loosening them will increase the chances of including $SNP_X$s as instruments, the choice of parameters used so far seems reasonable.

***Figure S7. Corrected-to-uncorreected ratio (CUR) successfully distinguishes mediated and non-mediated associations.*** ***(a)*** *Graph showing mediated and non-mediated pathways. The values of CUR that different types of simulated SNPs (Gx, Gy, Gu) assume at different explained variances ($\sigma^2$) of X->Y when $\beta$(Y->X)≠0, i.e. presence of reverse causality* ***(b)****. The values we used for defining a "non-mediated" variant are highlighted in purple.* ***(c)*** *The proportion of variants that are truly Gy, that is directly associated with the trait of interest, across a range of CUR.* ***(d)*** *The overall proportion of variants directly associated with the trait (SNPy) whose CUR falls inside the specified ranges, i.e., the probability of detecting SNPy over all possible scenarios.When the effect of Y->X is equal to zero, Gy is clearly distinguishable from Gx and Gu using CUR (Fig. S5), however, when $\beta$(Y->X) increases, values of CUR for both Gy and Gx start varying and overlapping (Fig. S6b). We thus determined which values of CUR would maximise the probability of correctly selecting Gy under all scenarios. Clearly the parameters we have chosen for defining a "non-mediated" SNPs maximise both the probability of  correctly selecting a SNPy.*

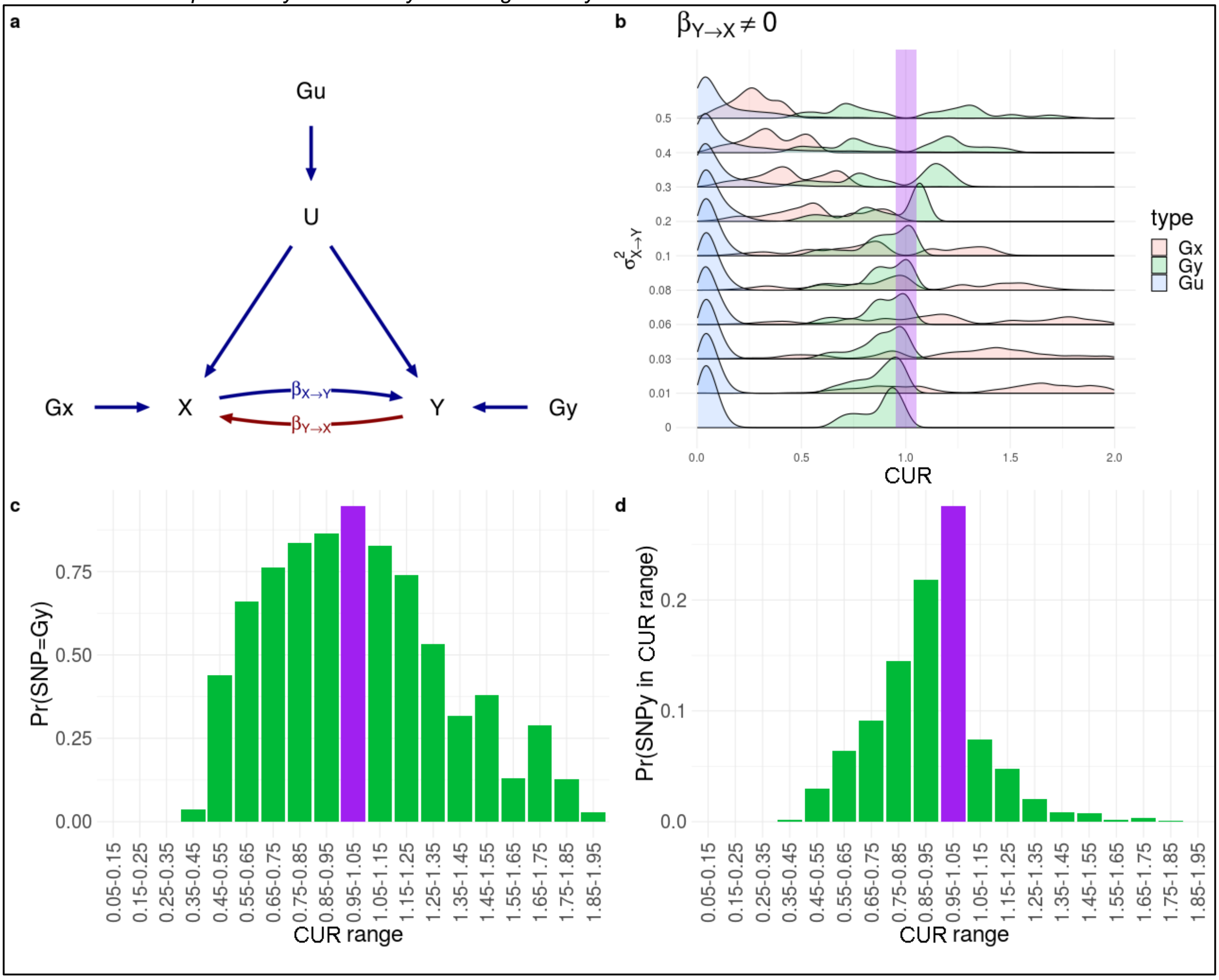

**2.2 Effects of GWAS correction on genetic correlations.** To investigate how the mediation procedure affected the genetic correlations amongst the consumption traits, we compared the correlation patterns using the uncorrected and corrected results (Table S10). Overall, the two genetic correlation matrices were very similar, but with some important differences. In particular, the number of Bonferroni-corrected significant correlations diminished in the corrected results, reflecting the fact that the conditioning on the health-related traits will weaken those correlations which are partly due to those traits. The hierarchical clustering of the traits (Fig. S7a) using the two different matrices shows an improvement in the quality of the clustering with the groups formed using the corrected $r_G$ matrix being more interpretable than the uncorrected ones. While the group composed of healthy foods is stable across methods, we can see that using the uncorrected $r_G$s, salt clusters with beer and strongly alcoholic beverages, while wine clusters closer to coffee than other alcoholic drinks, perhaps reflecting medical advice. Also, the fatty foods (percentage fat in milk and adding spread to bread) cluster in an unexpected way, grouping closer to healthy foods than to meat. Looking at the corrected $r_G$s, clustering accords better with common sense, for example fatty foods grouping with meat and salt.

***Fig S8. Clustering of food consumption traits before and after correction.*** *Comparison between the hierarchical clustering of the food traits based on the uncorrected (on the left) and corrected (on the right) genetic correlations. Black lines connect the same traits for which the clustering has changed. Dendrograms connect the items in each case with the boldness of the line representing the strength of support for the tree nodes. Unique nodes are represented with a dashed line while shared nodes with a bold one. The thickness of the line is thicker for conserved higher level nodes.*

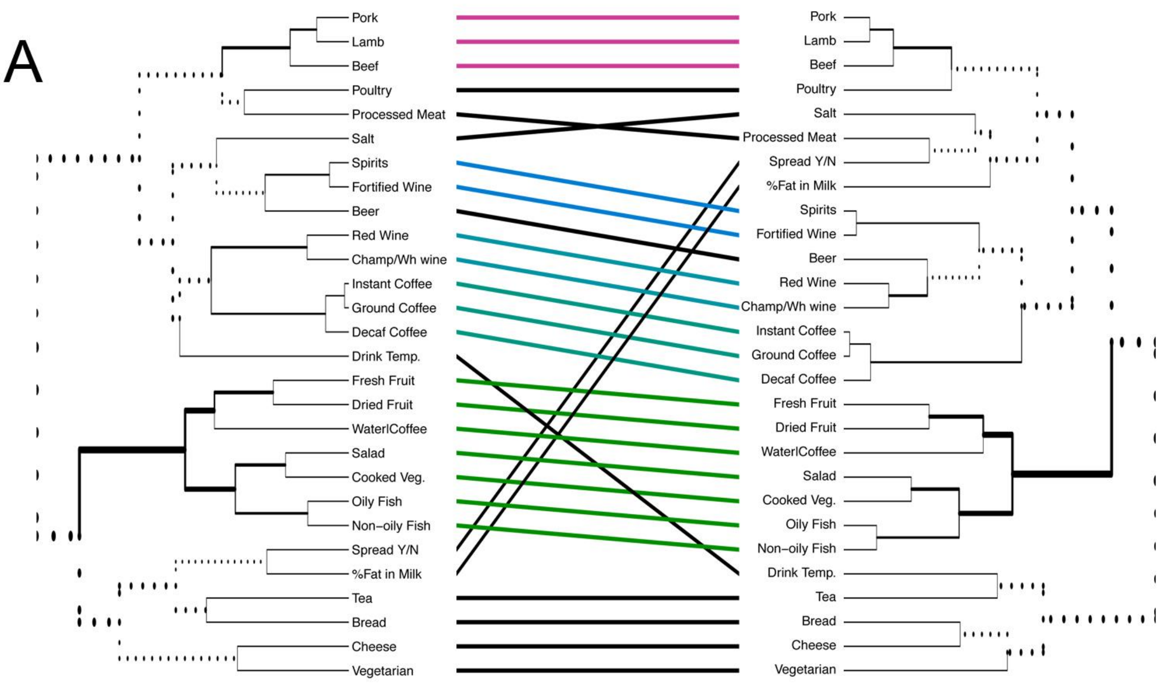


The clustering results show that the mediation correction procedure has been at least partly able to remove the genetic correlations due to shared heritable factors, rather than common biology. Such correction is extremely important, not only for creating homogeneous clusters of traits for multivariate GWAS, but also because these may change in a population- or age-dependent manner. It is reasonable to believe that some of these strong mediating factors (e.g. LDL cholesterol) are due to the older age of the samples in UKB (indeed ~27% are prescribed lipid-lowering therapy and will thus likely have been given medical advice to change their diet). It will be interesting to compare with a younger population less affected by perceived or actual medically advised lifestyle changes.

To explore the shared genetic underpinnings of food choices and a broad range of complex traits, we estimated genetic correlations with 832 traits present in LDhub[16]. We identified 6967 and 4943 significant (FDR<0.05) genetic correlations for uncorrected and corrected traits, respectively across a large number of traits (Table S10, interactive view available at https://npirastu.shinyapps.io/rg_plotter_2/). The correction affected greatly not only the genetic correlations with the traits used for the correction but also those with many others. We can only highlight a number of examples of changes in genetic correlations here. A notable instance is that prior to adjustment, CVD and percentage fat in milk showed a genetic correlation of -0.24, i.e. decreasing the %fat increased the chances of CVD, but after correction, $r_G$ was 0.02. Another example is again cheese consumption which has a genetic correlation with a longer paternal lifespan of 0.5 before adjustment, but only 0.2 afterwards. These results are particularly important because they suggest that the recent epidemiological findings associating higher consumption of fat in milk with protection from CVD[17,18] may be due to confounding and caution should be used when defining dietary policies.

**2.3 Multivariate association analysis.** Clustering of the traits using ICLUST identified five different groups (Fig 2): one composed of increased meat, fat, salt and decreased vegetarianism (labeled as “Meat/Fat”), one made up of alcoholic beverages and coffee (labeled “Psychoactive”) and one comprised of healthier items such as fish, fruit and vegetables (labeled “Healthy Foods”). Two final groups contained only two items each: drink temperature and tea; and cheese and bread; these were not used for the MV analysis. In order to explore if additional loci influence these groups we ran a multivariate GWAS using the package MultiABEL, which performs MANOVA on summary statistics. 168 additional associations, including 42 novel loci not identified at the single trait analysis, were identified in multivariate analysis of the three main food groups (Table S5) An example of these group-level loci is rs17400325, a non-synonymous variant at *PDE11A* associated with the consumption of Low Calorie Foods. When each trait was examined singularly we found that the C allele was associated with higher fresh and dried fruit consumption and a lower consumption of fish and vegetables. Mutations in this gene are responsible for primary pigmented nodular adrenocortical disease-2 (OMIM:610475), which leads to high cortisol levels, which in turn are associated with a higher consumption of highly palatable foods[19].

**2.4 Functional annotation of food consumption genes.** We used several approaches to understand the biological underpinnings of food choice. First we ran stratified LD-score regression[20] using the bias-corrected GWAS (Fig. S8). Looking at functional annotation (Fig. S8a), we found a strong enrichment for almost all food traits in conserved genomic regions with the exception of being vegetarian, decaffeinated coffee and fortified wine consumption. This is not surprising if we consider that nutrition is one of the most basic biological functions. We next looked at tissue enrichment in the Gtex[21] and Franke lab expression[22] datasets and epigenetic signatures from Roadmap[23]. There was substantial agreement across the two expression datasets with enrichment mostly limited to brain areas linked to reward and feeding, e.g. hypothalamus, nucleus accumbens, putamen. Most food traits were also enriched for epigenetic marks annotated to the male and female fetal brain, which underlies the importance of foetal development in determining food choices.

***Fig S9. Heatmap of tissue and functional enrichments.*** *The colour is proportional for the enrichment revealed by stratified LD-score regression. Only correlations with FDR<0.05 are reported.* ***(a)*** *Enrichment among different classes of functional annotation.* ***(b)*** *Tissue enrichment from Gtex expression.* ***(c)*** *Tissue enrichment from ROADMAP epigenetics.* ***(d)*** *Tissue enrichment from the Franke lab dataset.*

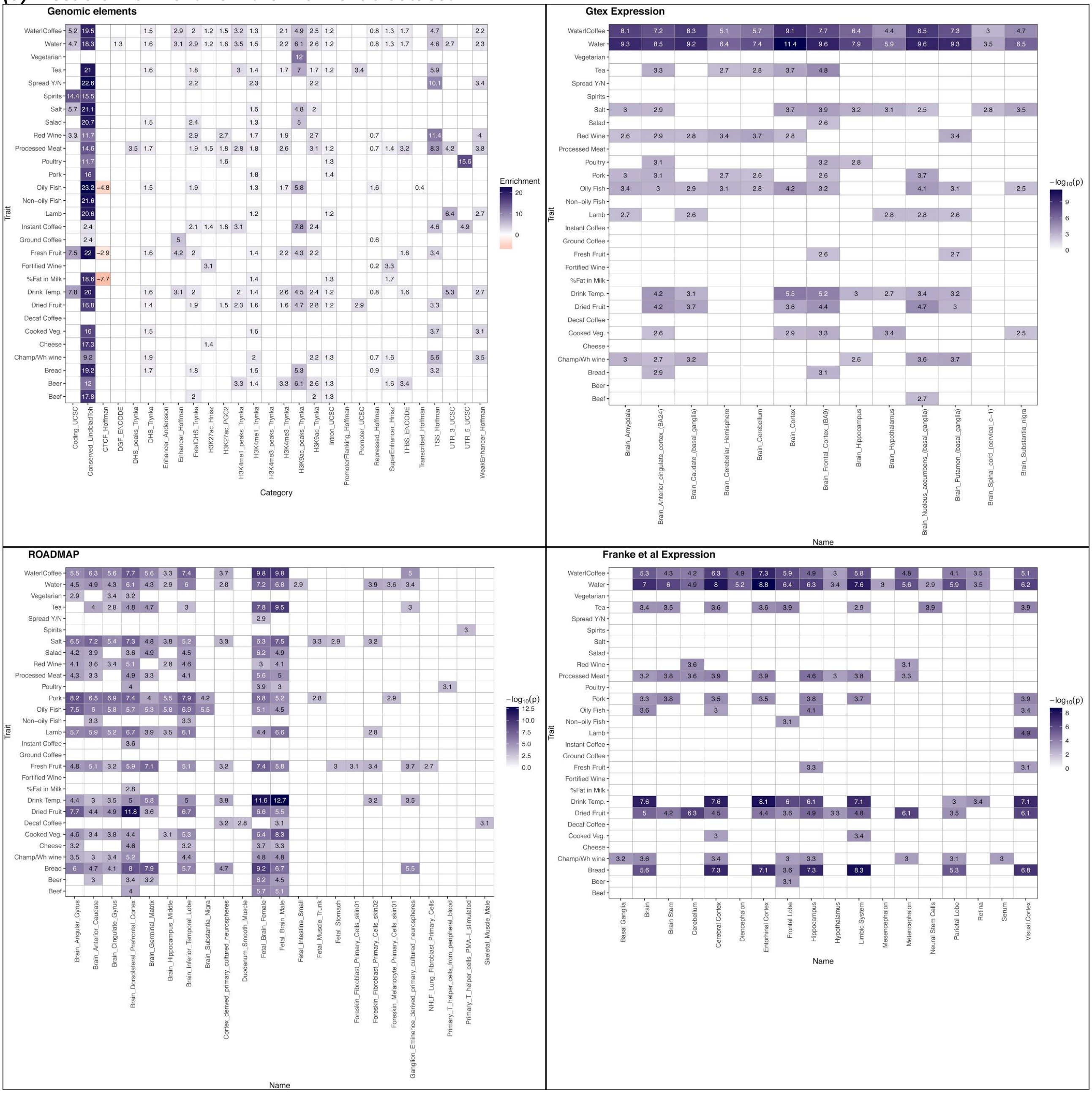


Tissue over-representation analysis using MAGMA[12] (which first runs a gene-wide test and then measures enrichment using those which results significant) on the merged GWAS results (for each SNP the lowest p-value was used) confirmed the results from LD-score regression highlighting the same brain areas, e.g. substantia nigra, nucleus accumbens, hypothalamus, amygdala, known to influence food choices and reward (Fig. S9). Very similar results were obtained when the analysis was conducted using only the prioritised genes in the "direct effect only" loci. In this case, as well as over-expression in the brain, we also detected under-expression in the kidney, stomach, pancreas, oesophagus, colon, lung and small intestine (Fig. S9).

***Fig. S10.* Dotplot of the overexpression analysis run on the prioritised genes from the non-mediated loci and the overrepresentation analysis performed with MAGMA.** The overexpressed tissue involved by the two methods were highly overlapping with the analysis performed on the prioritised genes showing also the tissues in which there is evidence of underexpression.

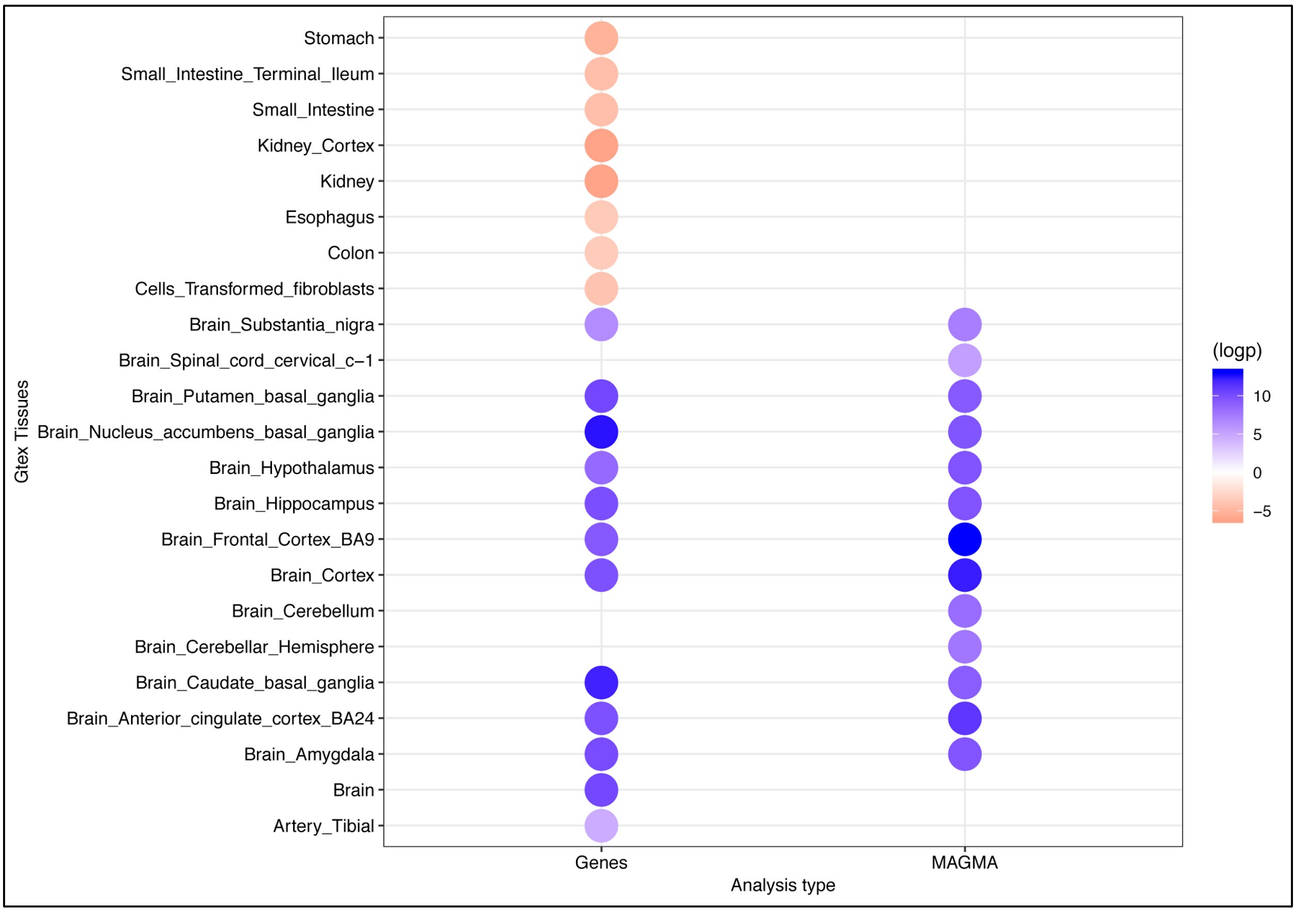


**2.6 Network analysis.** The fact that the genes we prioritised show the same enrichment pattern as stratified LD-score regression and MAGMA, also suggests that the prioritisation is in most cases correct. In order to explore any interactions between the selected genes, we used STRING[24] to build an interaction network between them. A large network is revealed (Fig. SXX, Table S13), including 132 genes (out of 215 overlapping the STRING database), sharing 224 edges ($p<1 \times 10^{-16}$).

***Fig S11. STRING network of genes in non mediated loci***. *Network plot of the genes in the non-mediated loci. After performing community detection we identified ten different clusters of genes each with its particular set of functions and expression patterns (see additional paragraph 2.6 for details). Nodes have been colored according to community membership.*

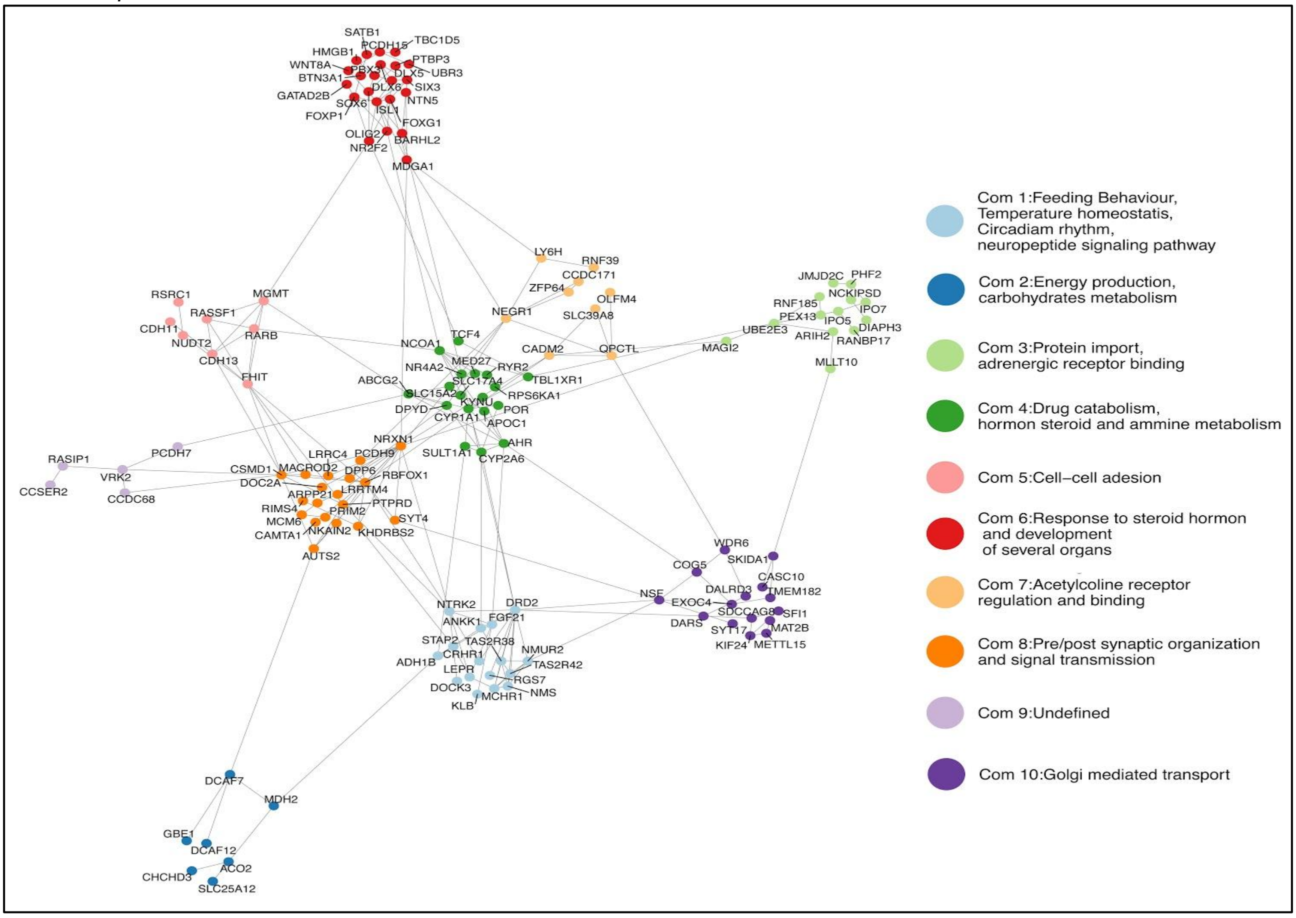


To identify if there were groups of genes which were more interconnected than the others, we performed community identification using Leuvain's method[24,25]. Ten communities are identified, each with specific characteristics in terms of function, cellular localisation and preferential expression (Fig. S11 for and overview and FigS 12-21 for specific comunities, Table S15 for significant Gene Ontology terms, Fig. S10, Table S16 for significant tissue enrichment). For example, community 1 genes are linked to numerous biological processes ranging from feeding behaviour and taste to hormone binding and transport of fatty acids, and preferentially include the genes expressed in several brain areas and the liver. Genes in community 2, on the other hand, are linked to energy and glucose metabolism, are preferentially located in the mitochondrion and are over-expressed in the skeletal muscle and tibial artery. Another interesting example is community 8 which contains genes specifically over-expressed in the brain and which are involved in synaptic assembly and organisation and in neurotransmitter secretion, while other communities relate to steroid hormone response, acetylcholine receptor regulation, drug metabolism and Golgimediated transport. Thus, although the overall expression analysis strongly links dietary choices to the central nervous system, there are actually several different groups of genes at play, with specific functions in specific tissues.

***Figure S12 Tissues which overexpress the genes in each community.***

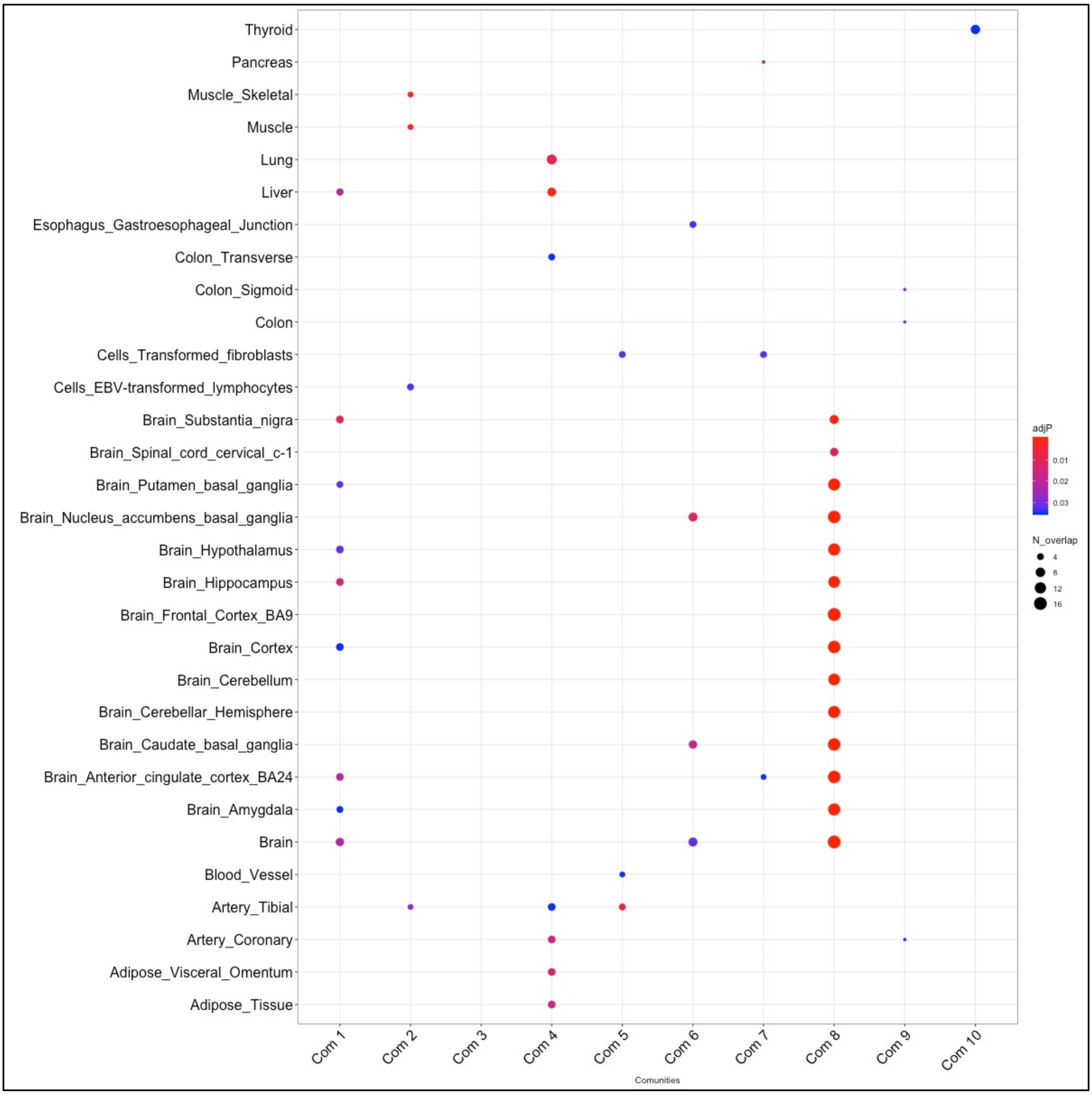

***Figure S13 Overlap in Go-terms between different communities.*** *The figure shows that there is no overlap (with the exception of 2 terms) between the terms enriched in each community. The labels have been removed as the plot is meant to only show the overlaps. Figure S12-22 show the enriched terms for each community separately.*

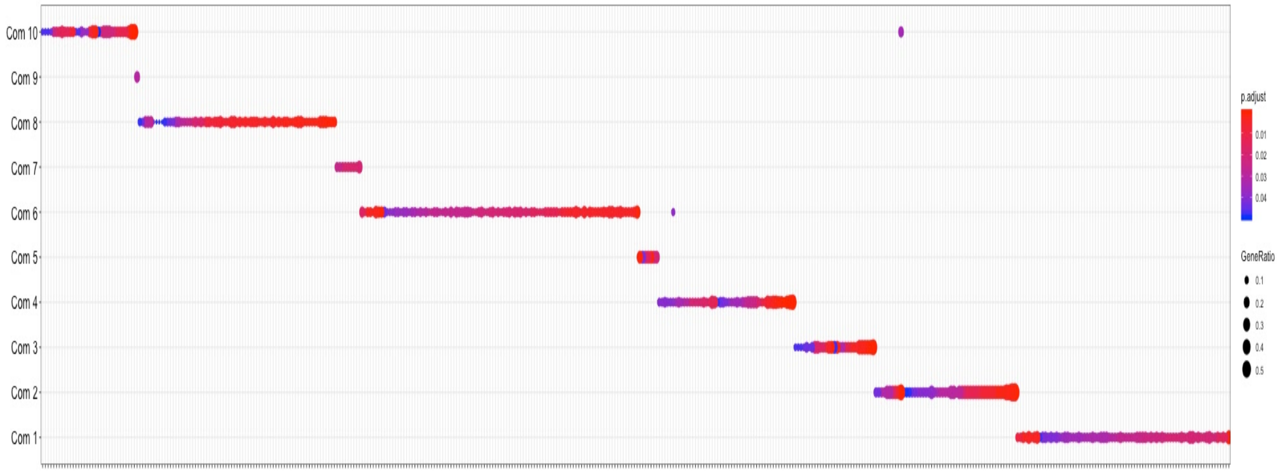

***Figure S14 Enriched GO-Terms for community 1.***

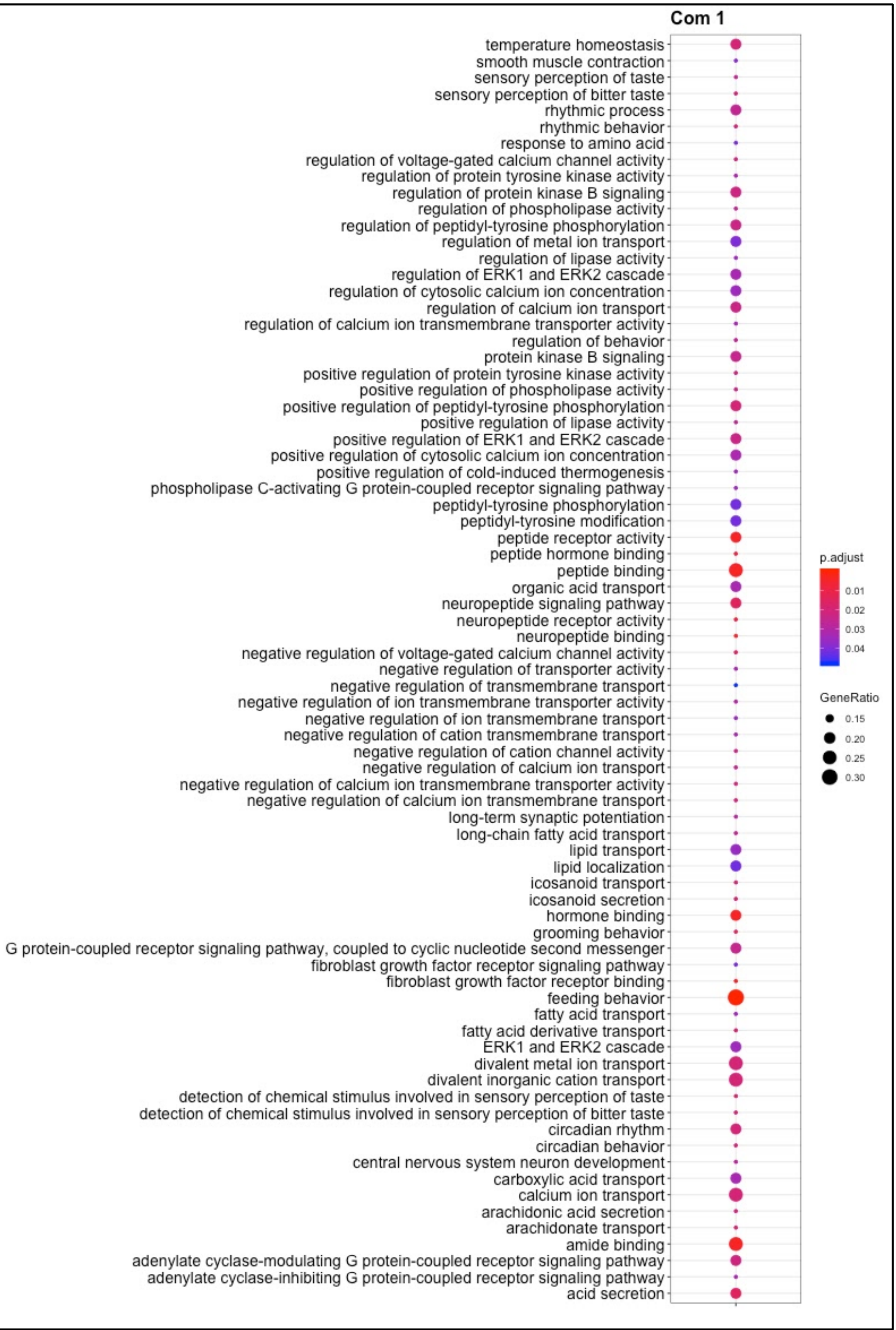

***Figure S15 Enriched GO-Terms for community 2***

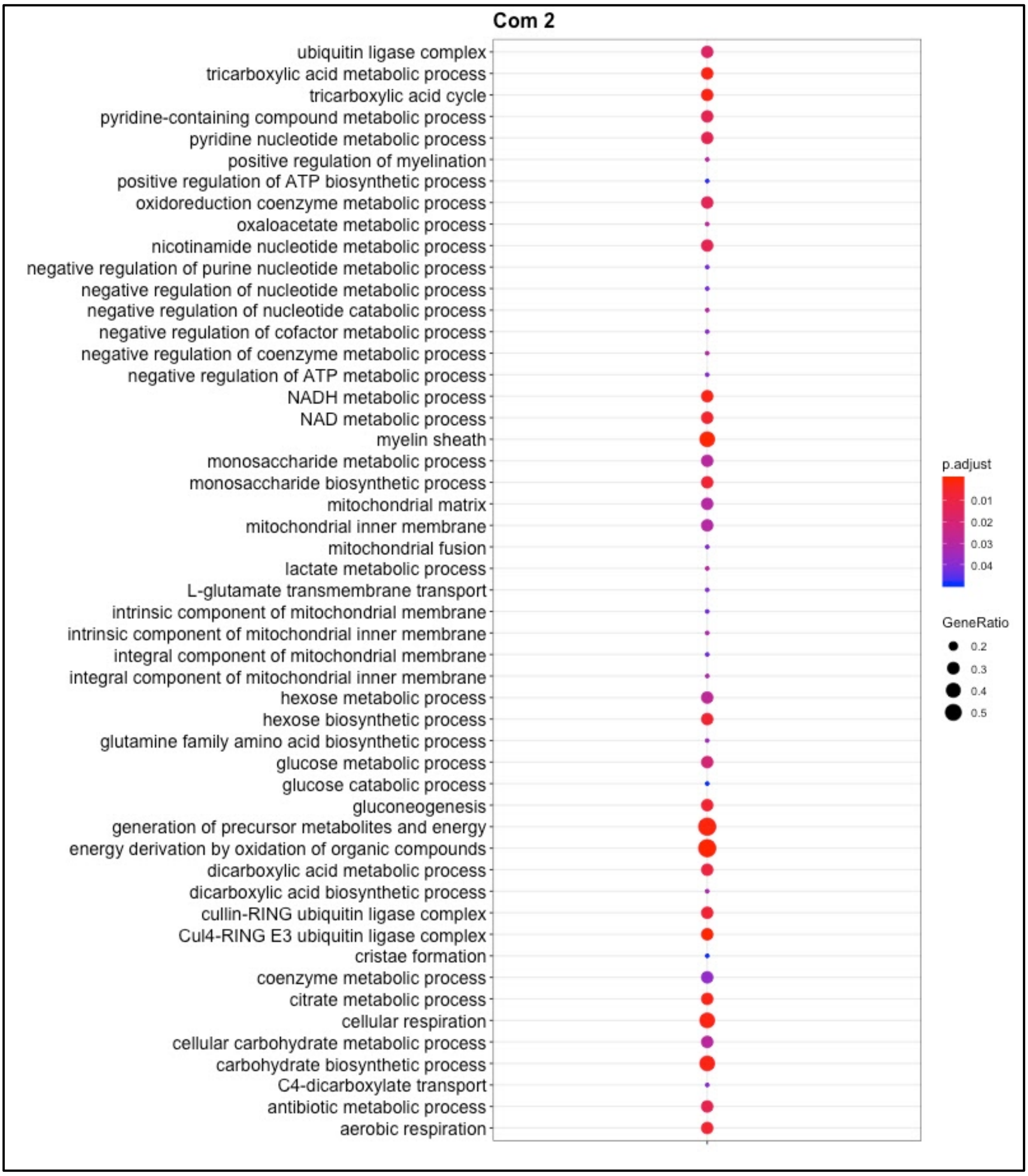

***Figure S16 Enriched GO-Terms for community 3***

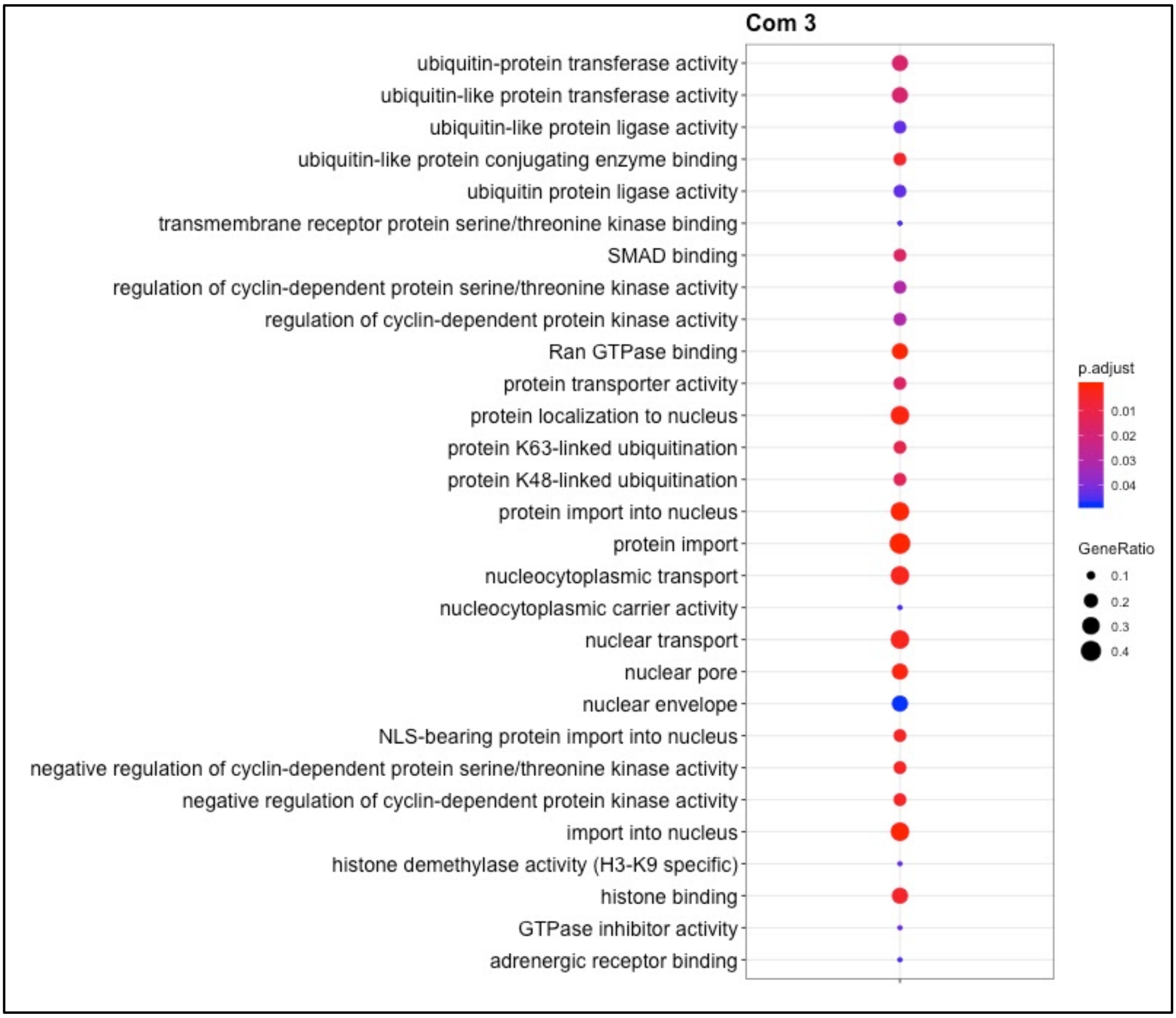

***Figure S17 Enriched GO-Terms for community 4***

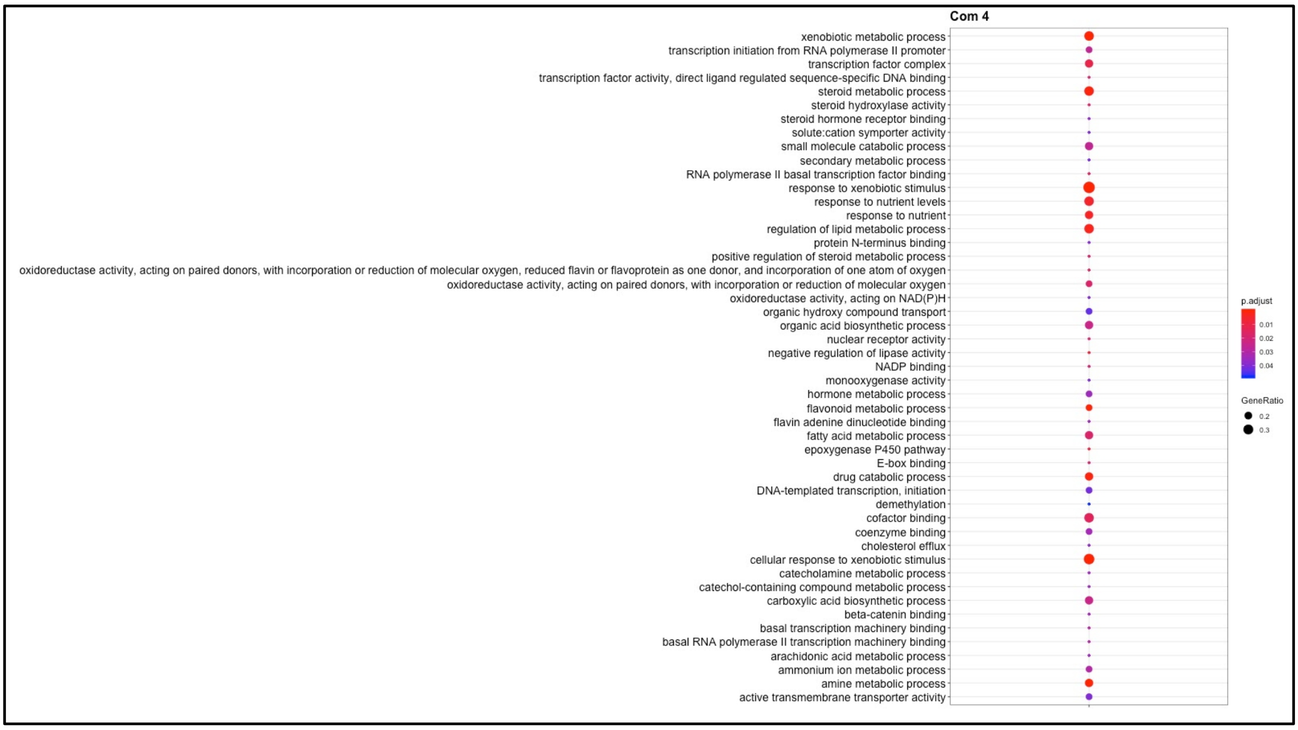


***Figure S18 Enriched GO-Terms for community 5***

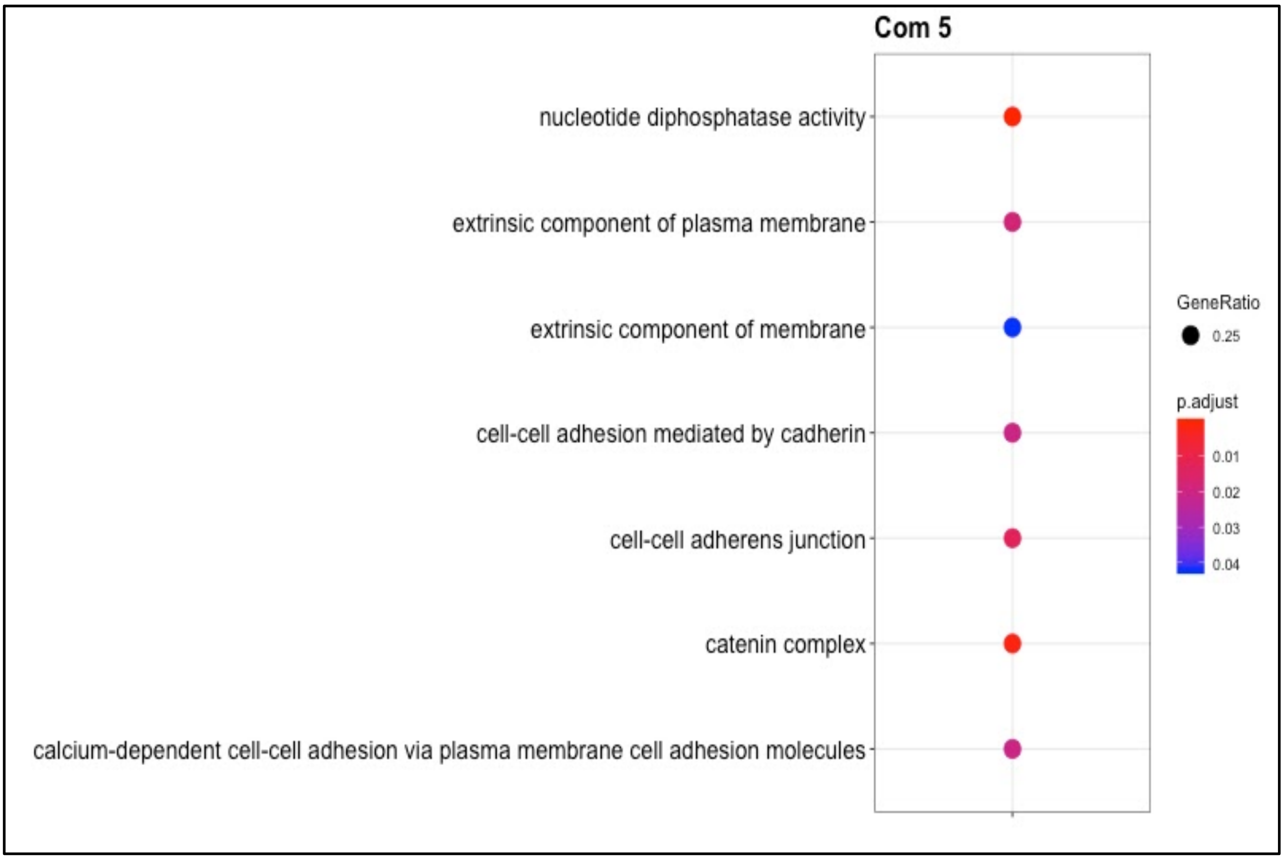

***Figure S19 Enriched GO-Terms for community 6***

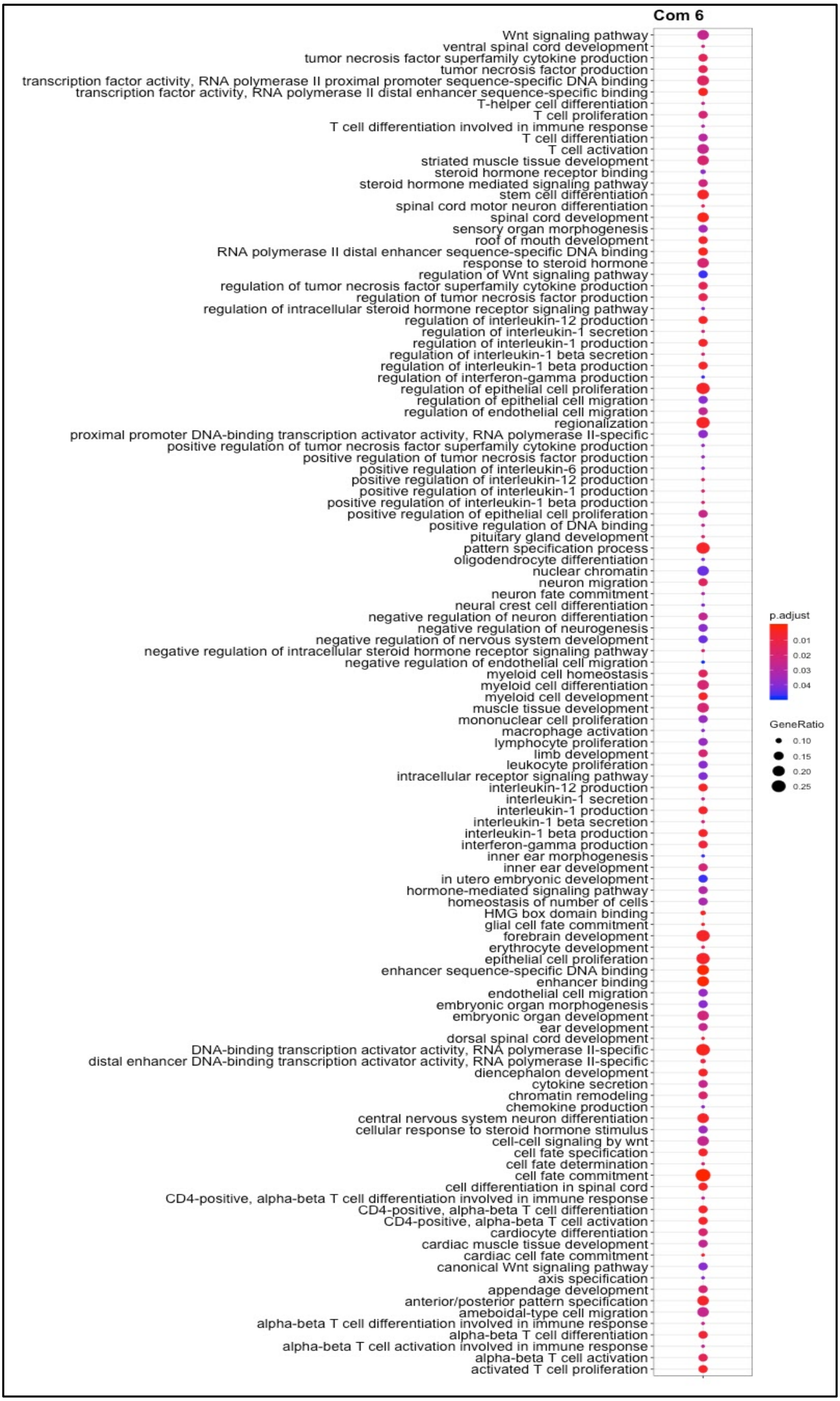

***Figure S20 Enriched GO-Terms for community 7***

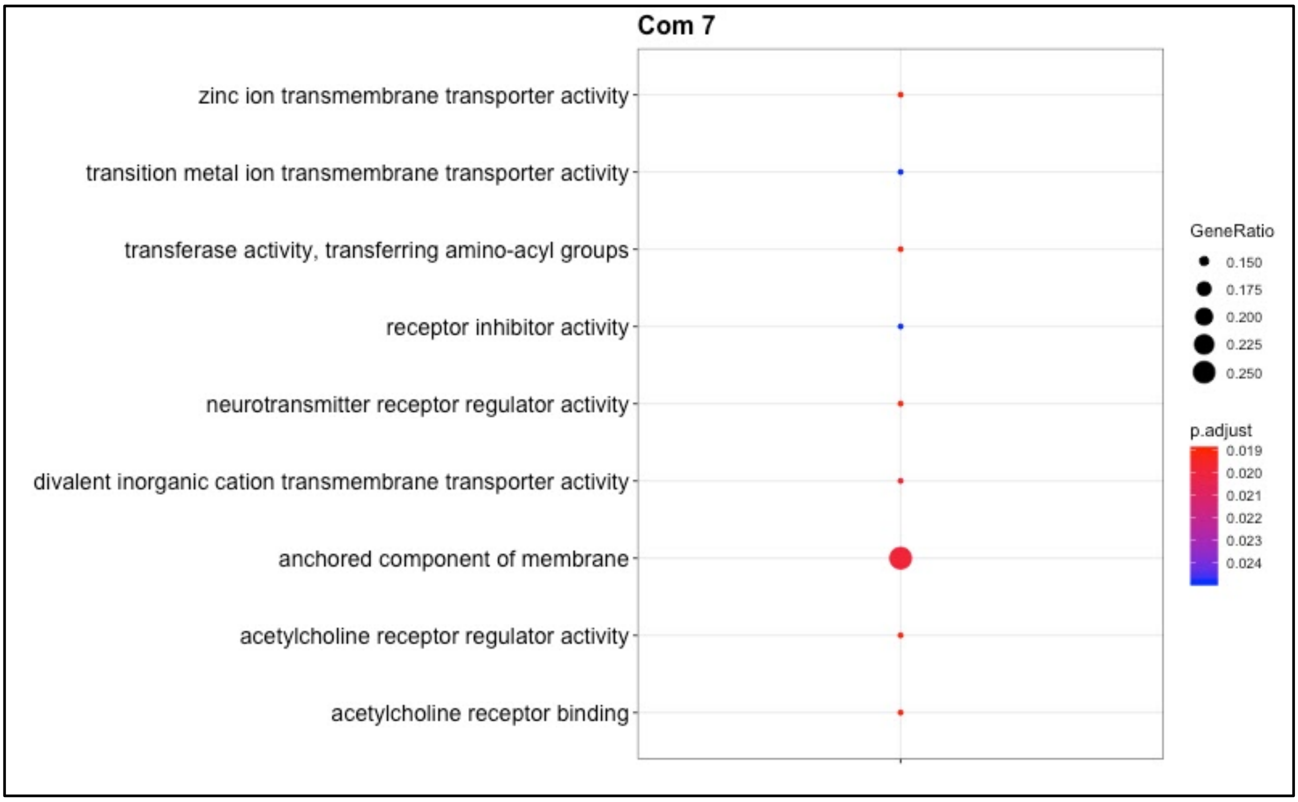

***Figure S21 Enriched GO-Terms for community 8***

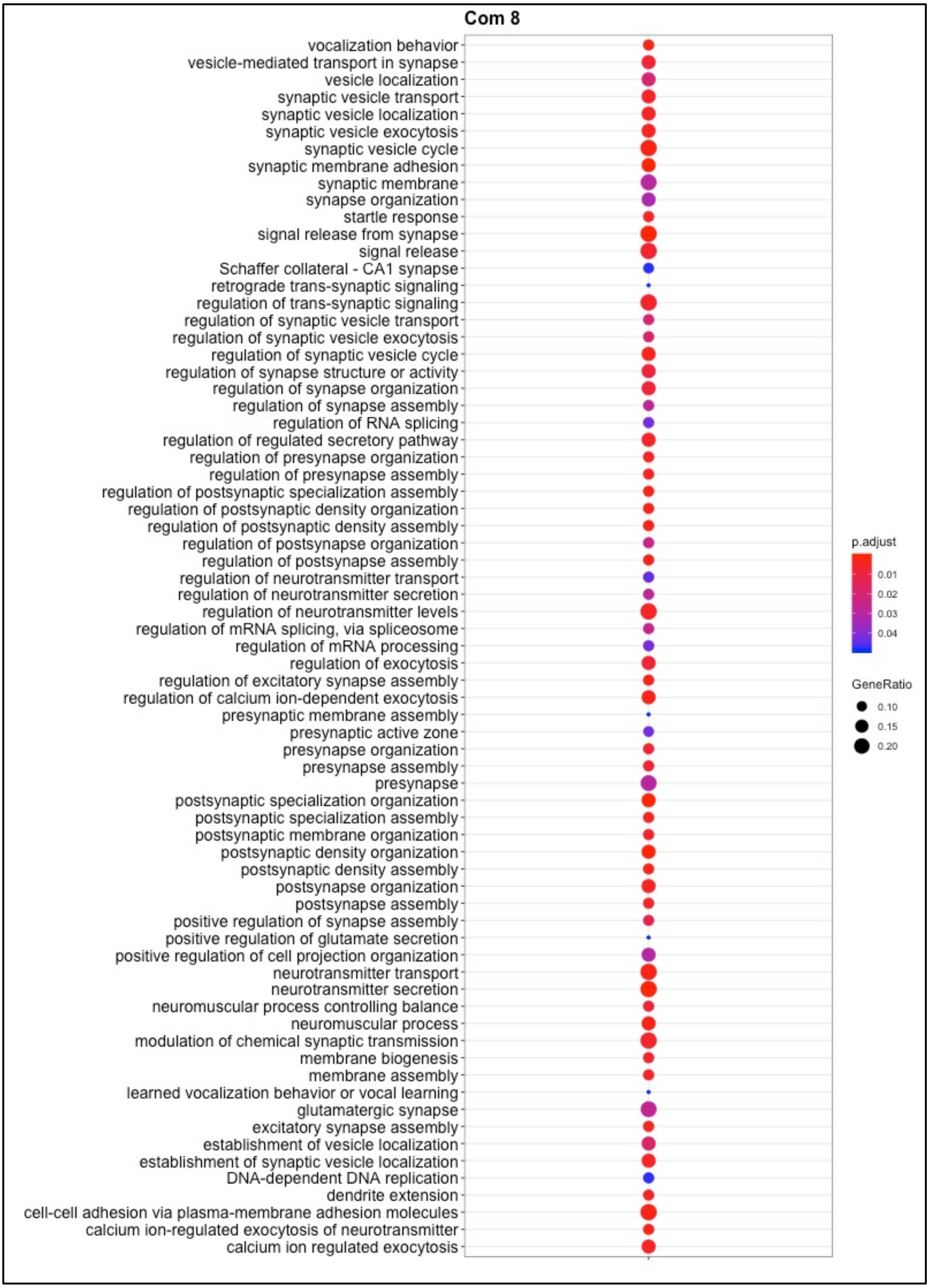

***Figure S22 Enriched GO-Terms for community 9***

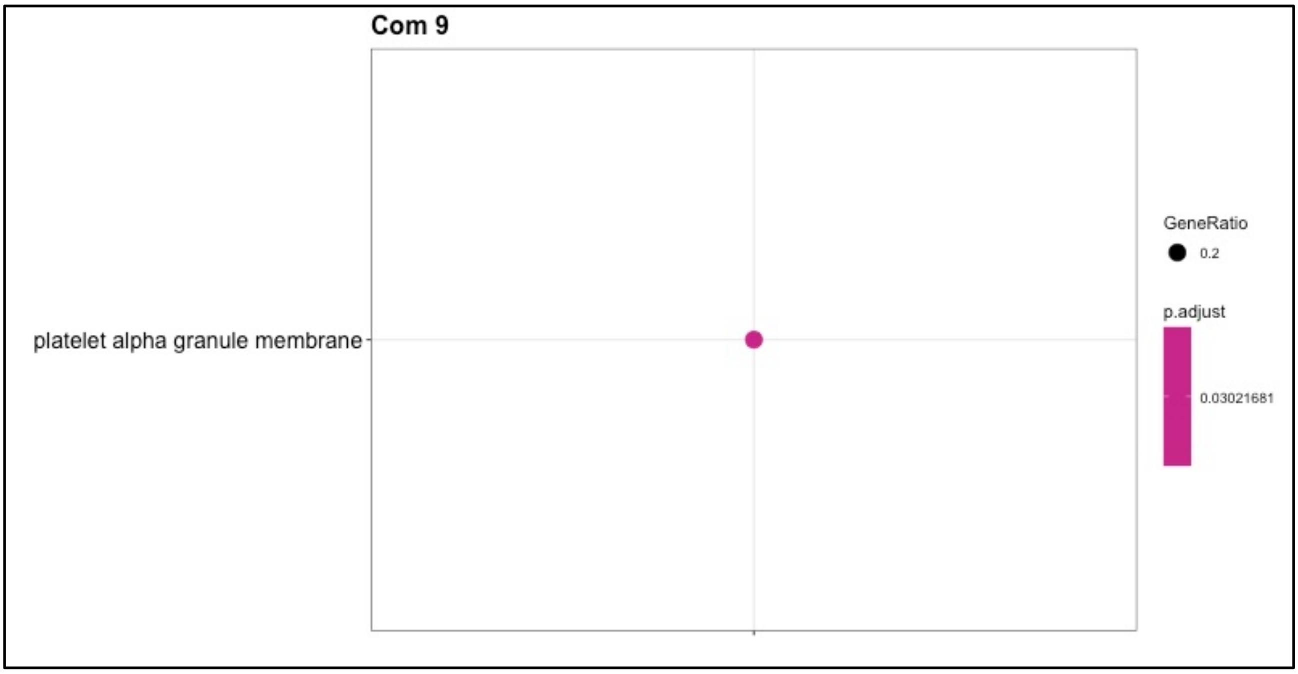

***Figure S23 Enriched GO-Terms for community 10***

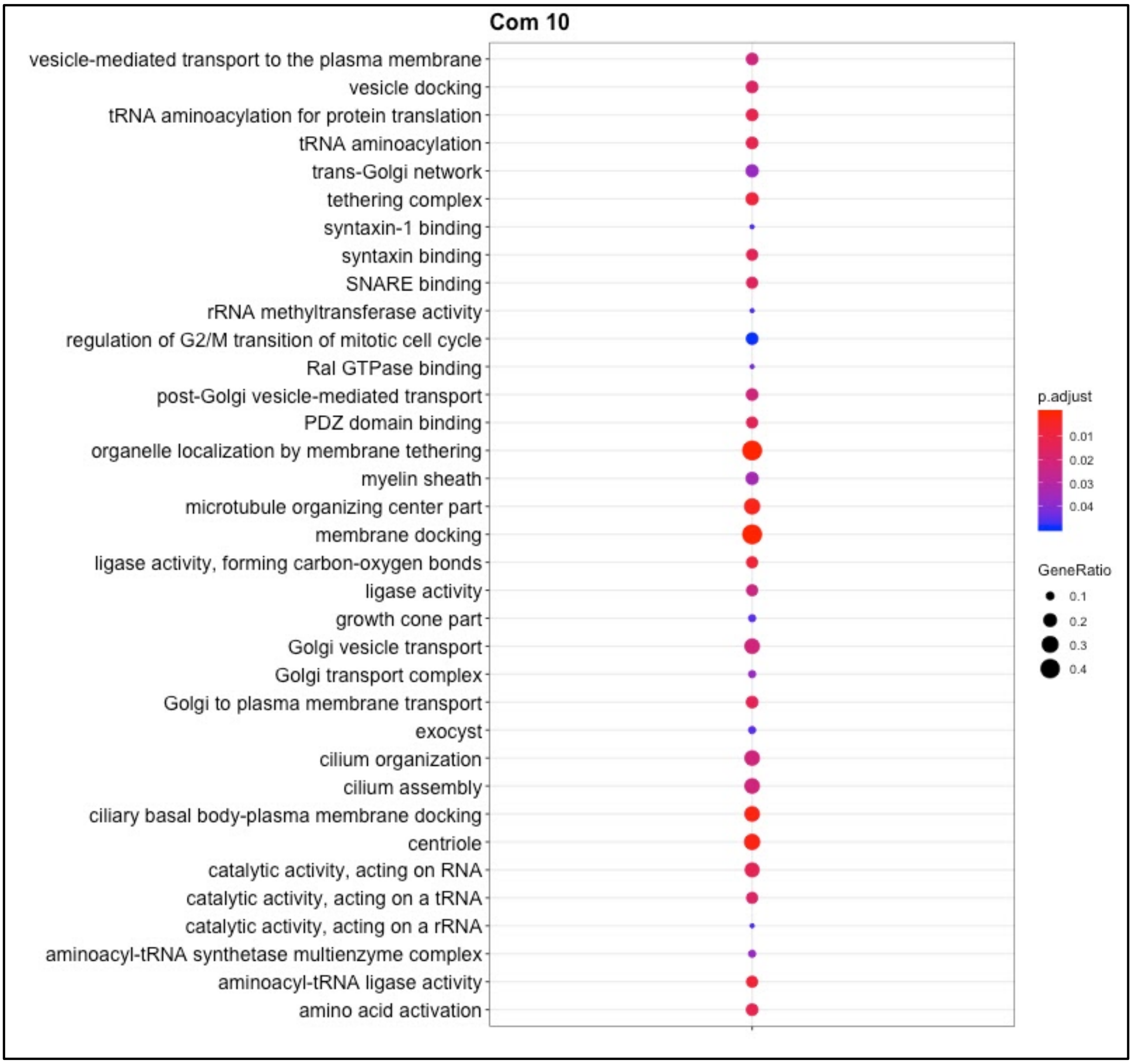

***Figure S24. Selected forest plots of MR-estimated effect sizes. A*** *Forest plot of the effect of Cheese and Meat consumption on lipid and obesity measures. Despite both foods have a high protein and high fat content their effects on lipid levels and BMI are different. Abbreviations BMI Body Mass Index, TRY Triglycerides, TC Total Cholesterol, LDL Low Density Lipoprotein***. B*** *Effect of several foods related to healthy foods on blood tryglicerides levels. Effect for all foods are very similar and make it impossible to distinguish the contribution of each food.*

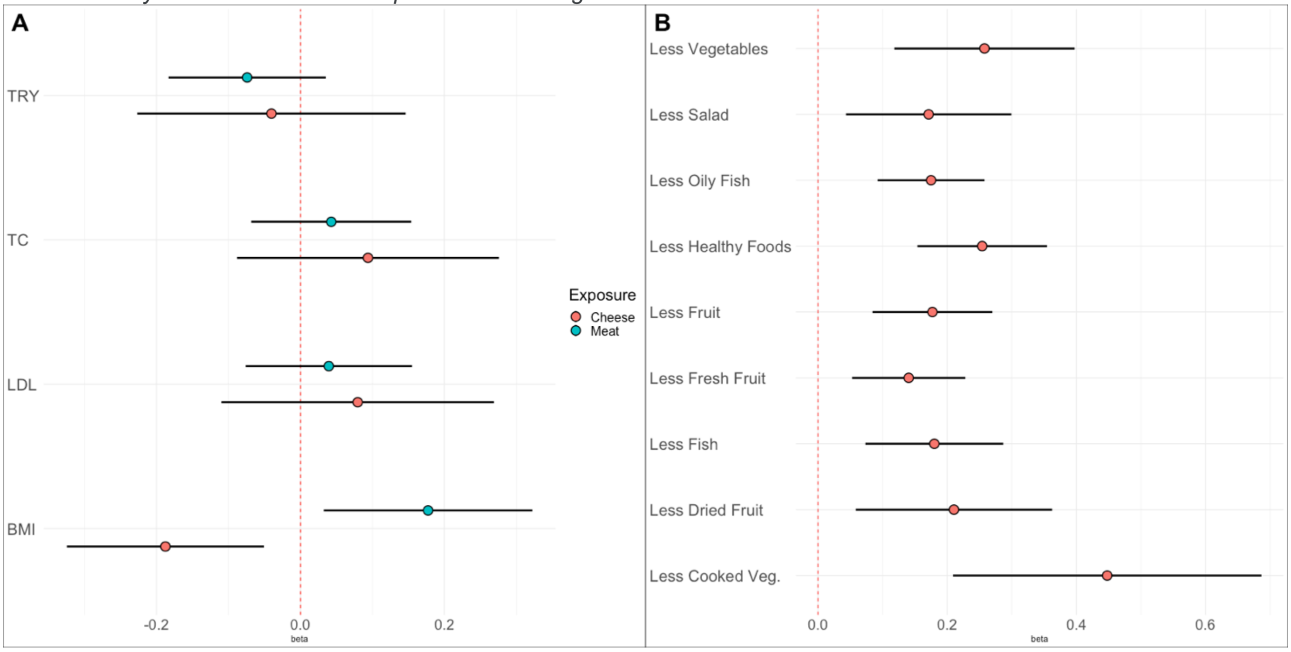

***Figure S25 Effect of food on obesity related measures.*** *The forest plot compares the effect of each food trait on four obesity related measures: BMI, Body Fat, Waist to Hip Ratio (WHR) and BMI adjusted WHR (WHR|BMI). Each color and shape represents a different obesity related measure.*

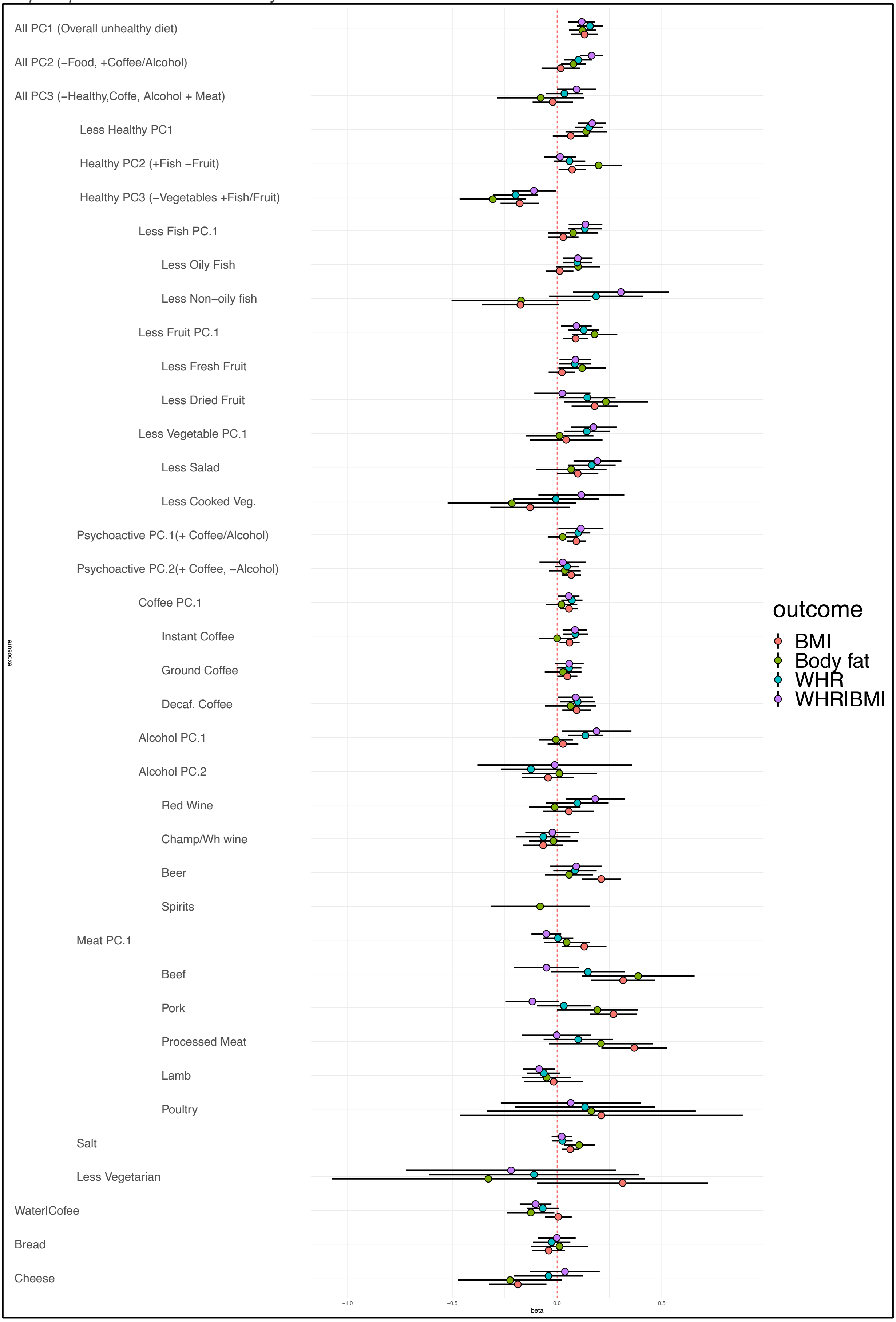

**2.6 Significant results from the uncorrected analysis.**

In order to understand what would have been the impact of performing MR without using CUR for filtering the IVs, we estimated Storey q-values using only the p-values coming from the uncorrected set of results. The following forest plots refer to these results and compare the uncorrected results with the CUR filtered IVs using the uncorrected betas.

Of the 216 significant exposure/outcome pairs which resulted significant after multiple test correction only 91 were in common with those significant at the CUR filtered analysis. In many cases this is clearly due to an overestimation of the effect size. This is particularly evident when looking at Cheese which seems to have a large number of beneficial effects (7) which all disappear (apart from BMI which is diminished in any case) after selecting only the IVs with non-mediated effects. This is unsurprising given that Cheese was the food that had the largest proportion of genetic variance explained by the health related traits (~40%).

These results show how risky it is to make causal claims based on the naive analysis. In our case we could have used these results to make claims of a huge number of beneficial effects of Cheese which do seem to be true and are likely due to the fact that people who consume a larger amount of cheese have a higher education and lower cholesterol which is thus creating the confounding effect. This is of particular importance as MR is generally considered (when performed properly) a sturdy and reliable method of testing causal relationships. However we have shown that there can be issues and particular care should be used when human behaviour is involved in the definition of the exposure trait.

A slightly different example is the case of dried fruit were, despite none of the effects are still significant after using the CUR filtered IVs, comparing the forest plots seems to suggest that this difference is in some cases due to an actual difference in effect size (Years of schooling, Lung Cancer and Ovarian Cancer) while in the rest of the cases the difference is due to a loss in power which has led to an increase in the standard errors of the estimates.

It is unfortunately impossible to perform a direct test of the difference in estimates as the wide confidence intervals of the MR estimates do allow to have enough power to detect the differences. This problem will be overcome in the future through the increase in power due to the increasing size of GWAs studies but at the moment we are not able in many cases to distinguish which associations are not significant any more due to power or difference in effect size. We have however reported the forest plot comparing the two methods and have provided an online tool that each researcher can evaluate all the different estimates coming from different methods and thus make up their own mind based also on external evidence.

***Fig S26-S60 Forest plots of the exposure/outcome pairs significant at the uncorrected analysis.*** *The forest plots represent the estimated effect sizes for all the non CUR filtered MR analyses. The squares represent the point estimates while the bars the 95% confidence intervals. Results from the uncorrected analysis (raw) and CUR filtered IVs (CUR) are reported. The exposure trait is indicated in the header of the plots while the row labels refer to the outcomes. Beta's always refer to standard deviations for the exposure while for the outcomes it is standard deviations for the quantitative traits and log(OR) for the disease traits.*

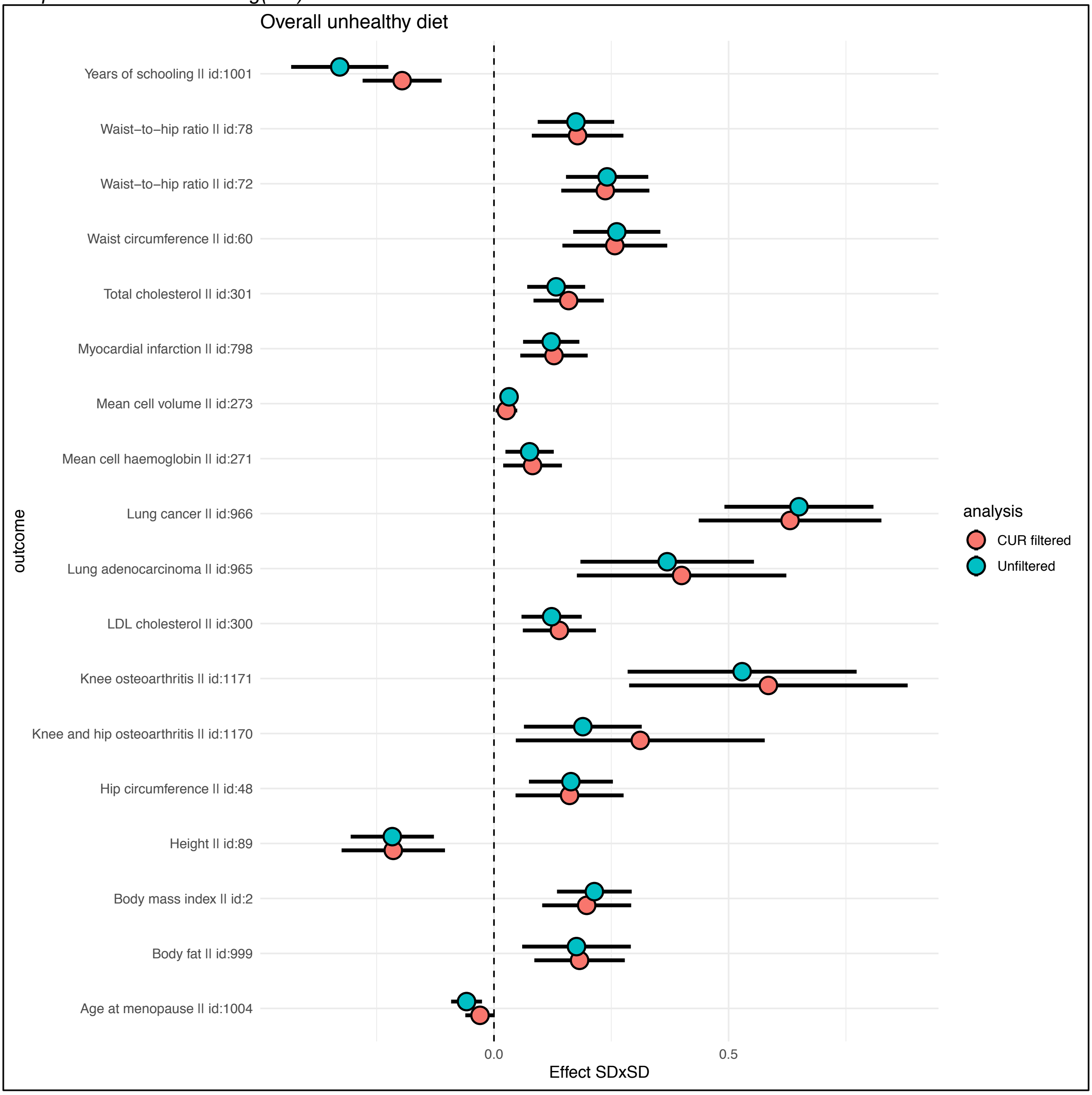

***Fig S27***

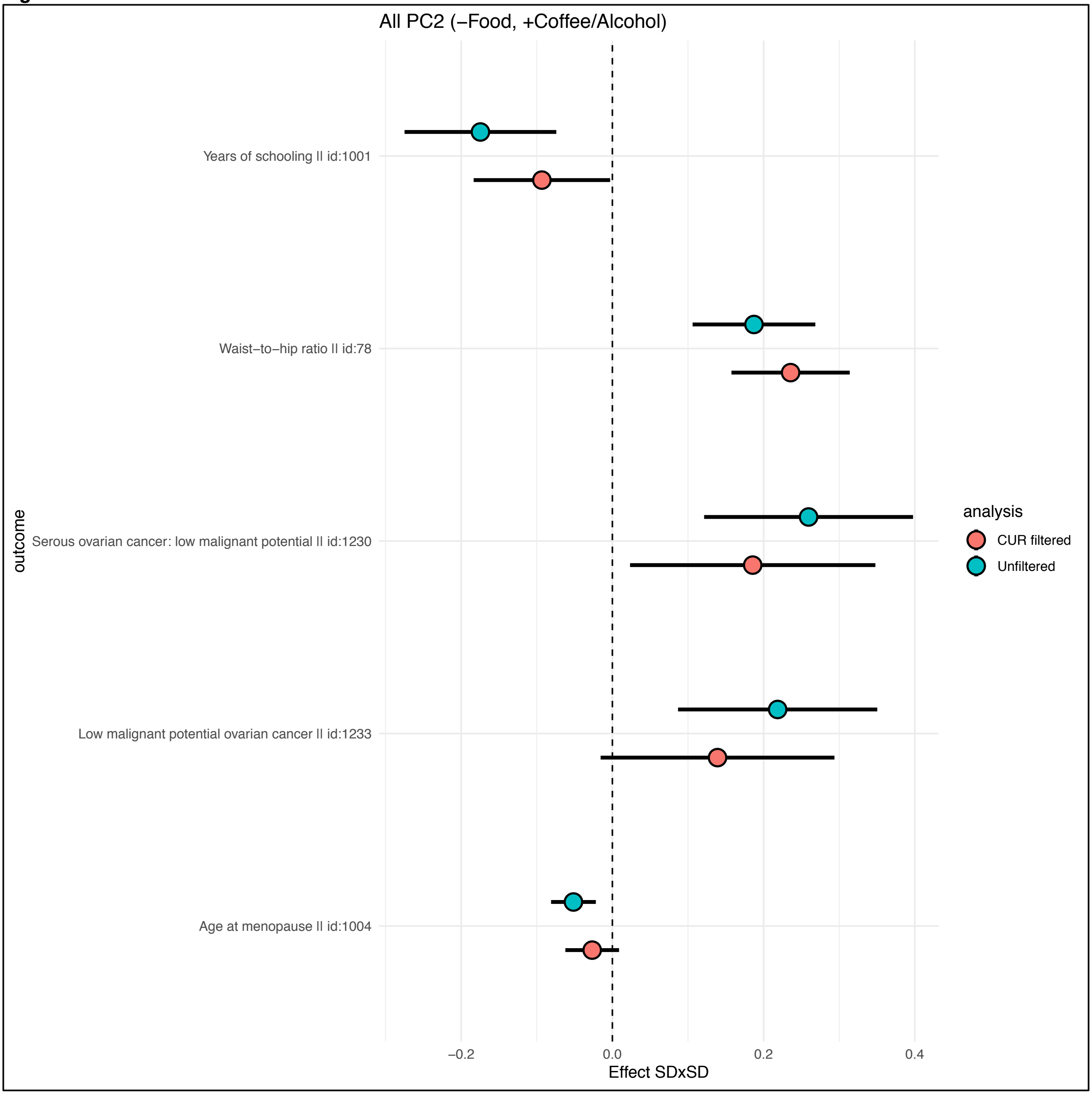
All PC2 (–Food, +Coffee/Alcohol)
Years of schooling || id:1001
Waist–to–hip ratio || id:78
Serous ovarian cancer: low malignant potential || id:1230
Low malignant potential ovarian cancer || id:1233
Age at menopause || id:1004
outcome
–0.2
0.0
0.2
0.4
Effect SDxSD
analysis
CUR filtered
Unfiltered

***Fig S28***

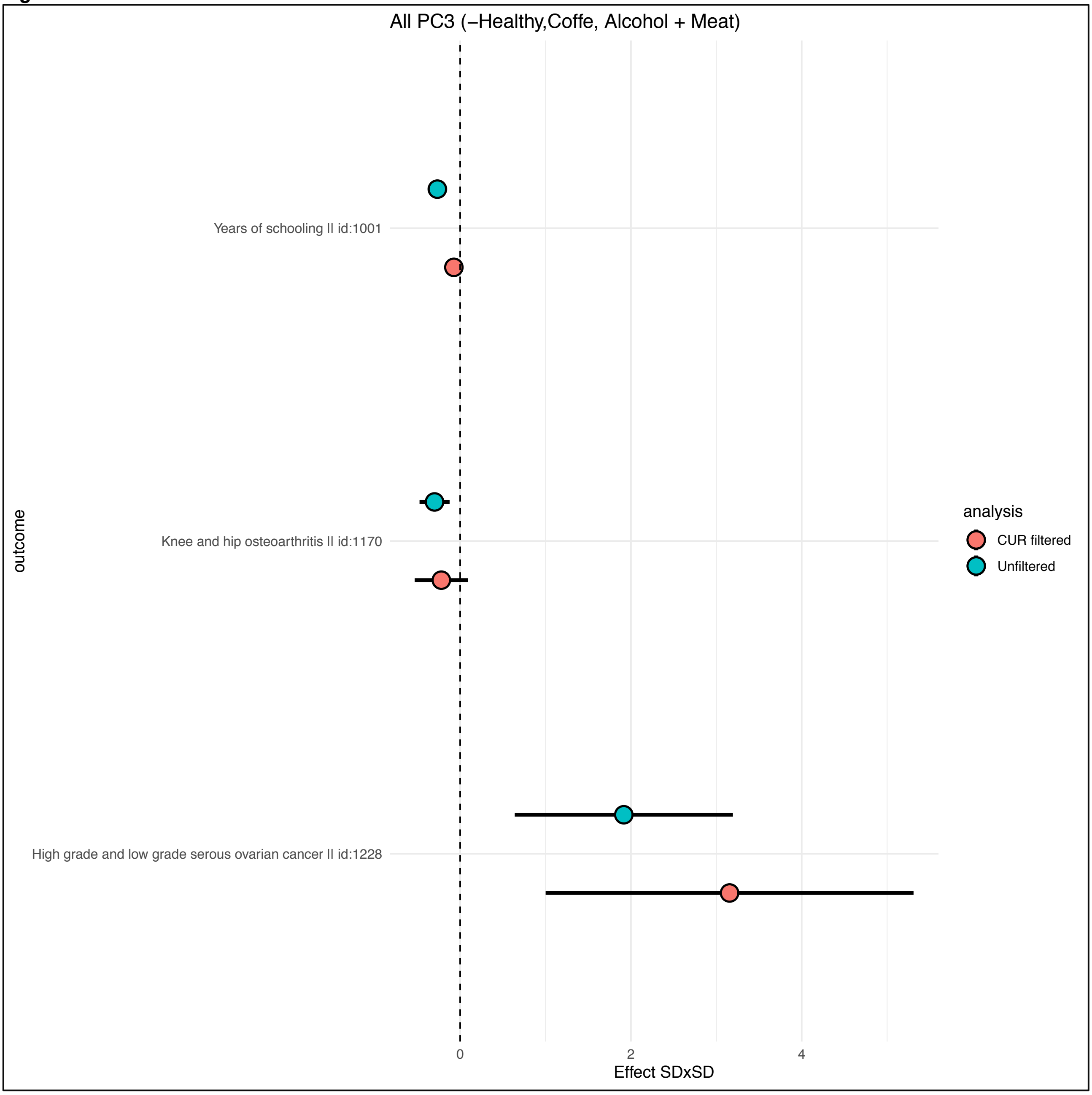
All PC3 (−Healthy,Coffe, Alcohol + Meat)
Years of schooling || id:1001
Knee and hip osteoarthritis || id:1170
High grade and low grade serous ovarian cancer || id:1228
outcome
0
2
4
Effect SDxSD
analysis
CUR filtered
Unfiltered

***Fig S29***

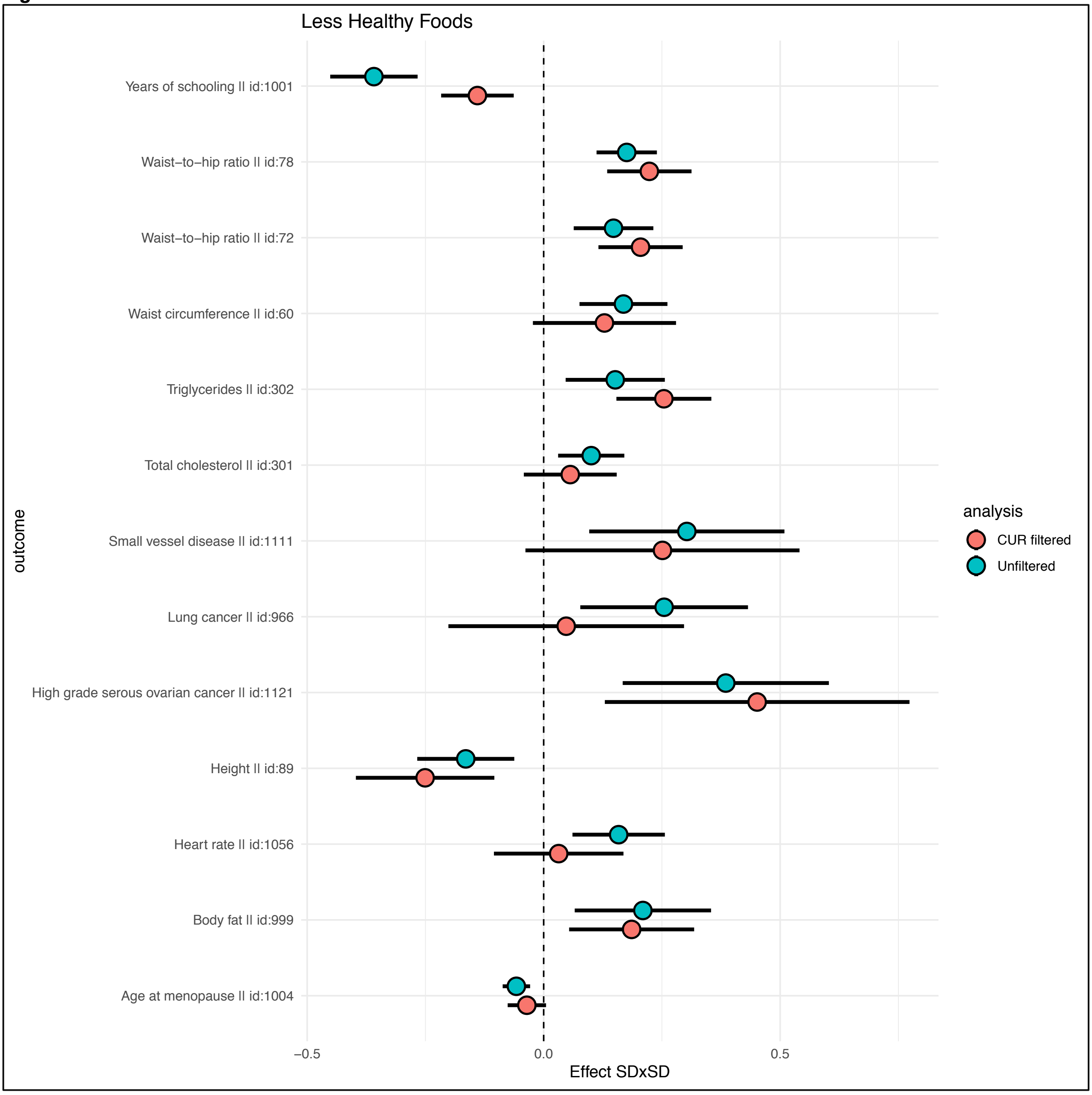
Less Healthy Foods
Years of schooling || id:1001
Waist−to−hip ratio || id:78
Waist−to−hip ratio || id:72
Waist circumference || id:60
Triglycerides || id:302
Total cholesterol || id:301
Small vessel disease || id:1111
Lung cancer || id:966
High grade serous ovarian cancer || id:1121
Height || id:89
Heart rate || id:1056
Body fat || id:999
Age at menopause || id:1004
outcome
−0.5
0.0
0.5
Effect SDxSD
analysis
CUR filtered
Unfiltered

***Fig S30***

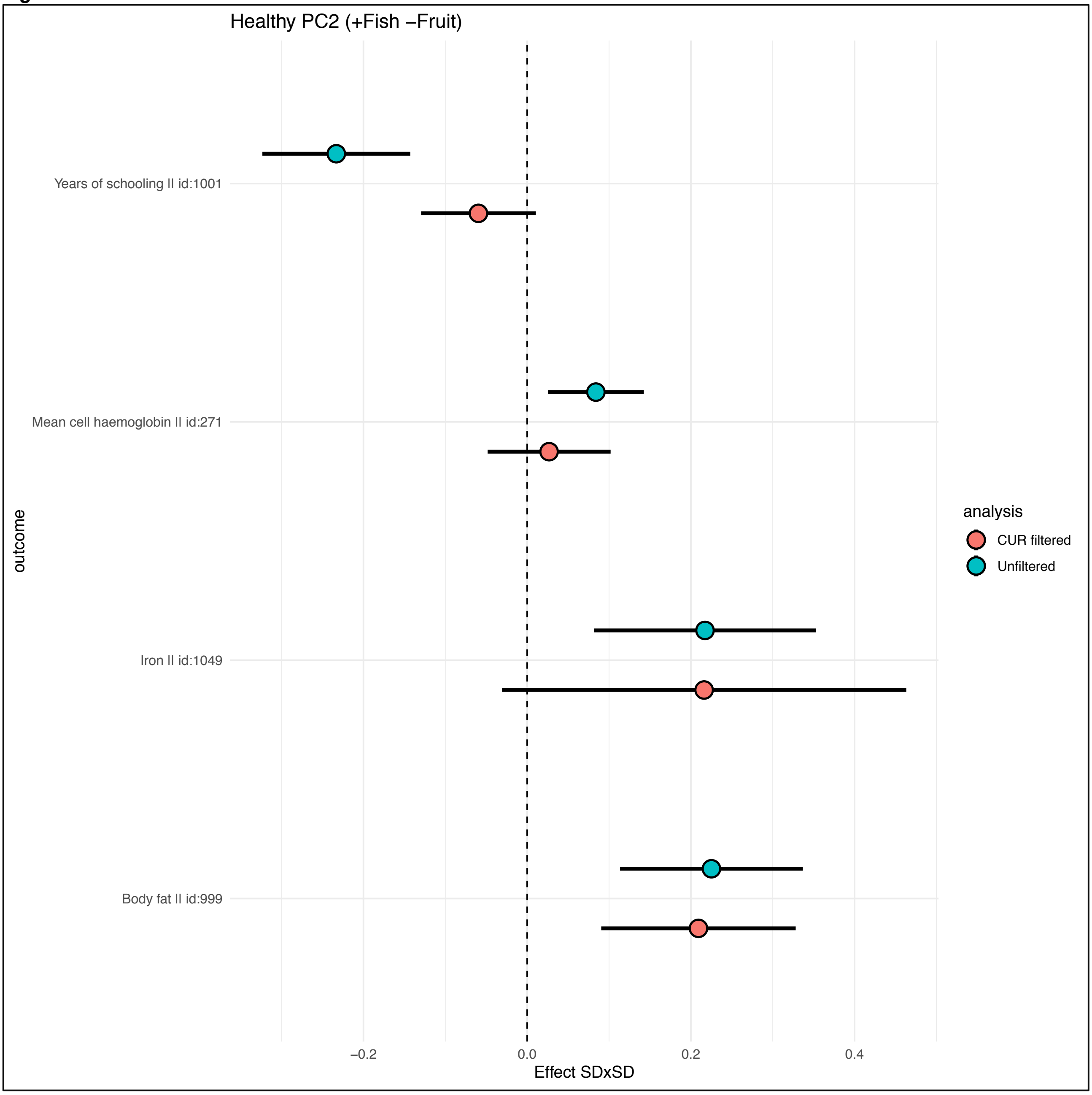

Healthy PC2 (+Fish −Fruit)
Years of schooling || id:1001
Mean cell haemoglobin || id:271
Iron || id:1049
Body fat || id:999
outcome
−0.2
0.0
0.2
0.4
Effect SDxSD
analysis
CUR filtered
Unfiltered

707 ***Fig S31***

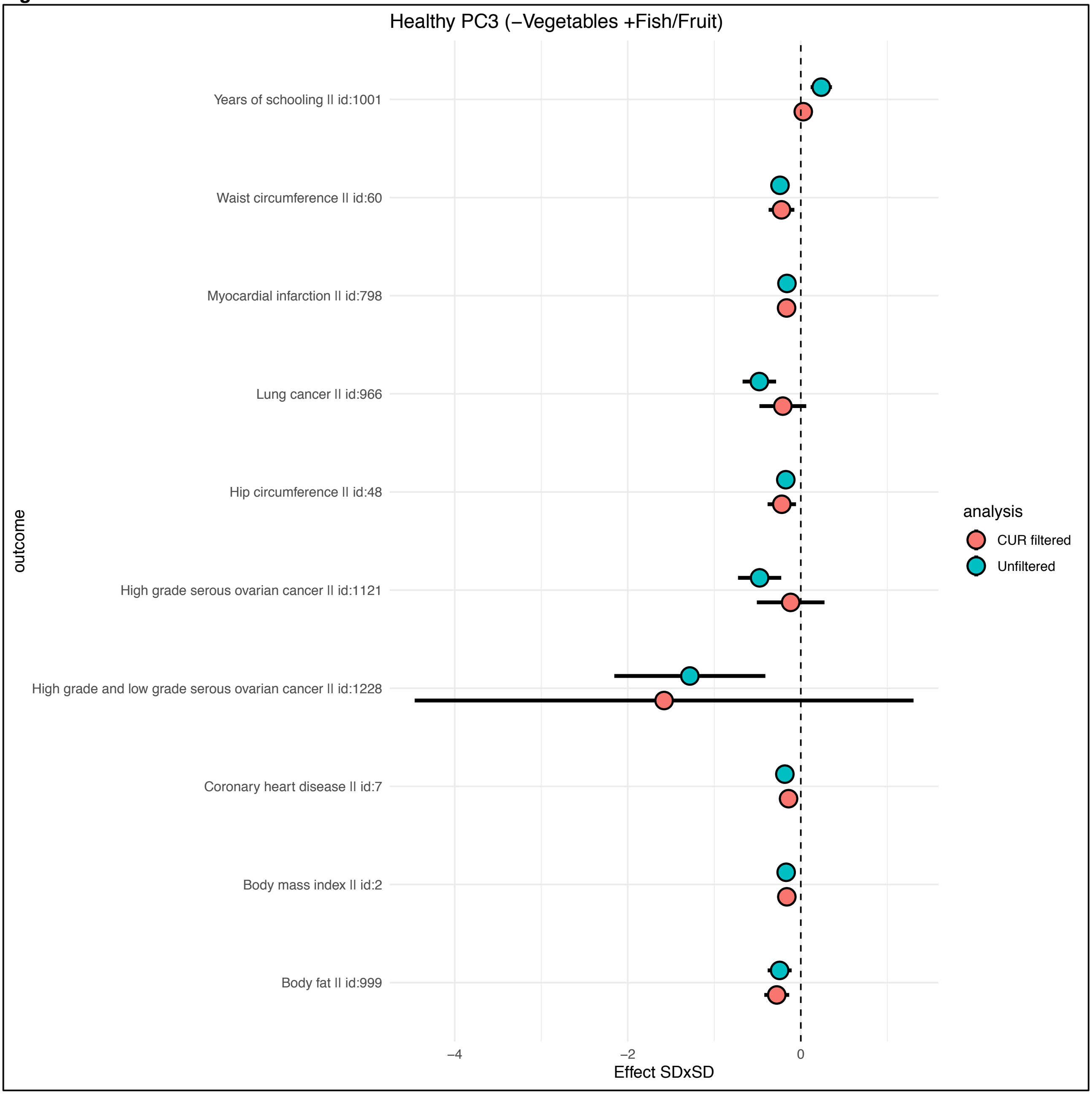


708

***Fig S32***

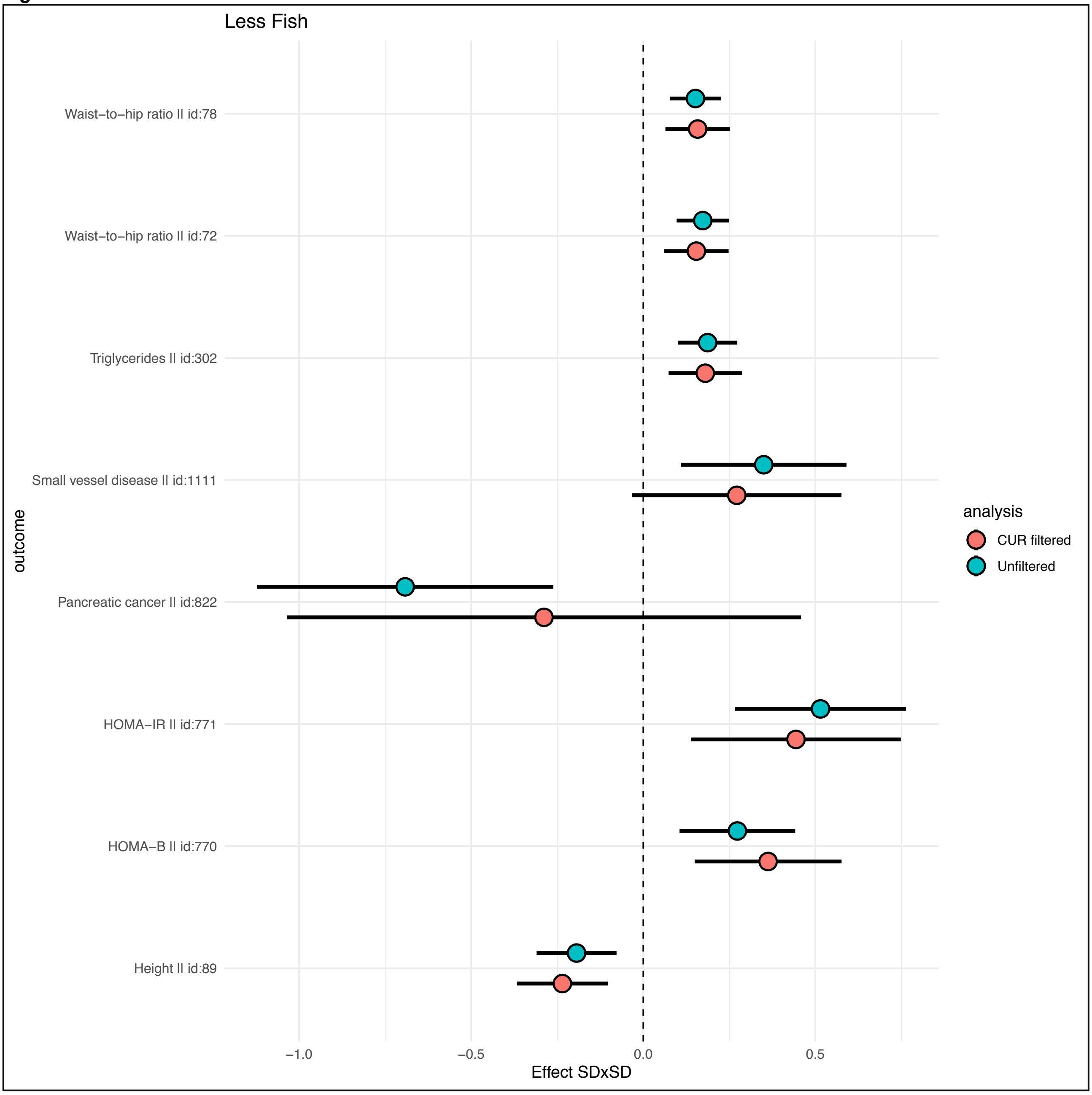
Less Fish
Waist−to−hip ratio || id:78
Waist−to−hip ratio || id:72
Triglycerides || id:302
Small vessel disease || id:1111
Pancreatic cancer || id:822
HOMA−IR || id:771
HOMA−B || id:770
Height || id:89
outcome
−1.0
−0.5
0.0
0.5
Effect SDxSD
analysis
CUR filtered
Unfiltered

***Fig S33***

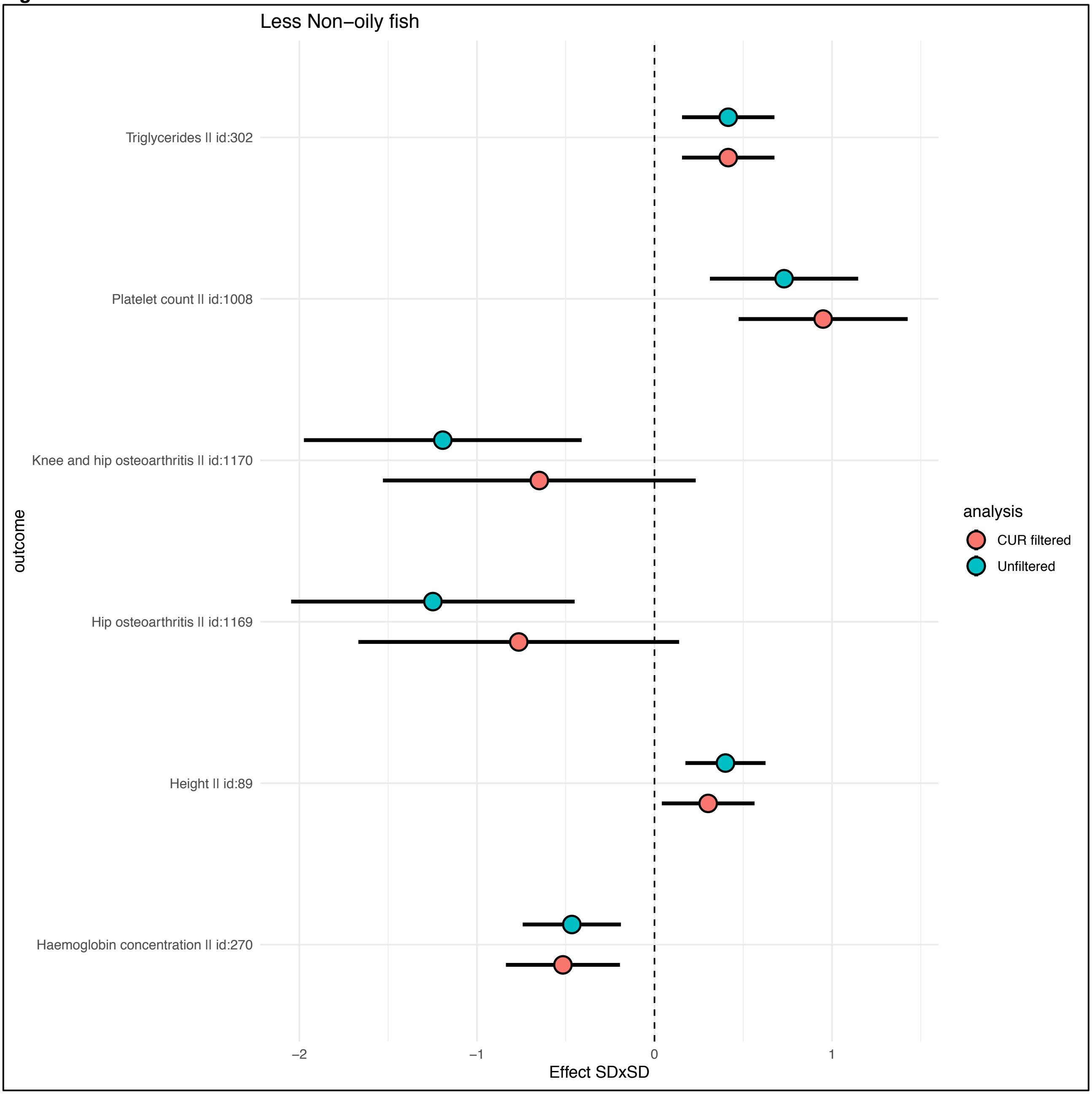
Less Non−oily fish
Triglycerides || id:302
Platelet count || id:1008
Knee and hip osteoarthritis || id:1170
Hip osteoarthritis || id:1169
Height || id:89
Haemoglobin concentration || id:270
outcome
−2
−1
0
1
Effect SDxSD
analysis
CUR filtered
Unfiltered

***Fig S34***

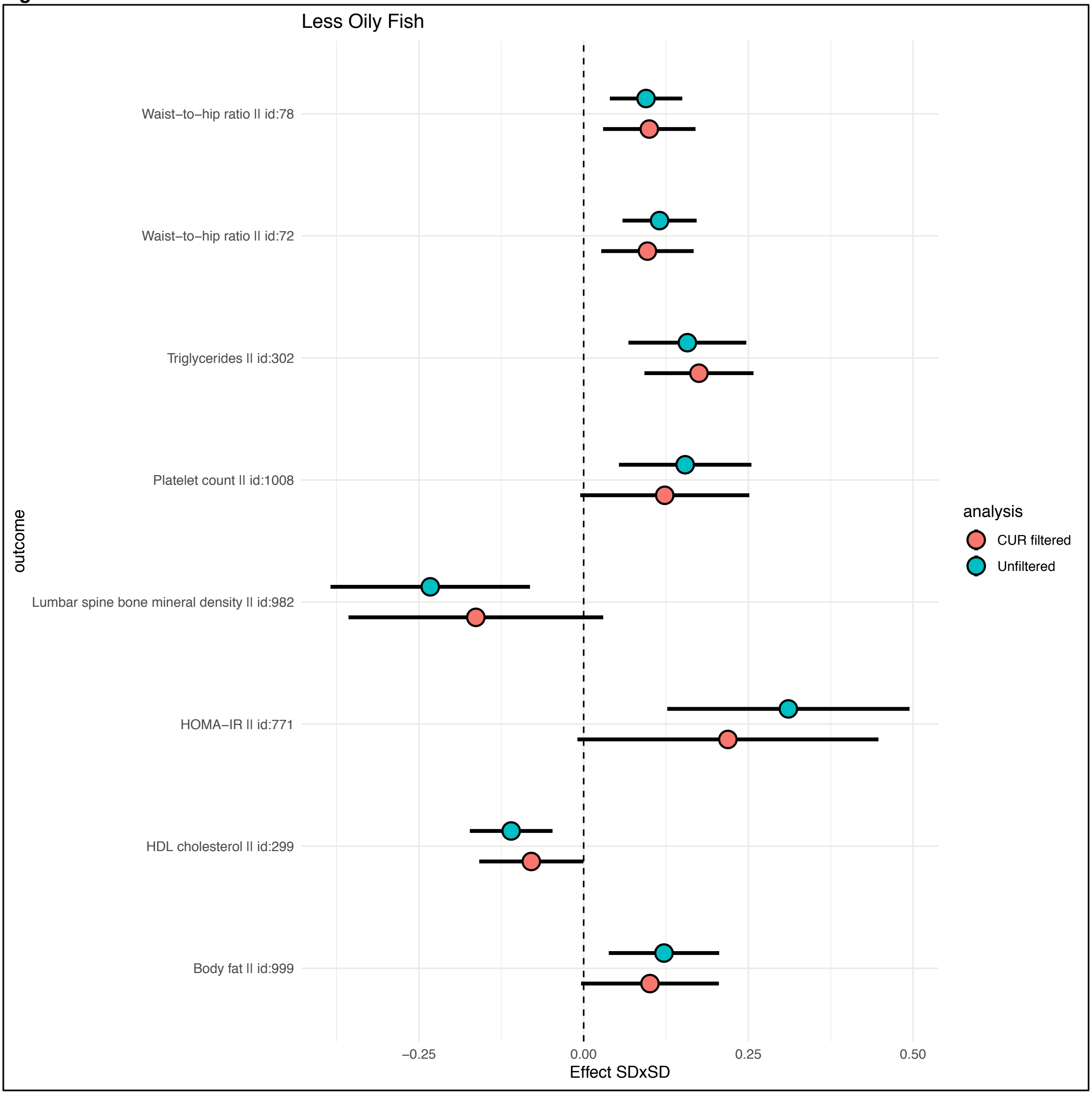
Less Oily Fish
Waist−to−hip ratio || id:78
Waist−to−hip ratio || id:72
Triglycerides || id:302
Platelet count || id:1008
Lumbar spine bone mineral density || id:982
HOMA−IR || id:771
HDL cholesterol || id:299
Body fat || id:999
outcome
−0.25
0.00
0.25
0.50
Effect SDxSD
analysis
CUR filtered
Unfiltered

***Fig S35***

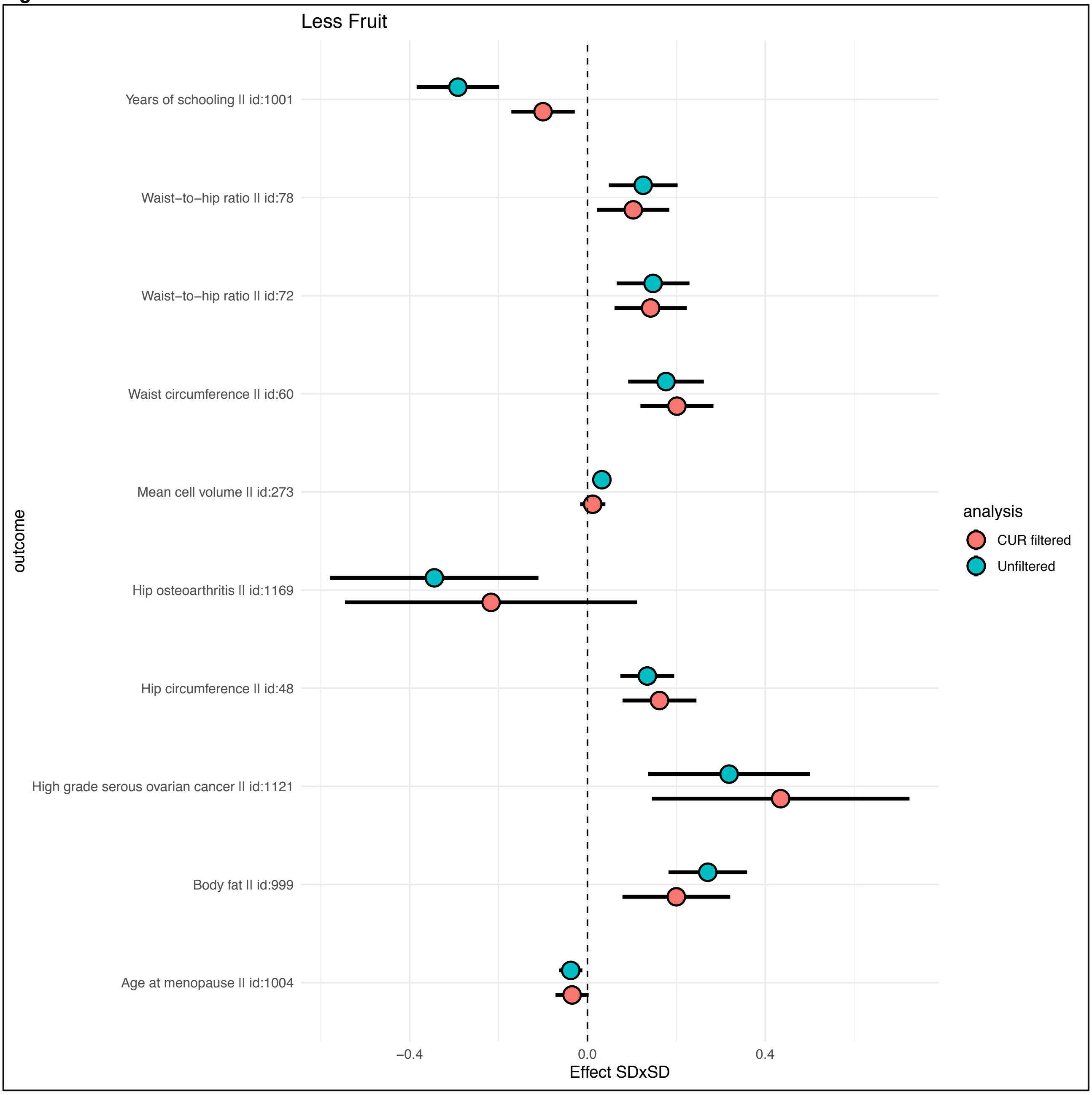

Less Fruit
Years of schooling || id:1001
Waist−to−hip ratio || id:78
Waist−to−hip ratio || id:72
Waist circumference || id:60
Mean cell volume || id:273
Hip osteoarthritis || id:1169
Hip circumference || id:48
High grade serous ovarian cancer || id:1121
Body fat || id:999
Age at menopause || id:1004
outcome
−0.4
0.0
0.4
Effect SDxSD
analysis
CUR filtered
Unfiltered

***Fig S36***

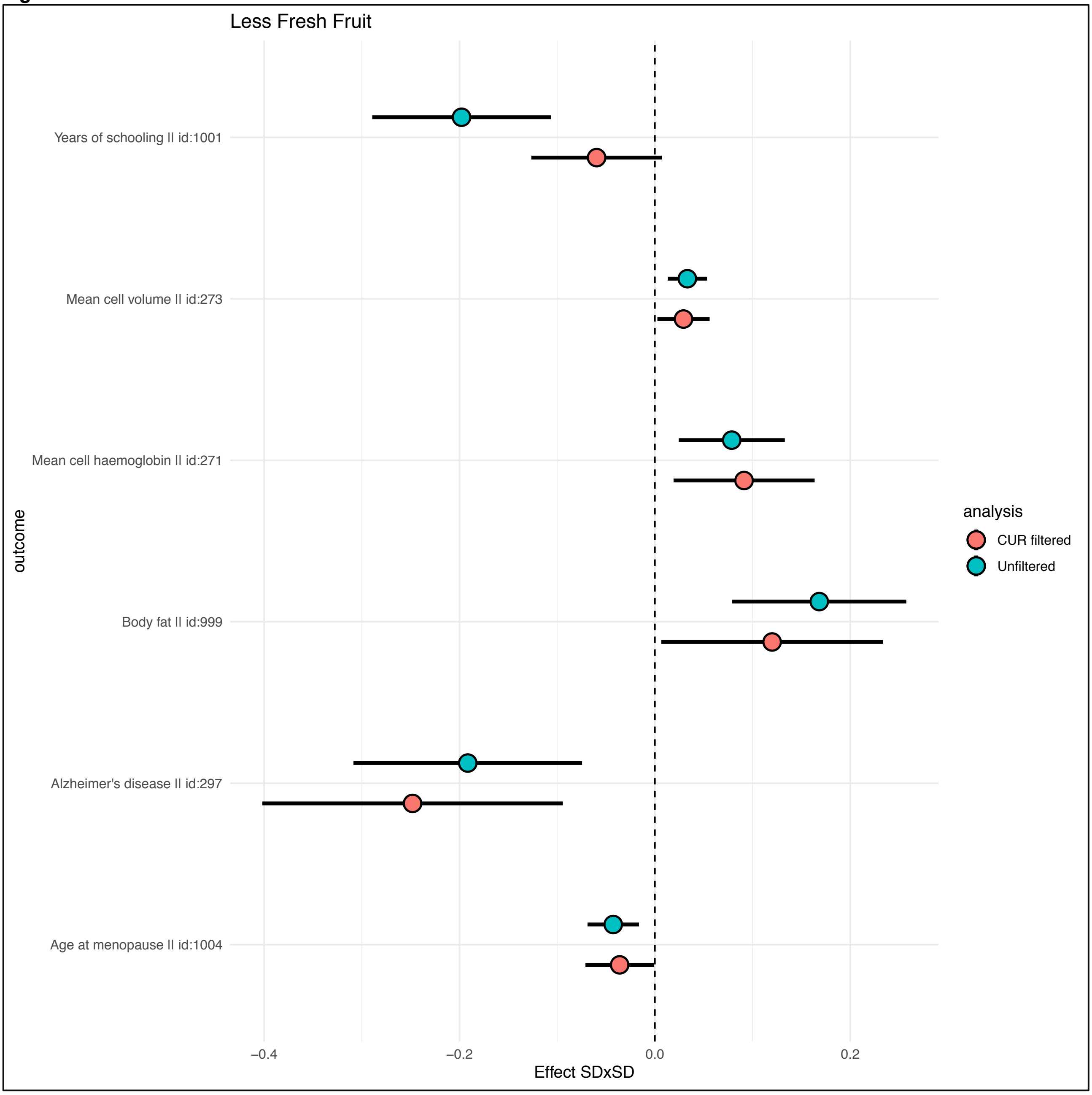
Less Fresh Fruit
Years of schooling II id:1001
Mean cell volume II id:273
Mean cell haemoglobin II id:271
Body fat II id:999
Alzheimer's disease II id:297
Age at menopause II id:1004
outcome
−0.4
−0.2
0.0
0.2
Effect SDxSD
analysis
CUR filtered
Unfiltered

*Fig S37*

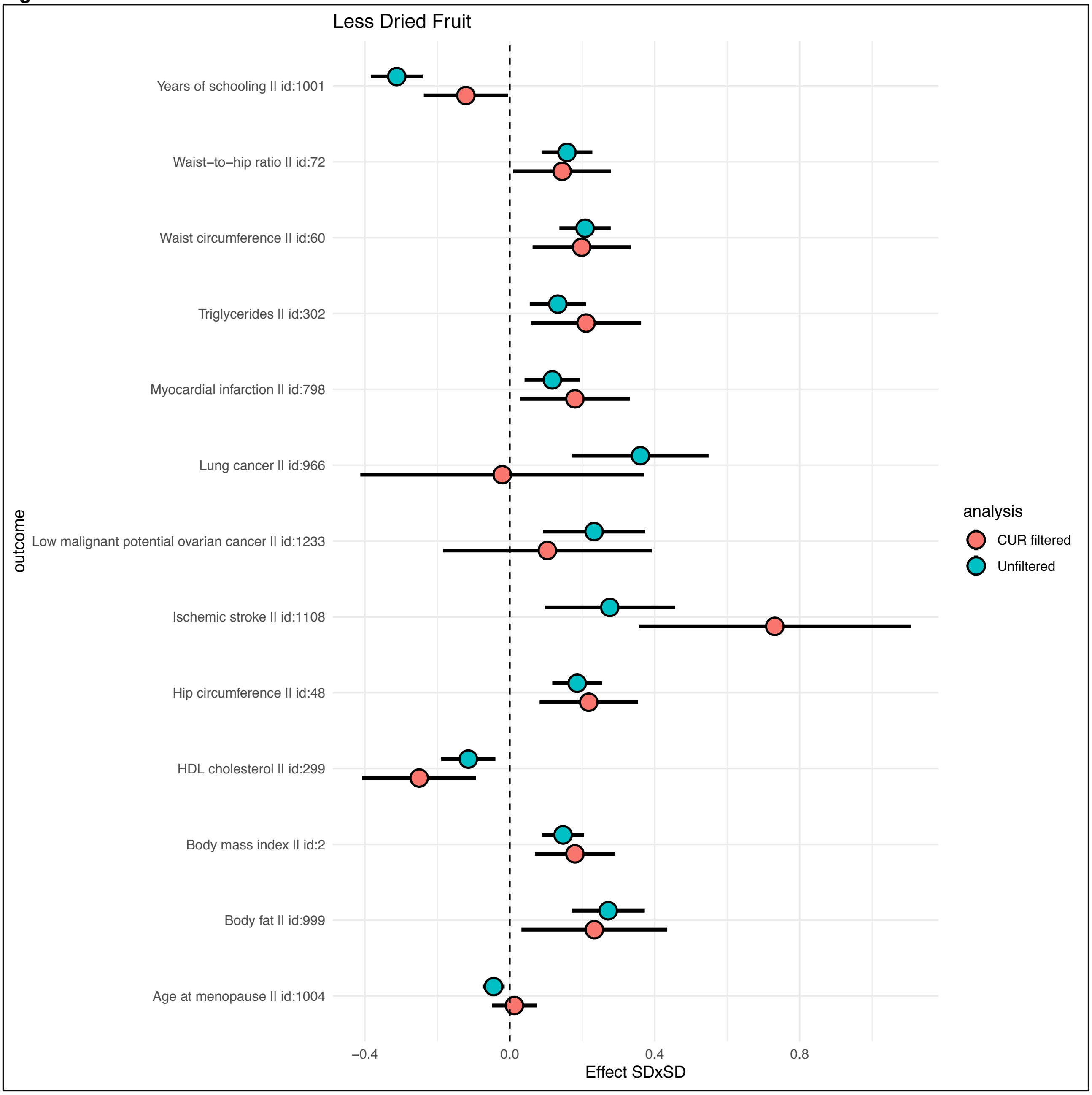

Less Dried Fruit
Years of schooling || id:1001
Waist−to−hip ratio || id:72
Waist circumference || id:60
Triglycerides || id:302
Myocardial infarction || id:798
Lung cancer || id:966
Low malignant potential ovarian cancer || id:1233
Ischemic stroke || id:1108
Hip circumference || id:48
HDL cholesterol || id:299
Body mass index || id:2
Body fat || id:999
Age at menopause || id:1004
outcome
−0.4
0.0
0.4
0.8
Effect SDxSD
analysis
CUR filtered
Unfiltered

***Fig S38***

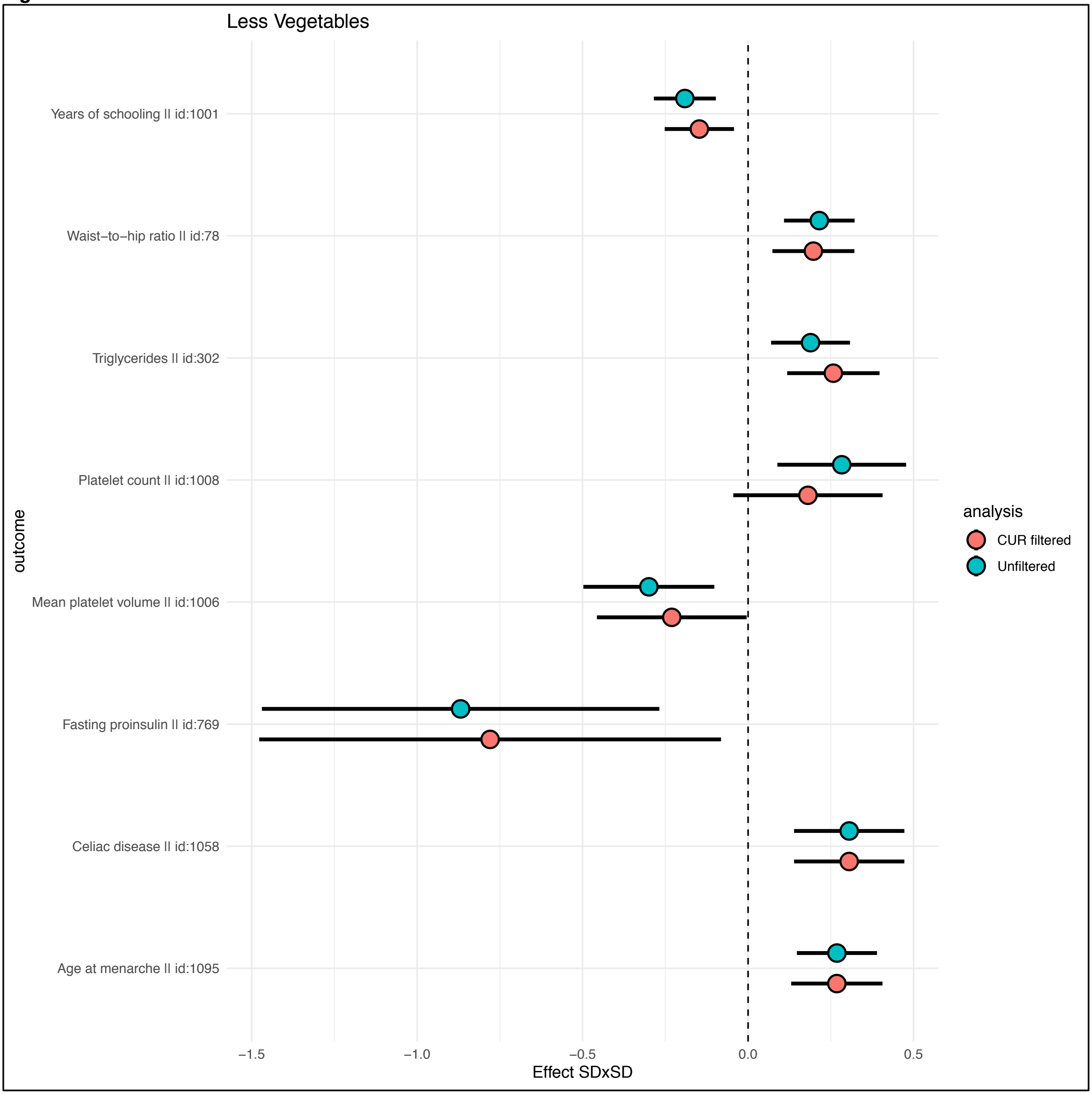
Less Vegetables
Years of schooling || id:1001
Waist–to–hip ratio || id:78
Triglycerides || id:302
Platelet count || id:1008
Mean platelet volume || id:1006
Fasting proinsulin || id:769
Celiac disease || id:1058
Age at menarche || id:1095
outcome
−1.5
−1.0
−0.5
0.0
0.5
Effect SDxSD
analysis
CUR filtered
Unfiltered

723 *Fig S39*

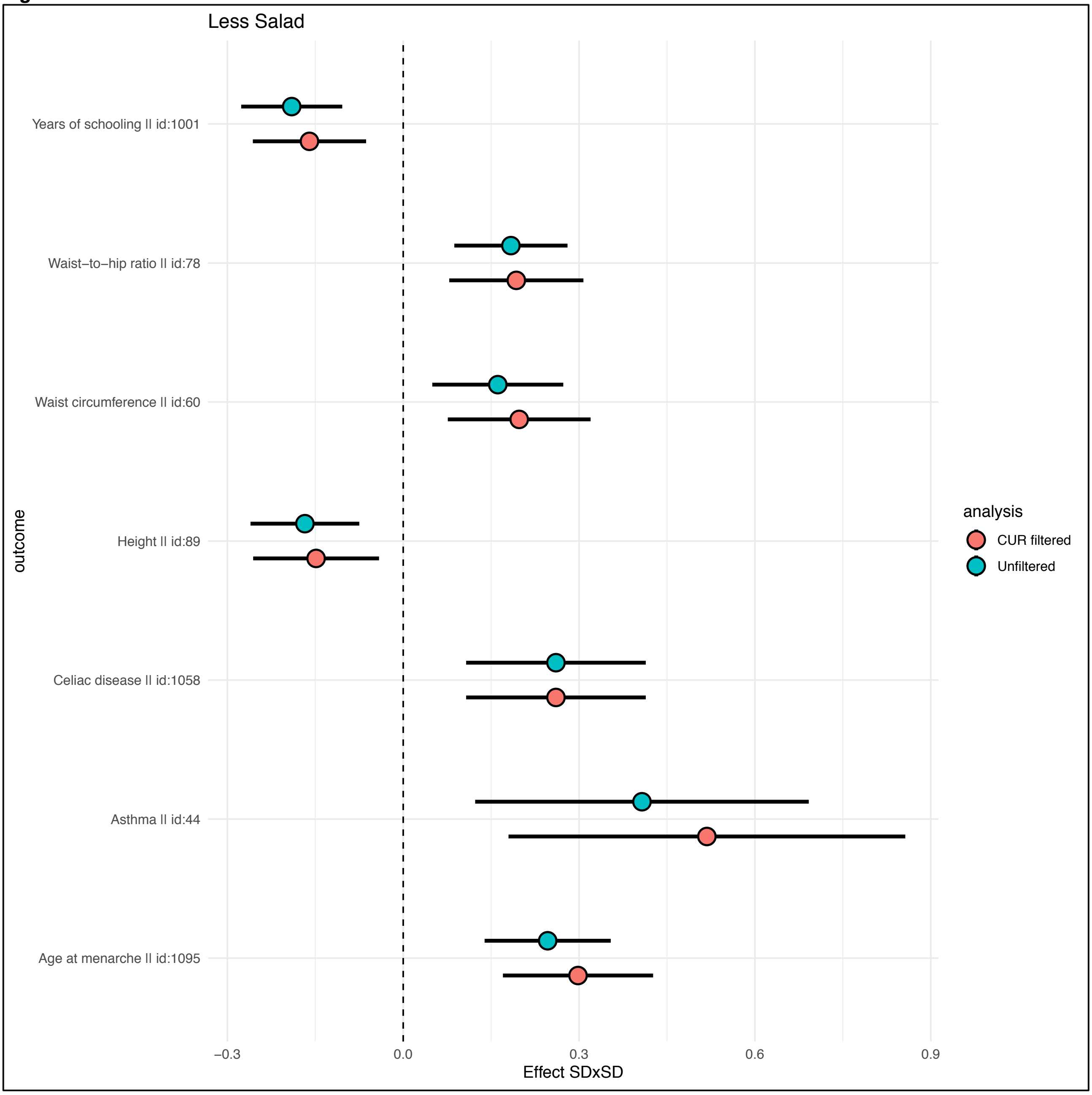


724

***Fig S40***

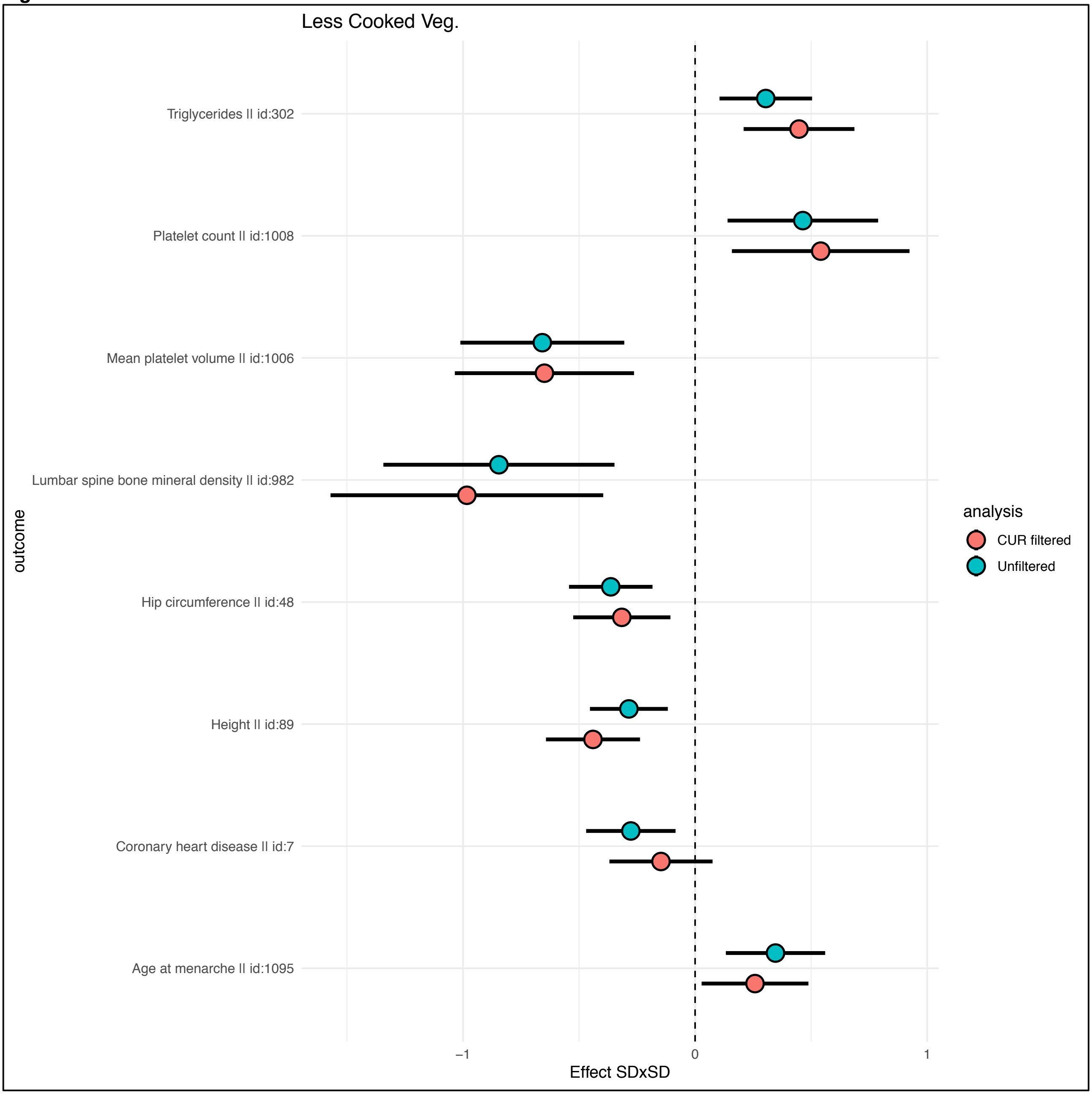
Less Cooked Veg.
Triglycerides II id:302
Platelet count II id:1008
Mean platelet volume II id:1006
Lumbar spine bone mineral density II id:982
Hip circumference II id:48
Height II id:89
Coronary heart disease II id:7
Age at menarche II id:1095
outcome
−1
0
1
Effect SDxSD
analysis
CUR filtered
Unfiltered

***Fig S41***

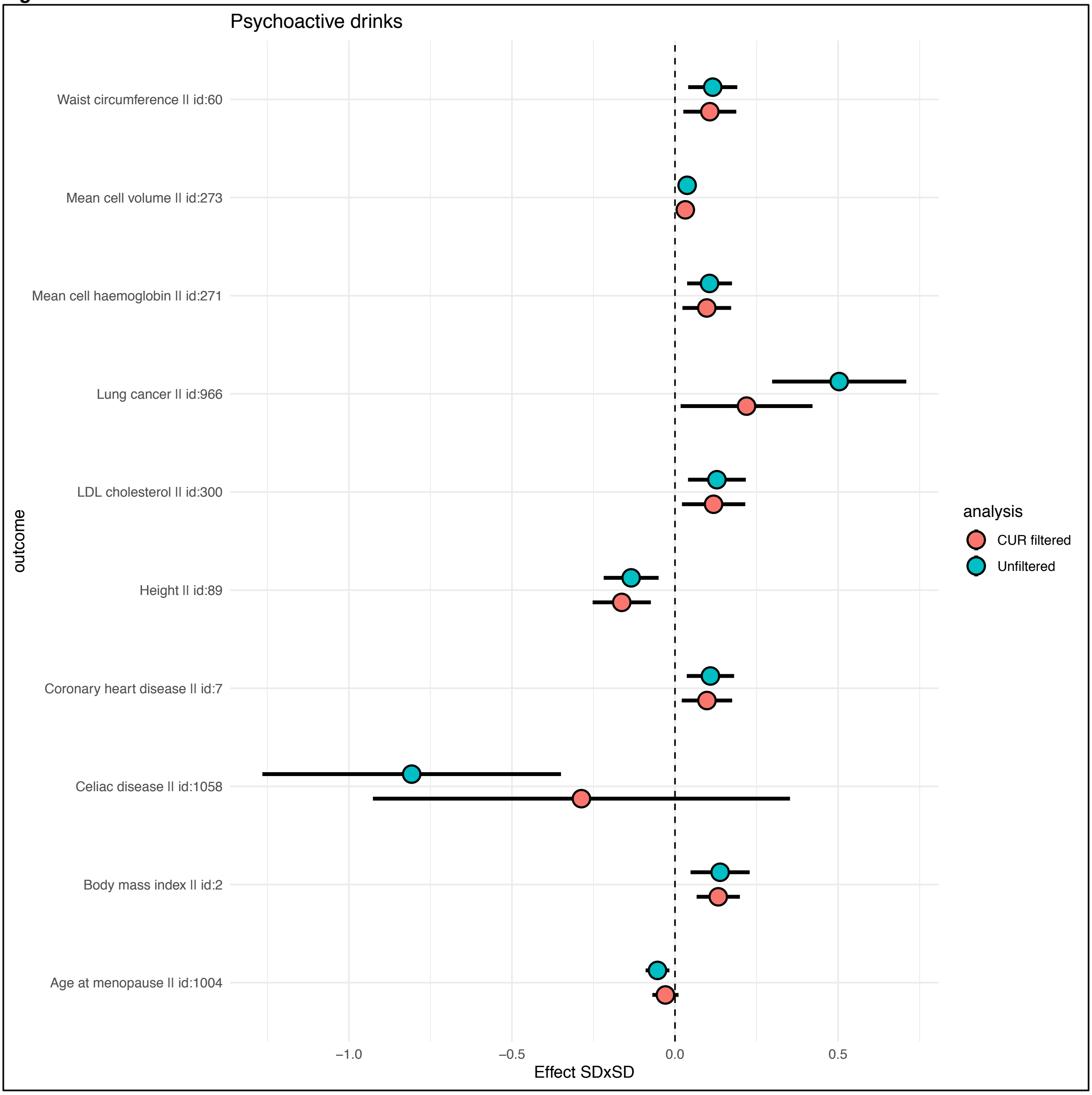

Psychoactive drinks
Waist circumference || id:60
Mean cell volume || id:273
Mean cell haemoglobin || id:271
Lung cancer || id:966
LDL cholesterol || id:300
Height || id:89
Coronary heart disease || id:7
Celiac disease || id:1058
Body mass index || id:2
Age at menopause || id:1004
outcome
−1.0
−0.5
0.0
0.5
Effect SDxSD
analysis
CUR filtered
Unfiltered

***Fig S42***

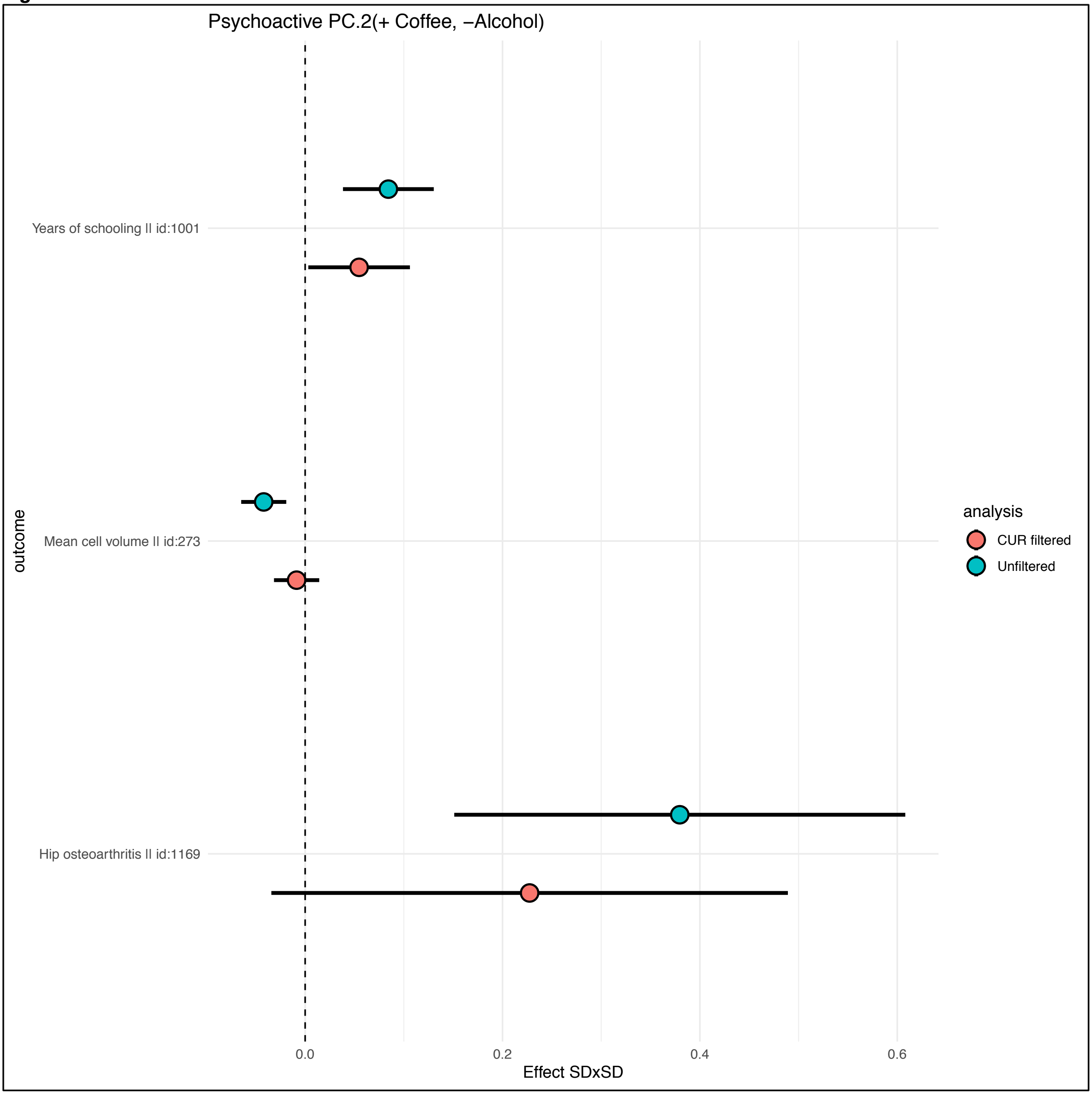

Psychoactive PC.2(+ Coffee, −Alcohol)
Years of schooling || id:1001
Mean cell volume || id:273
Hip osteoarthritis || id:1169
outcome
0.0
0.2
0.4
0.6
Effect SDxSD
analysis
CUR filtered
Unfiltered

***Fig S43***

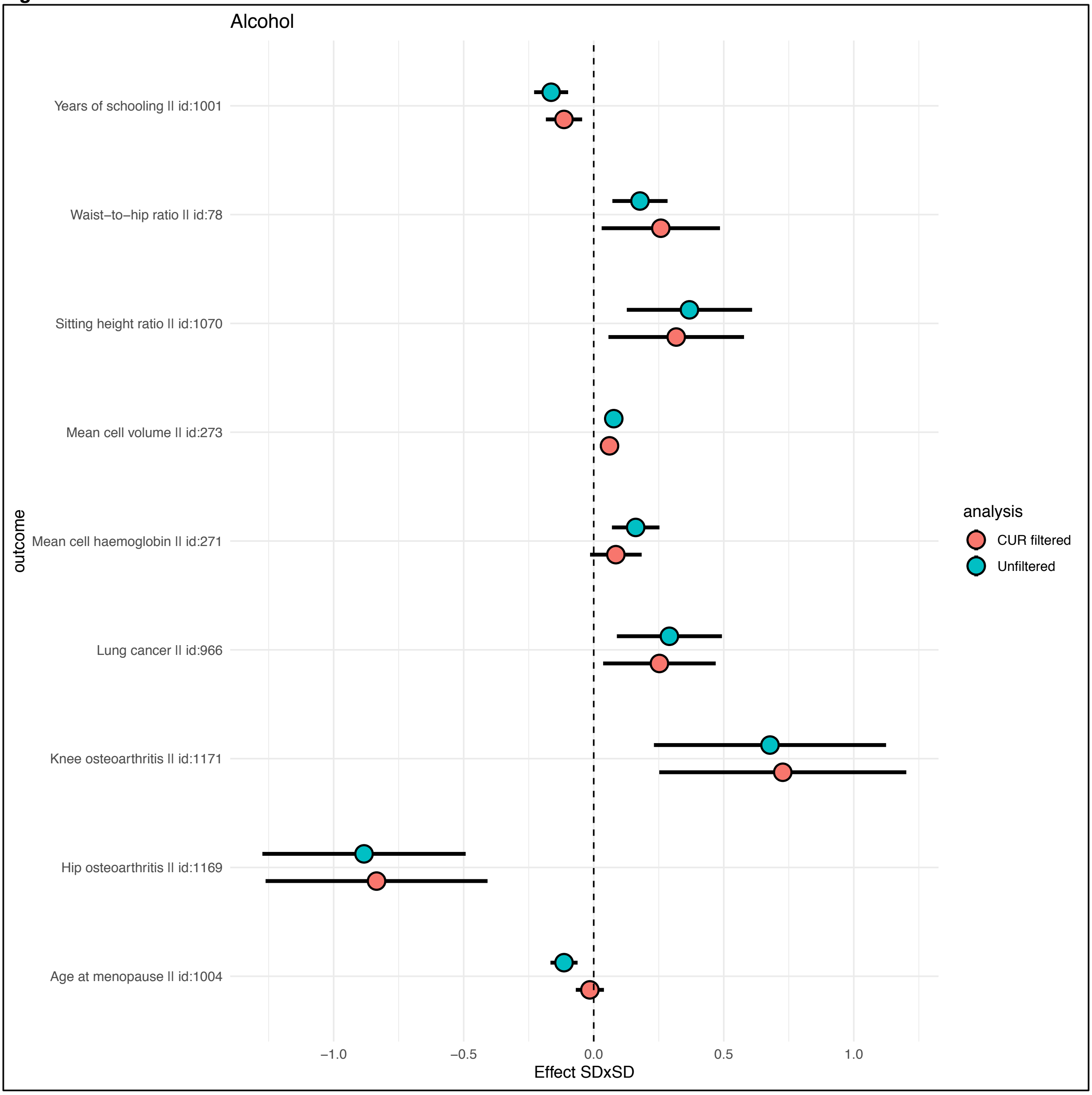
Alcohol
Years of schooling || id:1001
Waist−to−hip ratio || id:78
Sitting height ratio || id:1070
Mean cell volume || id:273
Mean cell haemoglobin || id:271
Lung cancer || id:966
Knee osteoarthritis || id:1171
Hip osteoarthritis || id:1169
Age at menopause || id:1004
outcome
−1.0
−0.5
0.0
0.5
1.0
Effect SDxSD
analysis
CUR filtered
Unfiltered

733 ***Fig S44***

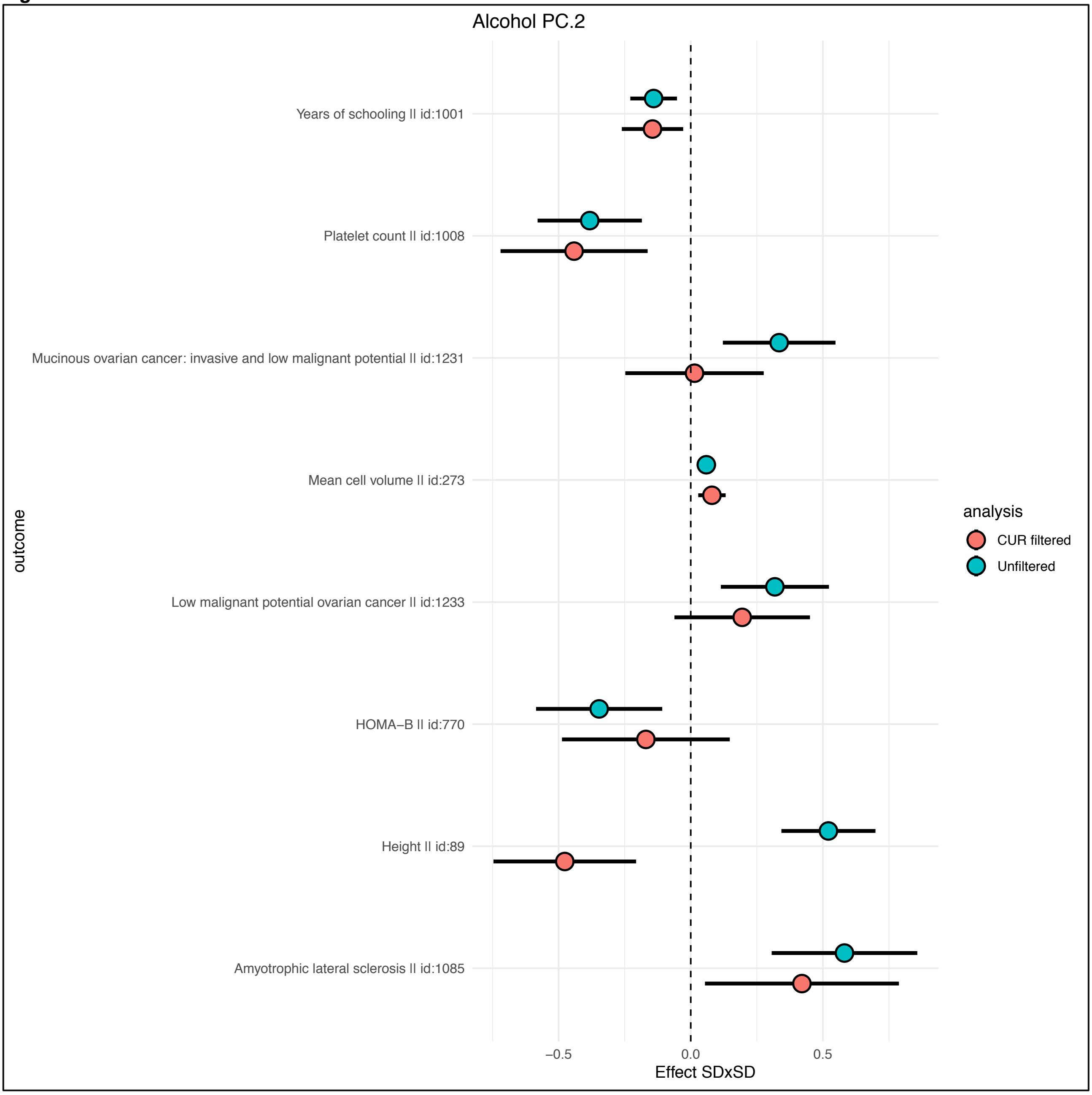


734

735 *Fig S45*

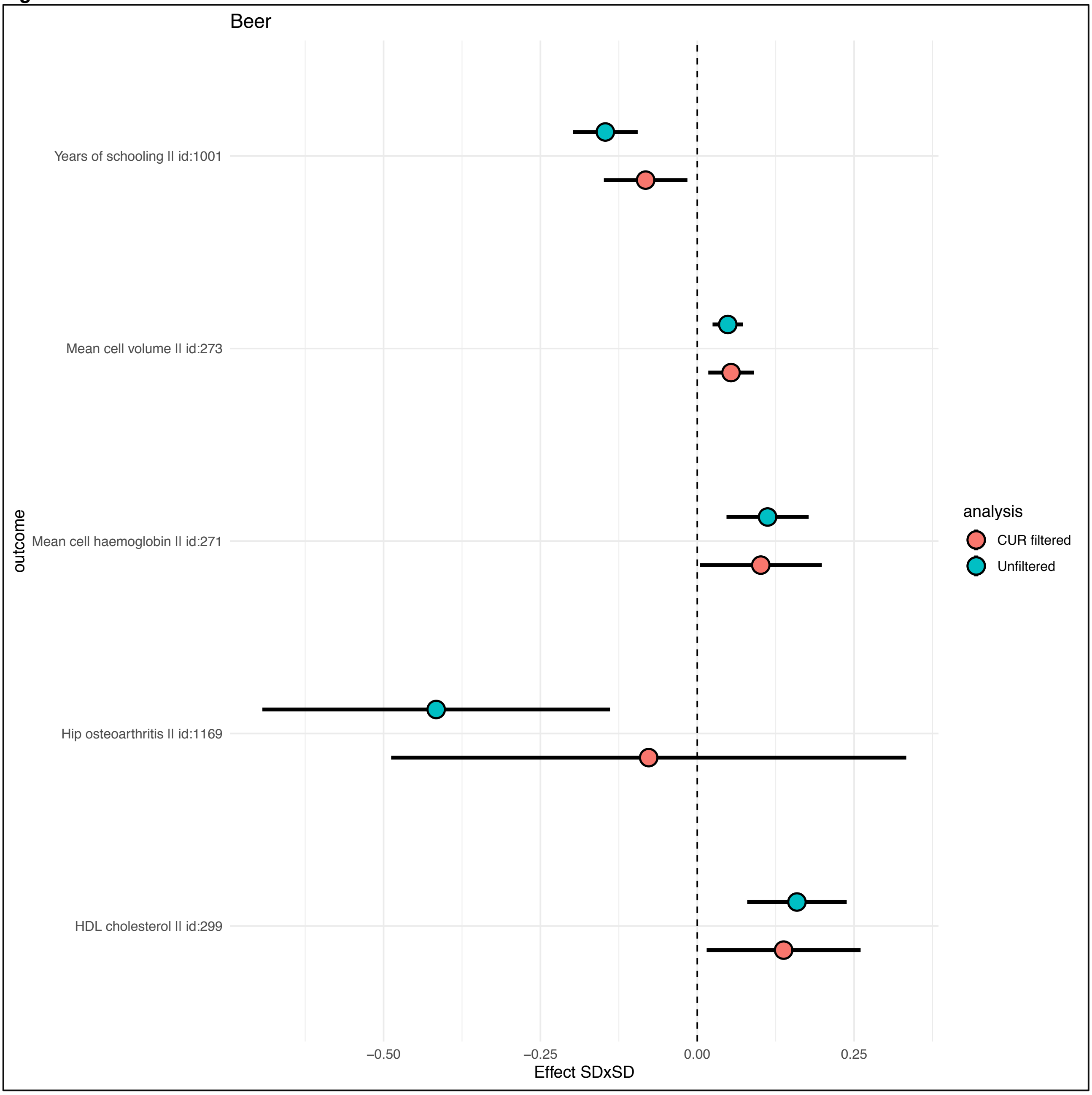


736

***Fig S46***

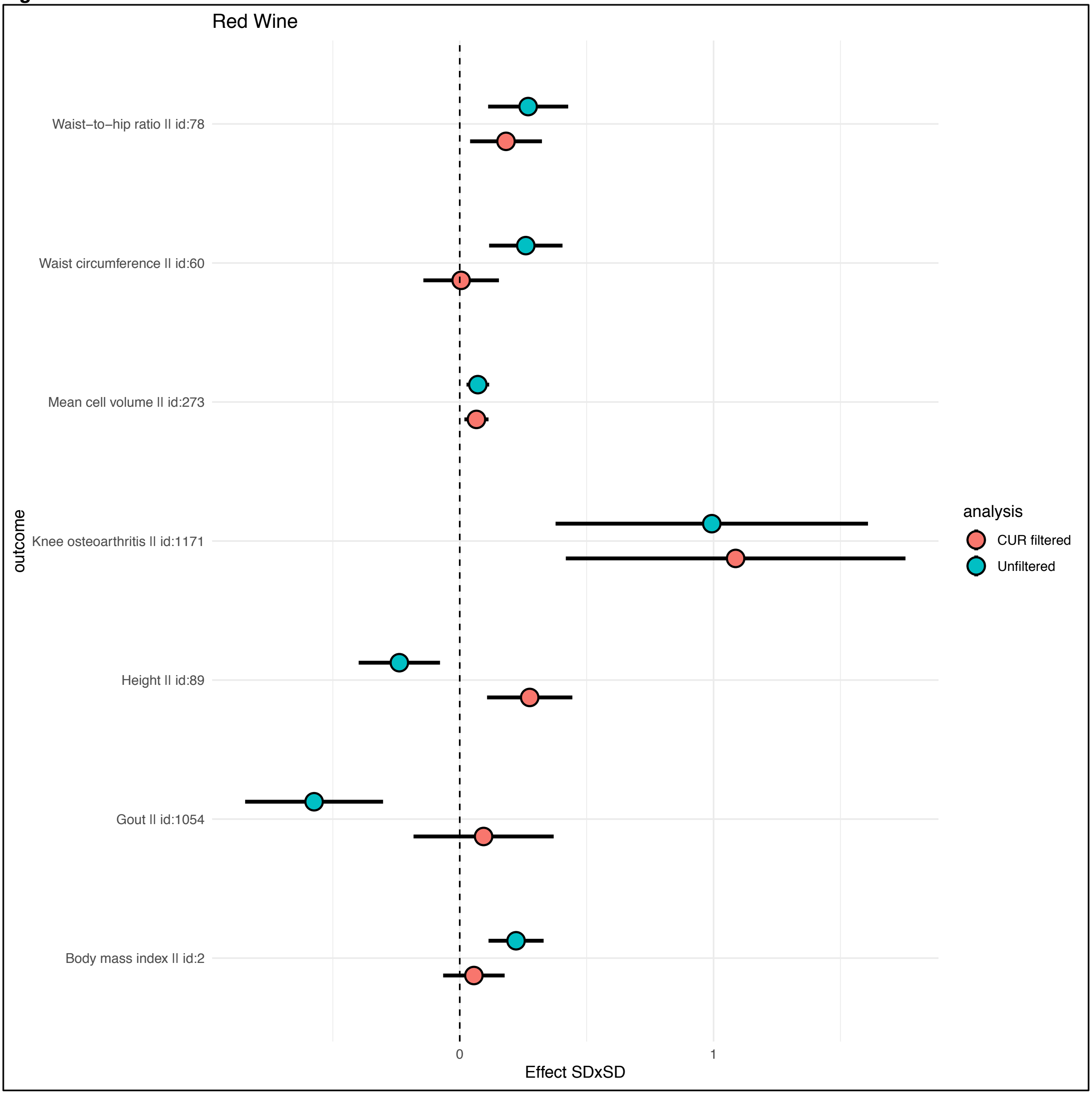
Red Wine
Waist−to−hip ratio || id:78
Waist circumference || id:60
Mean cell volume || id:273
Knee osteoarthritis || id:1171
Height || id:89
Gout || id:1054
Body mass index || id:2
outcome
0
1
Effect SDxSD
analysis
CUR filtered
Unfiltered

***Fig S47***

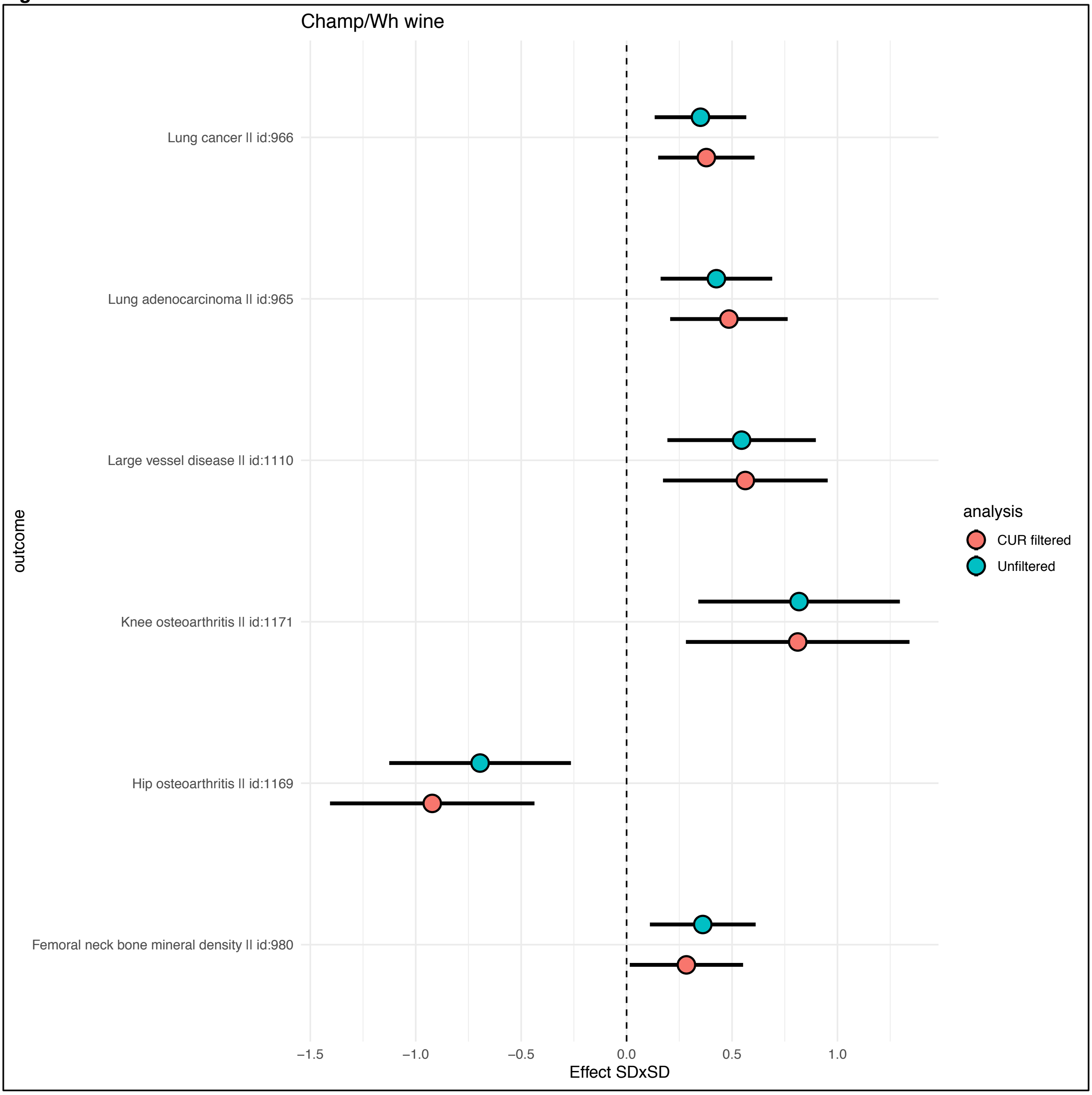
Champ/Wh wine
Lung cancer || id:966
Lung adenocarcinoma || id:965
Large vessel disease || id:1110
Knee osteoarthritis || id:1171
Hip osteoarthritis || id:1169
Femoral neck bone mineral density || id:980
outcome
−1.5
−1.0
−0.5
0.0
0.5
1.0
Effect SDxSD
analysis
CUR filtered
Unfiltered

***Fig S48***

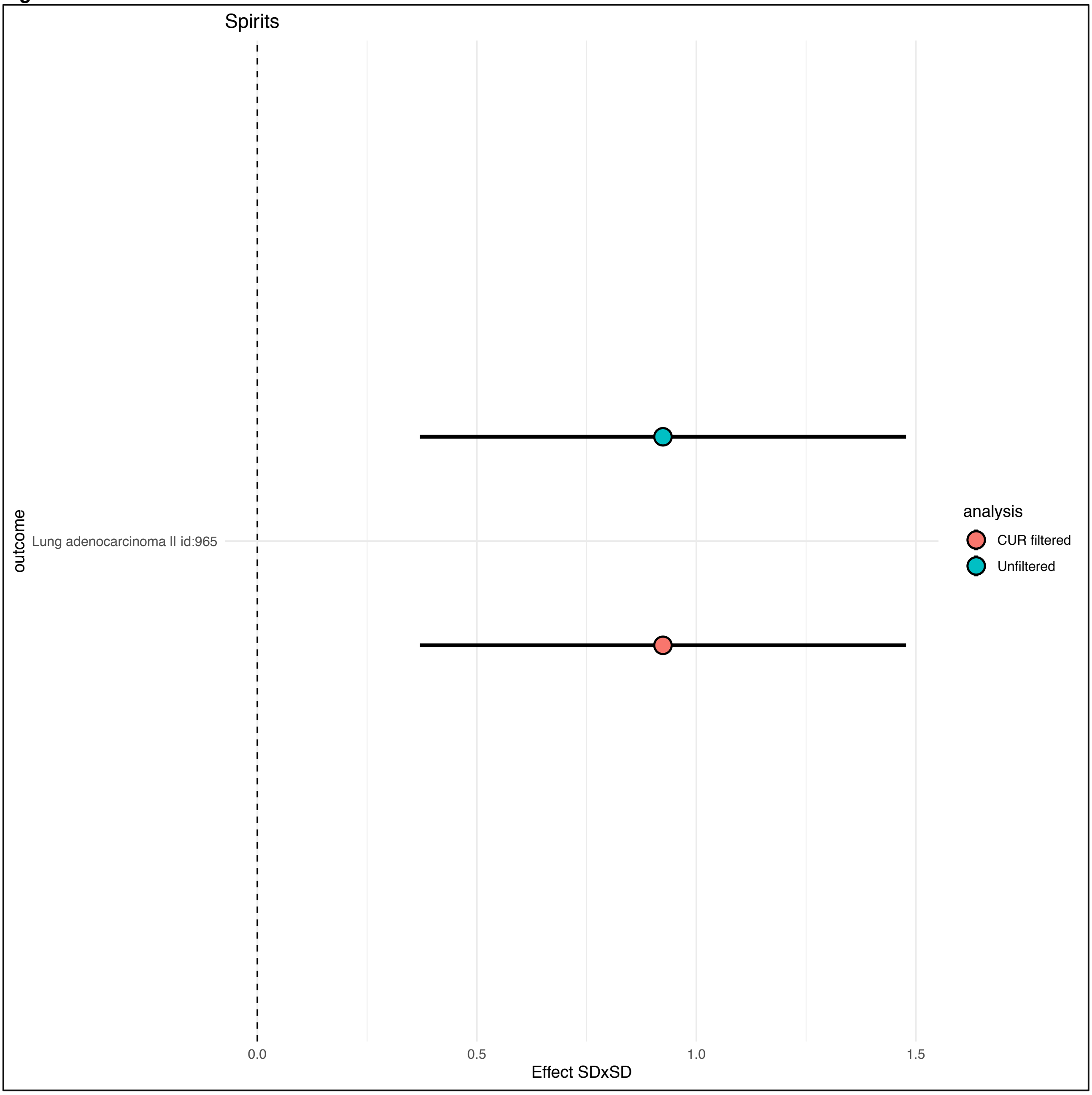
Spirits
outcome
Lung adenocarcinoma II id:965
0.0
0.5
1.0
1.5
Effect SDxSD
analysis
CUR filtered
Unfiltered

***Fig S49***

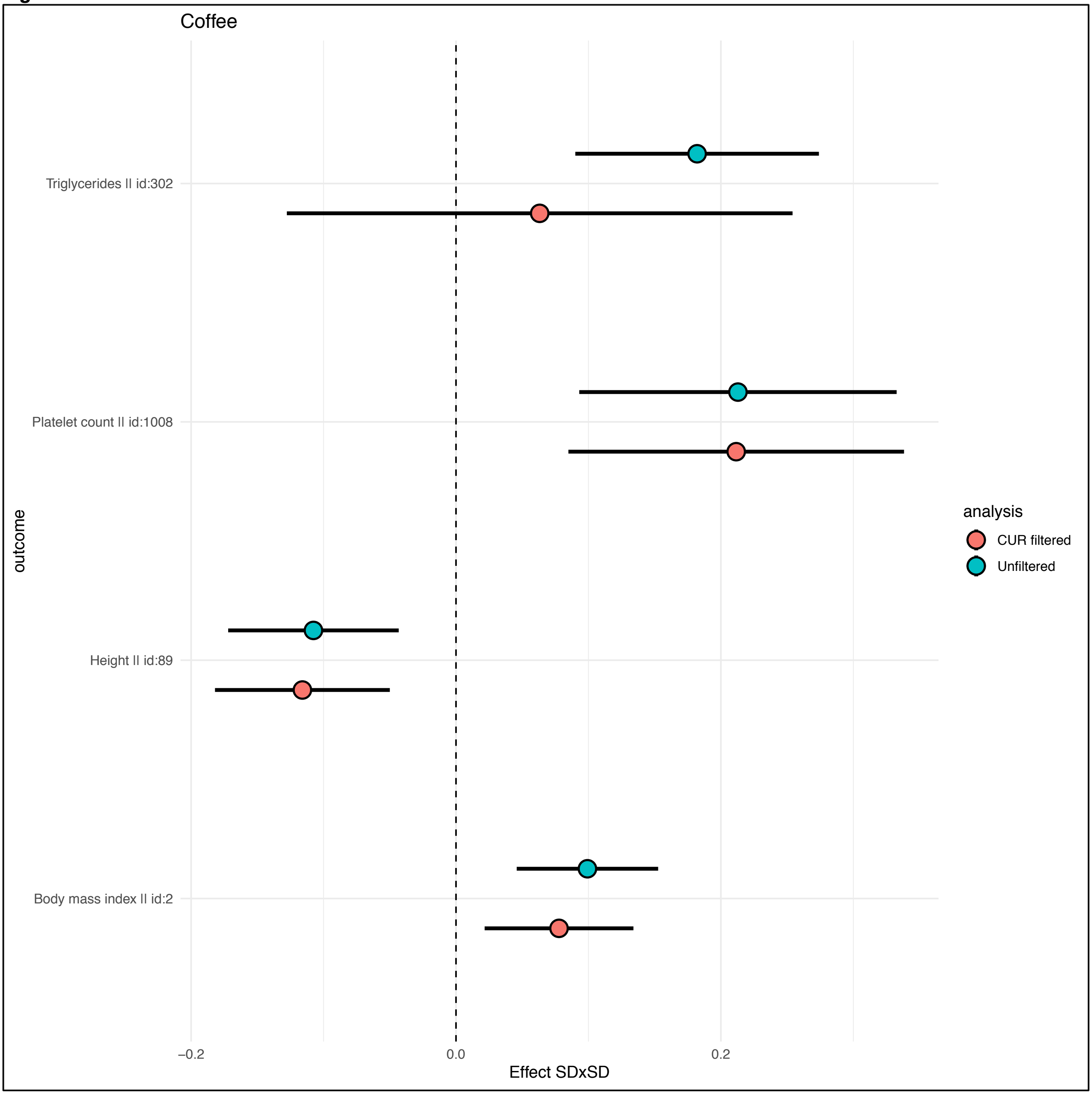

Coffee
Triglycerides II id:302
Platelet count II id:1008
Height II id:89
Body mass index II id:2
outcome
analysis
CUR filtered
Unfiltered
−0.2
0.0
0.2
Effect SDxSD

***Fig S50***

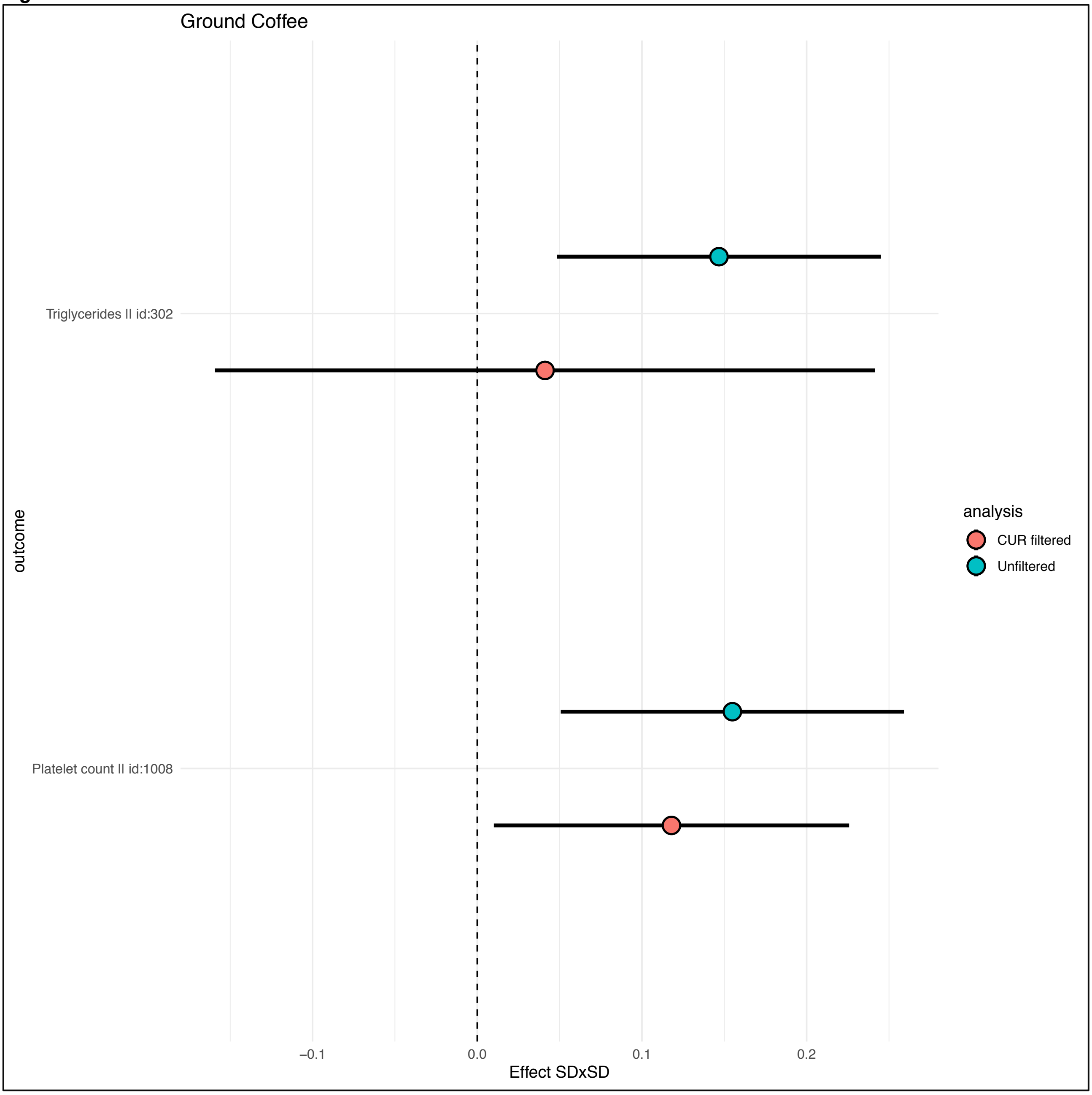
Ground Coffee
Triglycerides II id:302
Platelet count II id:1008
outcome
−0.1
0.0
0.1
0.2
Effect SDxSD
analysis
CUR filtered
Unfiltered

748 ***Fig S51***

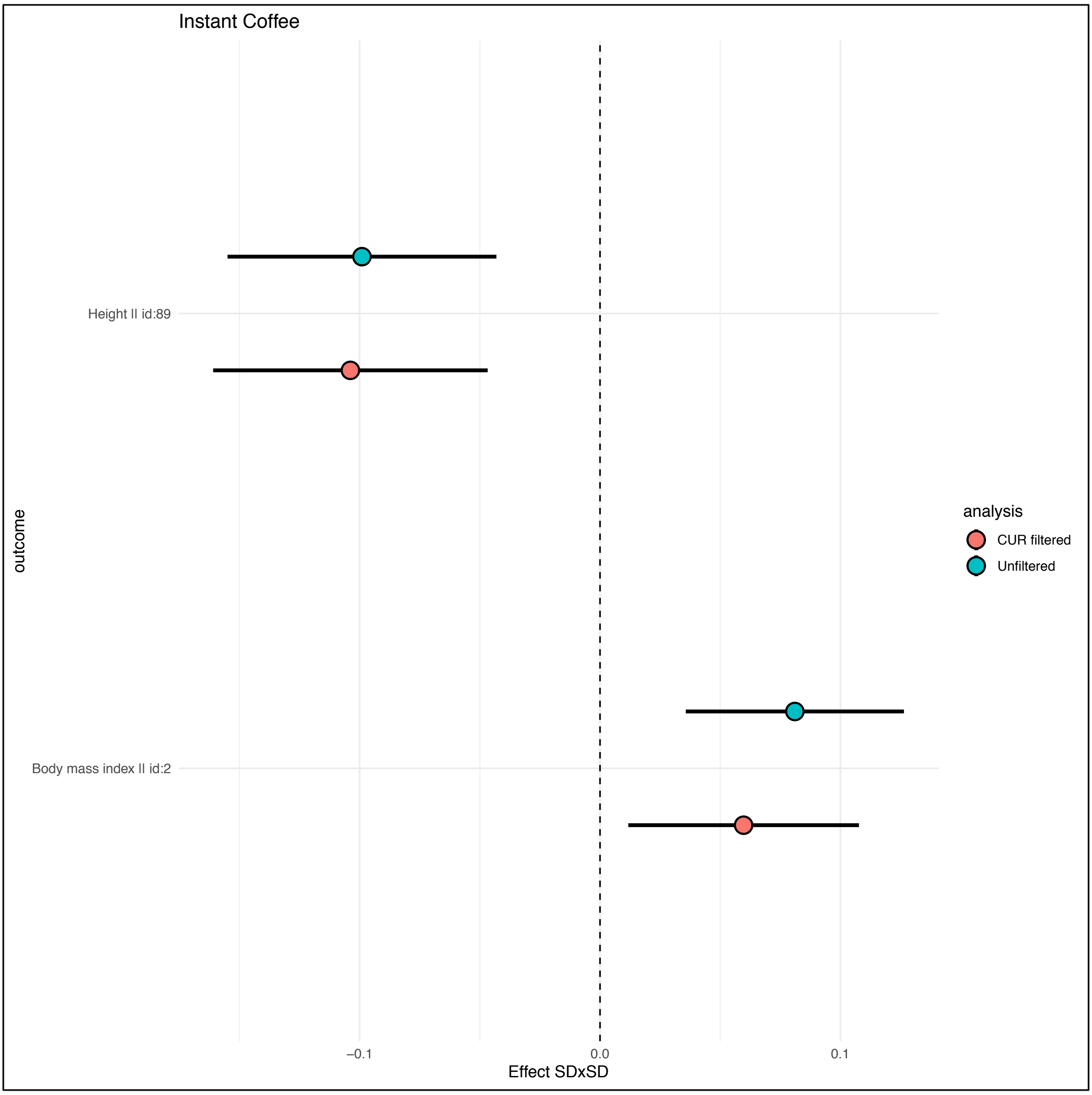


749

***Fig S52***

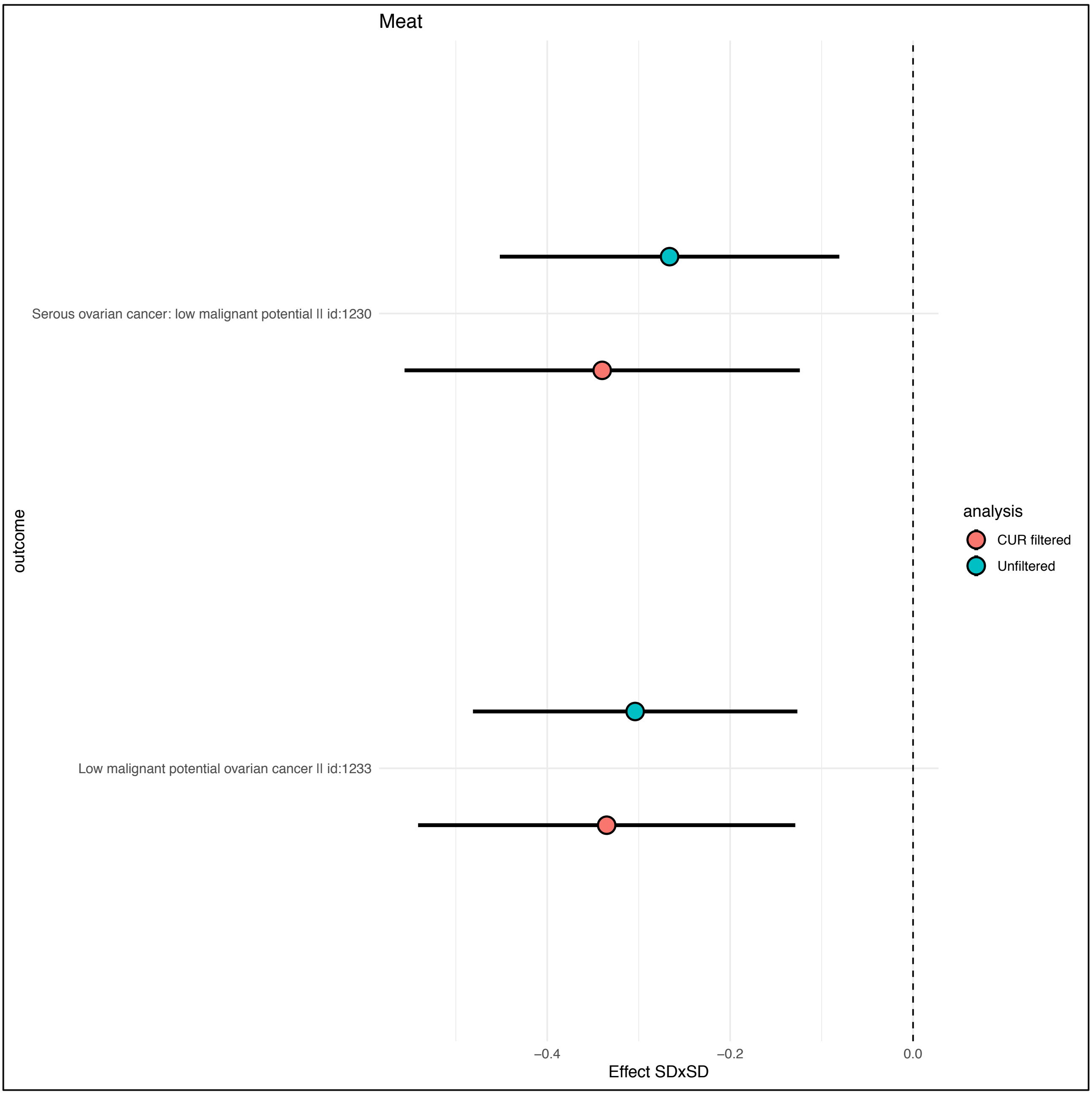
Meat
Serous ovarian cancer: low malignant potential II id:1230
Low malignant potential ovarian cancer II id:1233
outcome
−0.4
−0.2
0.0
Effect SDxSD
analysis
CUR filtered
Unfiltered

***Fig S53***

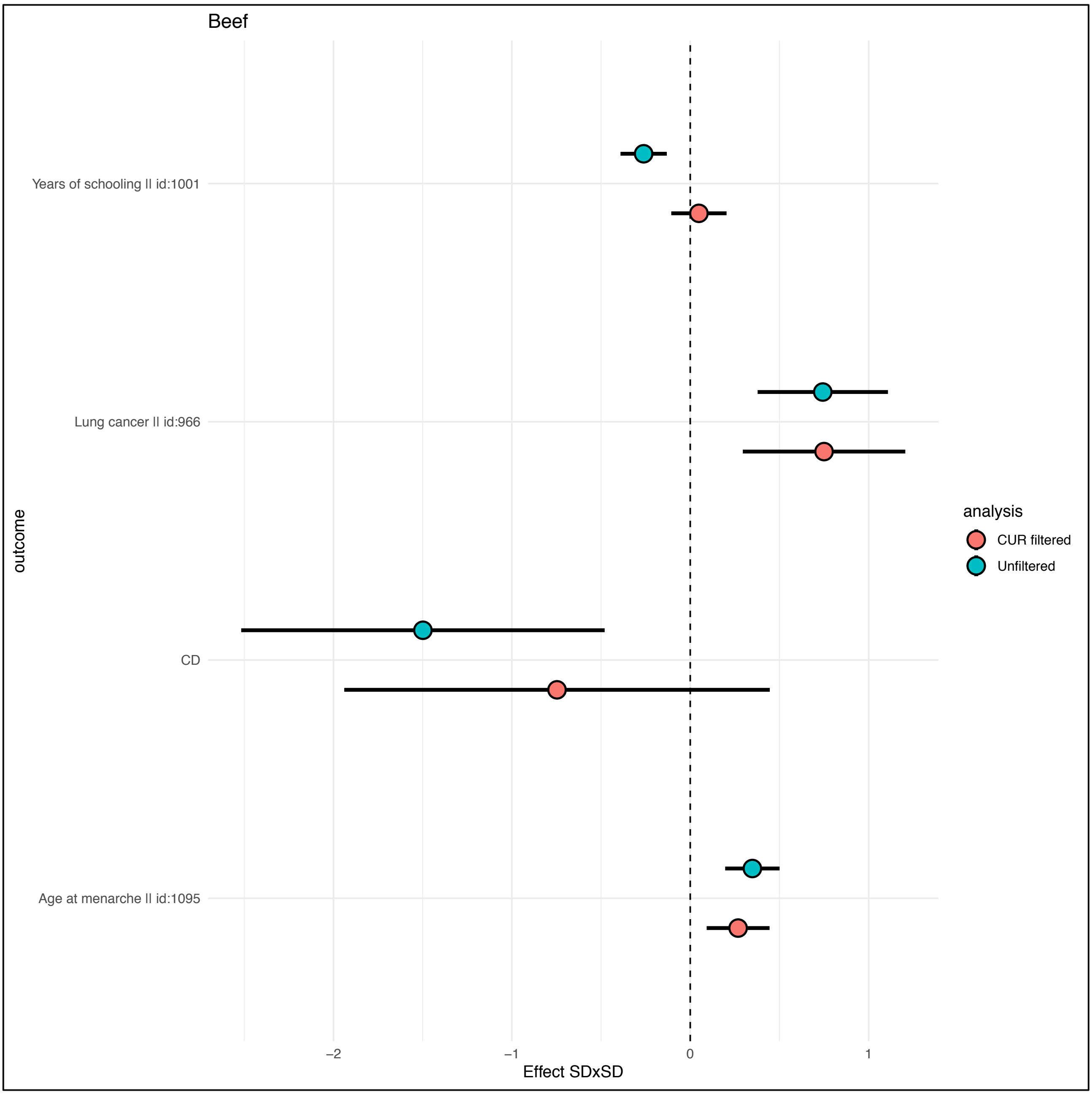

Beef
Years of schooling || id:1001
Lung cancer || id:966
CD
Age at menarche || id:1095
outcome
−2
−1
0
1
Effect SDxSD
analysis
CUR filtered
Unfiltered

***Fig S54***

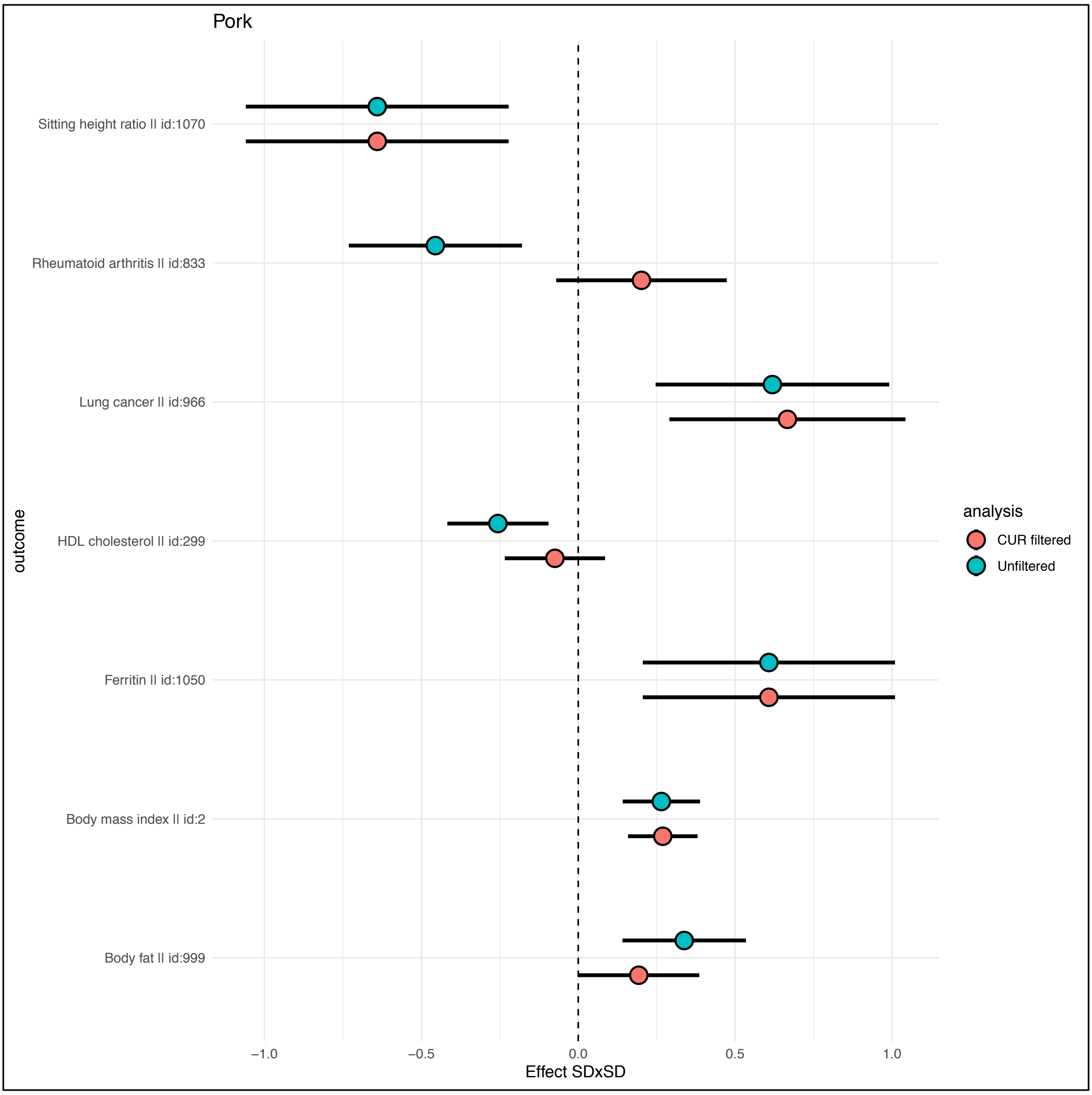
Pork
Sitting height ratio II id:1070
Rheumatoid arthritis II id:833
Lung cancer II id:966
HDL cholesterol II id:299
Ferritin II id:1050
Body mass index II id:2
Body fat II id:999
outcome
−1.0
−0.5
0.0
0.5
1.0
Effect SDxSD
analysis
CUR filtered
Unfiltered

756 ***Fig S55***

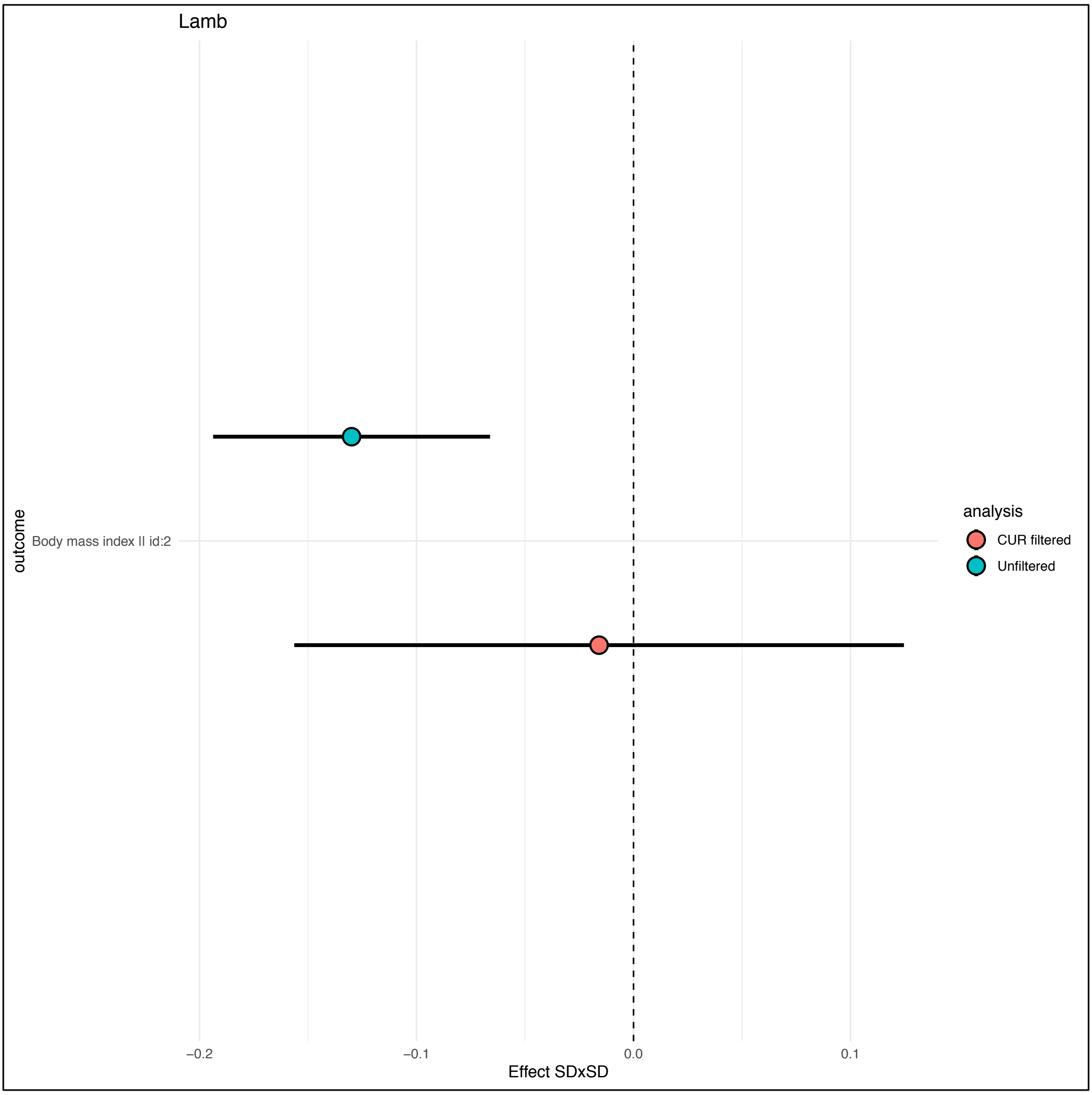


757

*Fig S56*

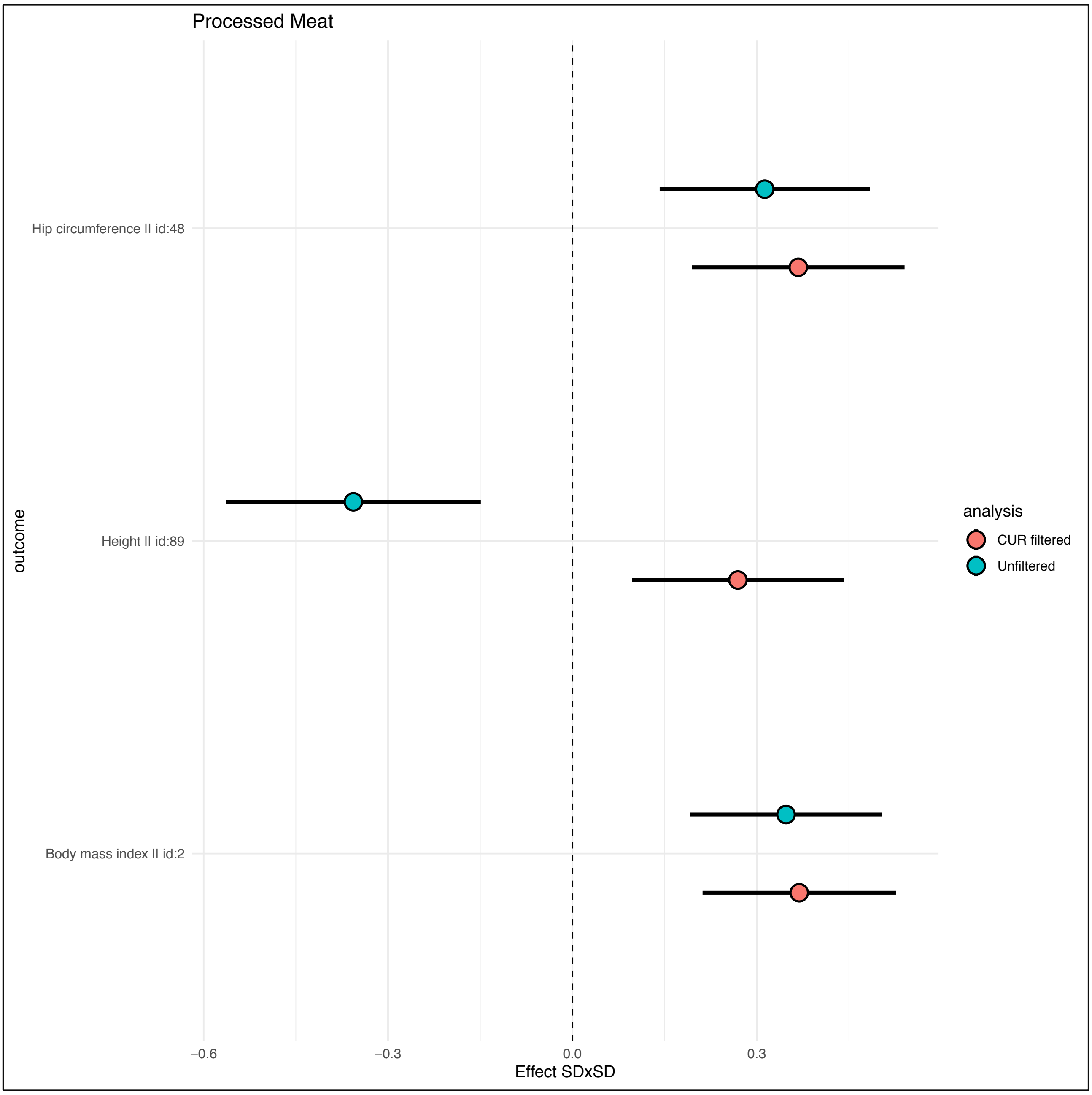

Processed Meat
Hip circumference || id:48
Height || id:89
Body mass index || id:2
outcome
−0.6
−0.3
0.0
0.3
Effect SDxSD
analysis
CUR filtered
Unfiltered

***Fig S57***

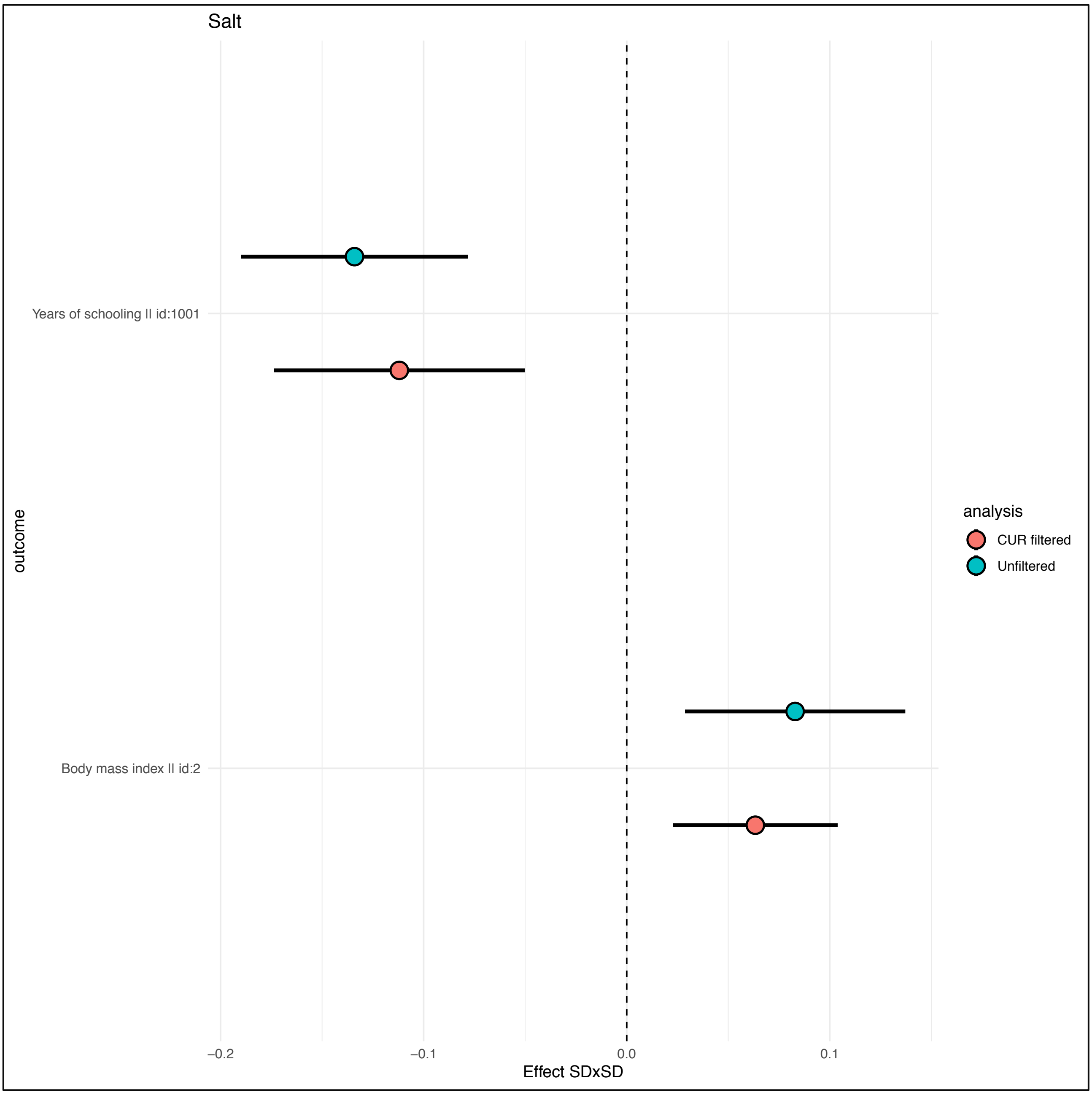
Salt
Years of schooling || id:1001
Body mass index || id:2
outcome
−0.2
−0.1
0.0
0.1
Effect SDxSD
analysis
CUR filtered
Unfiltered

*Fig S58*

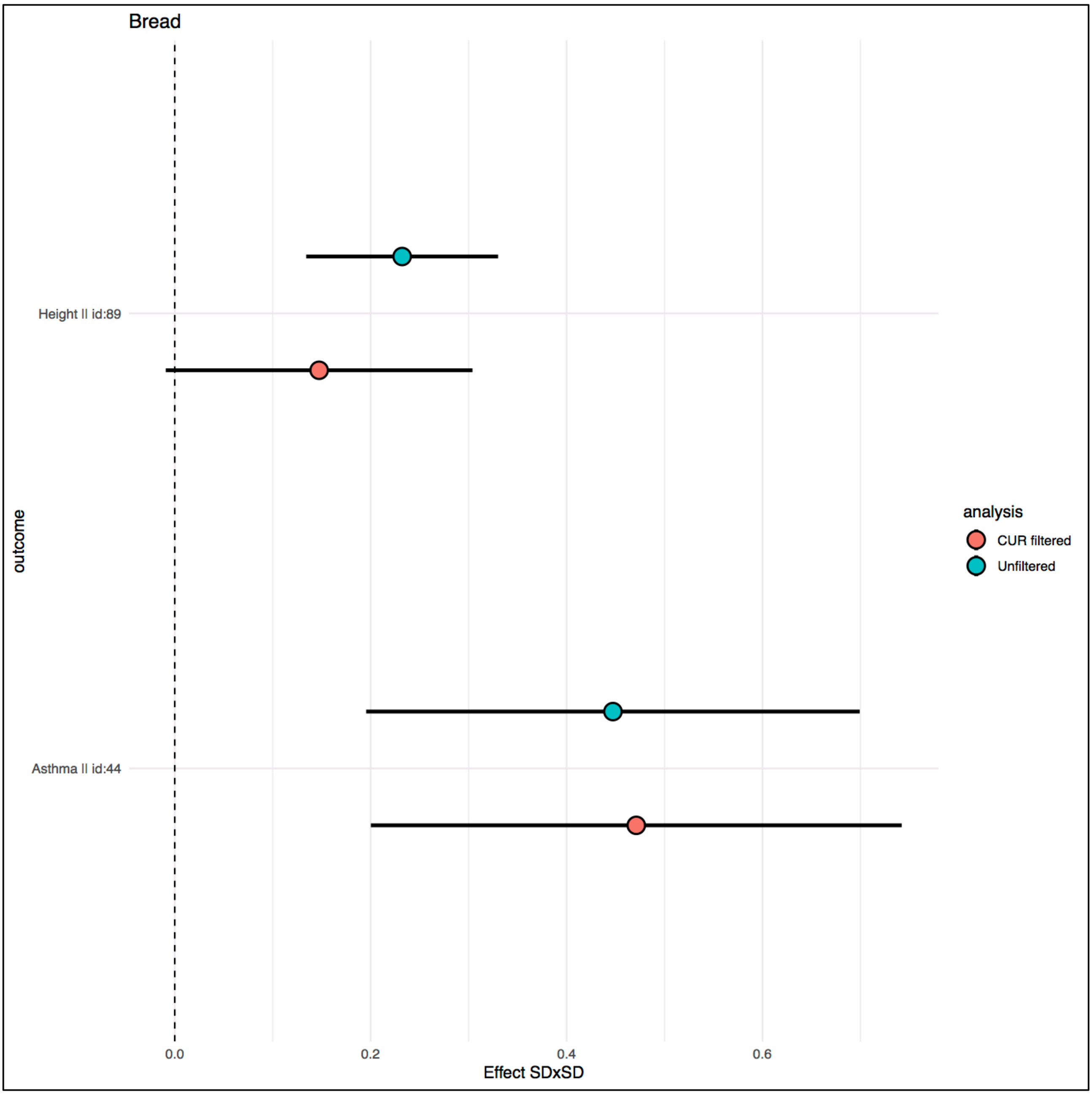
Bread
Height || id:89
Asthma || id:44
outcome
0.0
0.2
0.4
0.6
Effect SDxSD
analysis
CUR filtered
Unfiltered

765 ***Fig S59***

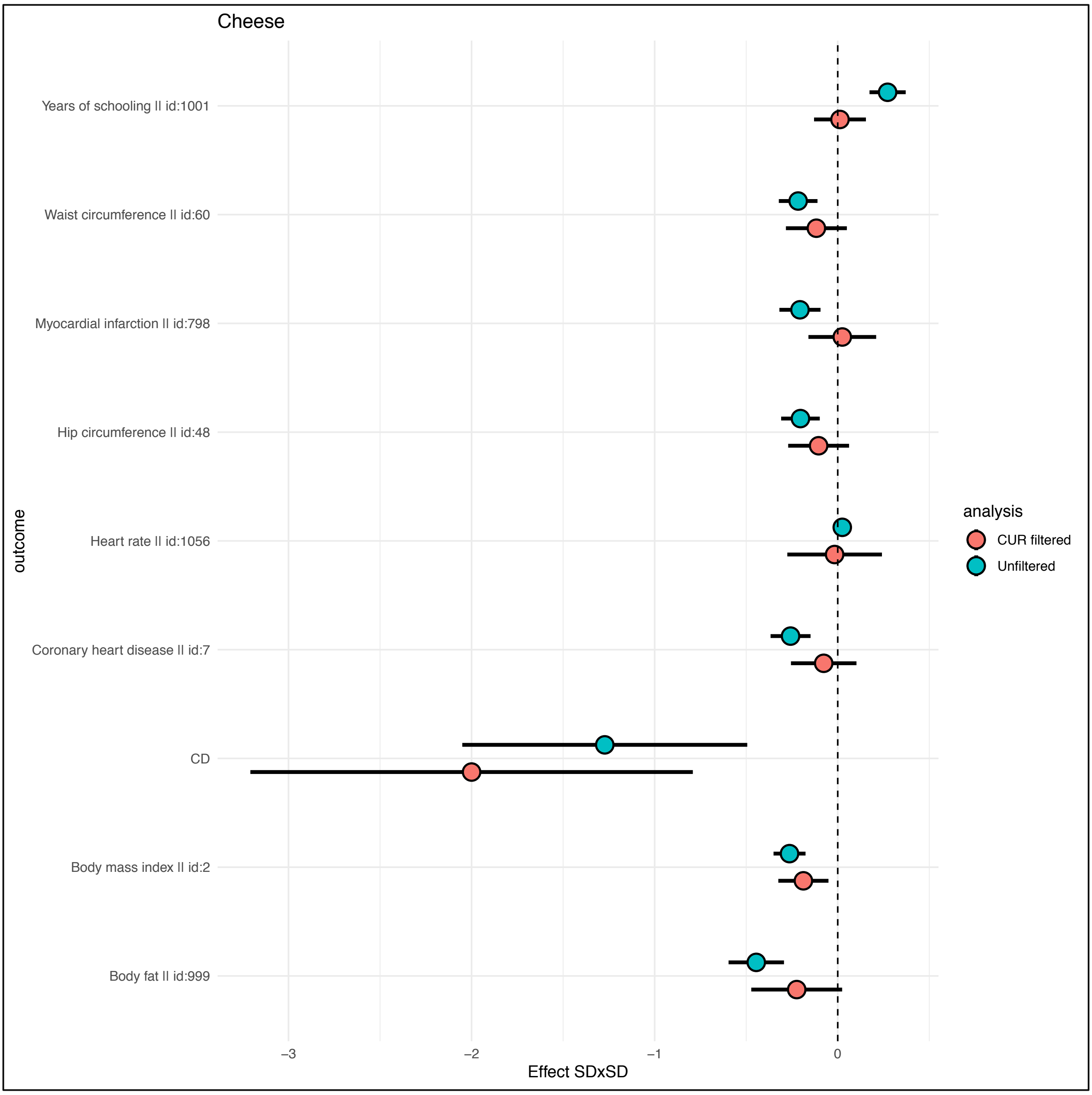


766

***Fig S60***

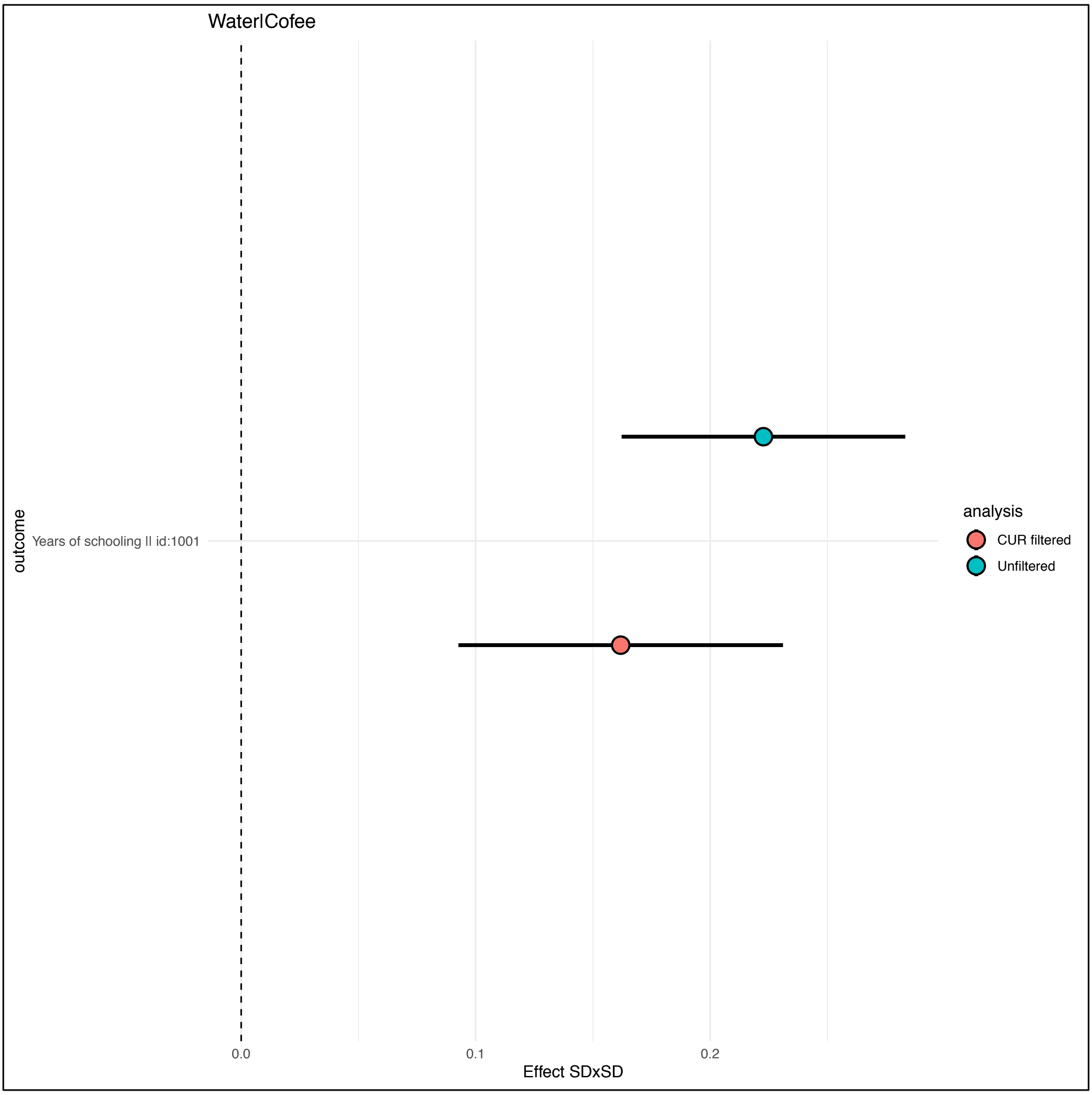


**2.7 Comparison of the CUR filtering with Mendellian randomization common practice.**

It is common practice in Mendellian randomization studies to determine the effect of horizontal pleiotropy through sensitivity analyses using methods such as the weighted median[26] or by evaluating the amount of heterogeneity in the effect estimates. In our case however we have shown that at least in some cases more than half of the variants can’t be considered reliable. This means on one side that the assumption of the weighted median that at least half of the SNPs are valid IVs is violated for several of our food stuff and on the other that heterogeneity filtering (as in

MR-Radial[27], or MR-PRESSO[28]) could potentially exclude the SNPs which are the true valid instruments. This is extremely problematic as it would lead to the wrong conclusions and results. We have in fact used the MR-Radial method in the MR pipeline and still obtained biased results. Another important point is that for the selection of the IVs it is in principle possible to use Steiger filtering[29] for understanding for each SNP if the causal pathway goes from the exposure to the outcome or the opposite. The test is based on evaluating how much variance of the exposure and the outcome is explained by the instrument, if the SNP explains more variance of the exposure than the outcome we can assume that the causal pathway goes from the exposure to the outcome, if the opposite is true, we can then infer that the causal pathway is going in the other direction. Although the test is generally quite sturdy, in our case the extremely high noise present in the phenotype makes using Steiger's test for distinguishing the SNPs directly associated to the food trait from those associated through other traits less reliable.

To understand this issue let's imagine for simplicity that BMI is the only trait which is influencing causally either food consumption or food frequency questionnaire answers. As we cannot distinguish between the two cases not having an objective measure of consumption we can regard the two as being the same phenomenon, in fact if BMI influences how much we eat this will be transferred to the FFQ accordingly. Given we cannot distinguish between the two we need to evaluate any such effect as having a biasing effect on the FFQ response.

Adding a SNP which is causal to BMI the resulting DAG would look as such

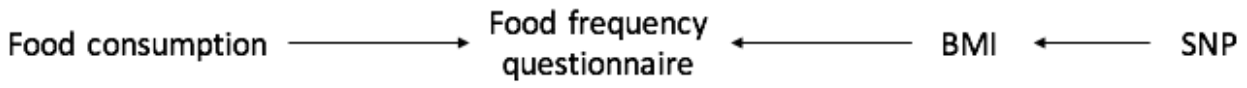


In this case Steiger test will work as the variance explained by the SNP of BMI will always be greater than that explained on the FFQ as the effect is mediated through BMI, this is even truer if the effect is not directly on the FFQ but is mediated through FC.

So if we knew the factors biasing the FFQ and if they were independent from each other we could in principle use Steiger test to distinguish which are the SNPs which are mediated through these factors.

The problem arises when food consumption is causal to the confounding factor (as in many of the traits we used). For example, BMI is clearly caused by Food consumption, so after adding this information the DAG would look like.

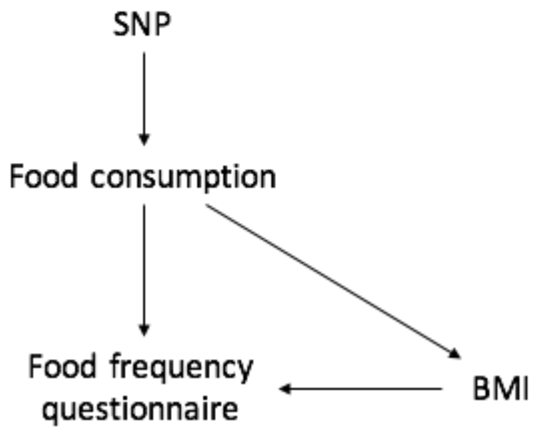


In this case we can think of FFQ as an independent trait which is caused by FC. In this case things get more complicated as the r2 of the SNP on FFQ (which is what we would be using for the comparison) depends on the correlation of FC and FFQ. This can be very variable but it rarely over 0.5[30] with the exception of alcohol and coffee consumption which have higher reliability. Thus, it can be realistic that the causal effect of FC on BMI is similar in size to the direct causal effect of FC on FFQ. In this case, the Steiger test may fail to detect the correct direction of effect. While our approach has similarities to Steiger's test, its aims and settings are quite different in many aspects. First, in our case the causal direction is clear because it is highly unlikely that FFQ items cause other traits, only the upstream FC can do so. Second, our underlying DAG is more complicated (with multiple exposures and underlying FC) than it is assumed by the Steiger test and our approach fully exploits the a priori knowledge of the DAG. On the other hand, the Steiger test could be used to select valid (direct) exposure (e.g. BMI) instruments in order to estimate the total (direct plus indirect) exposure->FFQ causal effect, which in turn could be used to derive direct and indirect SNP-FFQ effects.

#### Bibiography


1. Sudlow, C. *et al.* UK biobank: an open access resource for identifying the causes of a wide range of complex diseases of middle and old age. *PLoS Med.* **12**, e1001779 (2015).
2. Loh, P.-R. *et al.* Efficient Bayesian mixed model analysis increases association power in large cohorts. doi:10.1101/007799.
3. McCarthy, S. *et al.* A reference panel of 64,976 haplotypes for genotype imputation. *Nat. Genet.* **48**, 1279–1283 (2016).
4. Bulik-Sullivan, B. K. *et al.* LD Score regression distinguishes confounding from polygenicity in genome-wide association studies. *Nat. Genet.* **47**, 291–295 (2015).
5. Day, N. *et al.* EPIC-Norfolk: study design and characteristics of the cohort. European Prospective Investigation of Cancer. *Br. J. Cancer* **80 Suppl 1**, 95–103 (1999).
6. Lotta, L. A. *et al.* Integrative genomic analysis implicates limited peripheral adipose storage capacity in the pathogenesis of human insulin resistance. *Nat. Genet.* **49**, 17–26 (2017).
7. Bingham, S. A. *et al.* Comparison of dietary assessment methods in nutritional epidemiology: weighed records v. 24 h recalls, food-frequency questionnaires and estimated-diet records. *Br. J. Nutr.* **72**, 619–643 (1994).
8. Hemani, G. *et al.* The MR-Base platform supports systematic causal inference across the human phenome. *Elife* **7**, (2018).
9. McDaid, A. F. *et al.* Bayesian association scan reveals loci associated with human lifespan and linked biomarkers. *Nat. Commun.* **8**, 15842 (2017).
10. VanderWeele, T. J. A three-way decomposition of a total effect into direct, indirect, and interactive effects. *Epidemiology* **24**, 224–232 (2013).
11. Rüeger, S., McDaid, A. & Kutalik, Z. Evaluation and application of summary statistic imputation to discover new height-associated loci. *PLoS Genet.* **14**, e1007371 (2018).
12. de Leeuw, C. A., Mooij, J. M., Heskes, T. & Posthuma, D. MAGMA: generalized gene-set analysis of GWAS data. *PLoS Comput. Biol.* **11**, e1004219 (2015).

13. Watanabe, K., Taskesen, E., van Bochoven, A. & Posthuma, D. Functional mapping and annotation of genetic associations with FUMA. *Nat. Commun.* **8**, 1826 (2017).

14. Szklarczyk, D. *et al.* STRING v10: protein–protein interaction networks, integrated over the tree of life. *Nucleic Acids Res.* **43**, D447–D452 (2015).

15. Yu, G., Wang, L.-G., Han, Y. & He, Q.-Y. clusterProfiler: an R package for comparing biological themes among gene clusters. *OMICS* **16**, 284–287 (2012).

16. Zheng, J. *et al.* LD Hub: a centralized database and web interface to perform LD score regression that maximizes the potential of summary level GWAS data for SNP heritability and genetic correlation analysis. *Bioinformatics* vol. 33 272–279 (2017).

17. Paul, D. R., Rhodes, D. G., Kramer, M., Baer, D. J. & Rumpler, W. V. Validation of a food frequency questionnaire by direct measurement of habitual ad libitum food intake. *Am. J. Epidemiol.* **162**, 806–814 (2005).

18. Dehghan, M. *et al.* Association of dairy intake with cardiovascular disease and mortality in 21 countries from five continents (PURE): a prospective cohort study. *Lancet* **392**, 2288–2297 (2018).

19. Adam, T. C. & Epel, E. S. Stress, eating and the reward system. *Physiology & Behavior* vol. 91 449–458 (2007).

20. Finucane, H. K. *et al.* Partitioning heritability by functional category using GWAS summary statistics. doi:10.1101/014241.

21. Consortium, G. & GTEx Consortium. Genetic effects on gene expression across human tissues. *Nature* vol. 550 204–213 (2017).

22. Finucane, H. K. *et al.* Heritability enrichment of specifically expressed genes identifies disease-relevant tissues and cell types. *Nat. Genet.* **50**, 621–629 (2018).

23. Bernstein, B. E. *et al.* The NIH Roadmap Epigenomics Mapping Consortium. *Nat. Biotechnol.* **28**, 1045–1048 (2010).

24. Szklarczyk, D. *et al.* STRING v11: protein-protein association networks with increased coverage, supporting functional discovery in genome-wide experimental datasets. *Nucleic Acids Res.* **47**, D607–D613 (2019).

25. Blondel, V. D., Guillaume, J.-L., Lambiotte, R. & Lefebvre, E. Fast unfolding of communities in large networks. *Journal of Statistical Mechanics: Theory and Experiment* vol. 2008 P10008 (2008).

26. Bowden, J., Davey Smith, G., Haycock, P. C. & Burgess, S. Consistent Estimation in Mendelian Randomization with Some Invalid Instruments Using a Weighted Median Estimator. *Genet. Epidemiol.* **40**, 304–314 (2016).

27. Bowden, J. *et al.* Improving the visualisation, interpretation and analysis of two-sample summary data Mendelian randomization via the radial plot and radial regression. doi:10.1101/200378.

28. Verbanck, M., Chen, C.-Y., Neale, B. & Do, R. Publisher Correction: Detection of widespread horizontal pleiotropy in causal relationships inferred from Mendelian randomization between complex traits and diseases. *Nat. Genet.* **50**, 1196 (2018).

29. Hemani, G., Tilling, K. & Davey Smith, G. Orienting the causal relationship between imprecisely measured traits using GWAS summary data. *PLoS Genet.* **13**, e1007081 (2017).

30. Kristal, A. R., Peters, U. & Potter, J. D. Is it time to abandon the food frequency questionnaire? *Cancer Epidemiol. Biomarkers Prev.* **14**, 2826–2828 (2005).
